# Island songbirds as windows into evolution in small populations

**DOI:** 10.1101/2020.04.07.030155

**Authors:** Thibault Leroy, Marjolaine Rousselle, Marie-Ka Tilak, Aude E. Caizergues, Céline Scornavacca, María Recuerda, Jérôme Fuchs, Juan Carlos Illera, Dawie H. De Swardt, Guillermo Blanco, Christophe Thébaud, Borja Milá, Benoit Nabholz

**Author notes:** **Corresponding authors:** Thibault Leroy, Benoit Nabholz.

## Abstract

Due to their limited ranges and inherent isolation, island species have long been recognized as crucial systems for tackling a range of evolutionary questions, including in the early study of speciation [1,2]. Such species have been less studied in the understanding of the evolutionary forces driving DNA sequence evolution. Island species usually have lower census population sizes (*N*) than continental species and, supposedly, lower effective population sizes (*Ne*). Given that both the rates of change caused by genetic drift and by selection are dependent upon *Ne*, island species are theoretically expected to exhibit (i) lower genetic diversity, (ii) less effective natural selection against slightly deleterious mutations [3,4], and (iii) a lower rate of adaptive evolution [5–8, see also Note S1]. Here, we have used a large set of newly sequenced and published whole genome sequences of Passerida bird species or subspecies (14 insular and 11 continental) to test these predictions. We empirically confirm that island species exhibit lower census size and *Ne*, supporting the hypothesis that the smaller area available on islands constrains the upper bound of *Ne*. In the insular species, we find significantly lower nucleotide diversity in coding regions, higher ratios of non-synonymous to synonymous polymorphisms, and lower adaptive substitution rates. Our results provide robust evidence that the lower *Ne* experienced by island species has affected both the ability of natural selection to efficiently remove weakly deleterious mutations and also the adaptive potential of island species, therefore providing considerable empirical support for the nearly neutral theory. We discuss the implications for both evolutionary and conservation biology.

## Results

To assemble our dataset, we used population-level sequencing data from 25 passerine bird species or subspecies, consisting of 14 insular and 11 continental, with a total of 295 individual whole-genome sequences. All species belong to the Passerida lineage, a species-rich clade of songbirds with fairly similar life-history traits(Note S1 for details). Our dataset includes at least 4 independent continental-island transitions that occurred across the songbird phylogeny (Fig. S1) enabling us to efficiently account for phylogenetic structure in all statistical tests reported below (Phylogenetic Generalized Least Square (PGLS), see also Table S1 for additional tests).

**Table 1:**
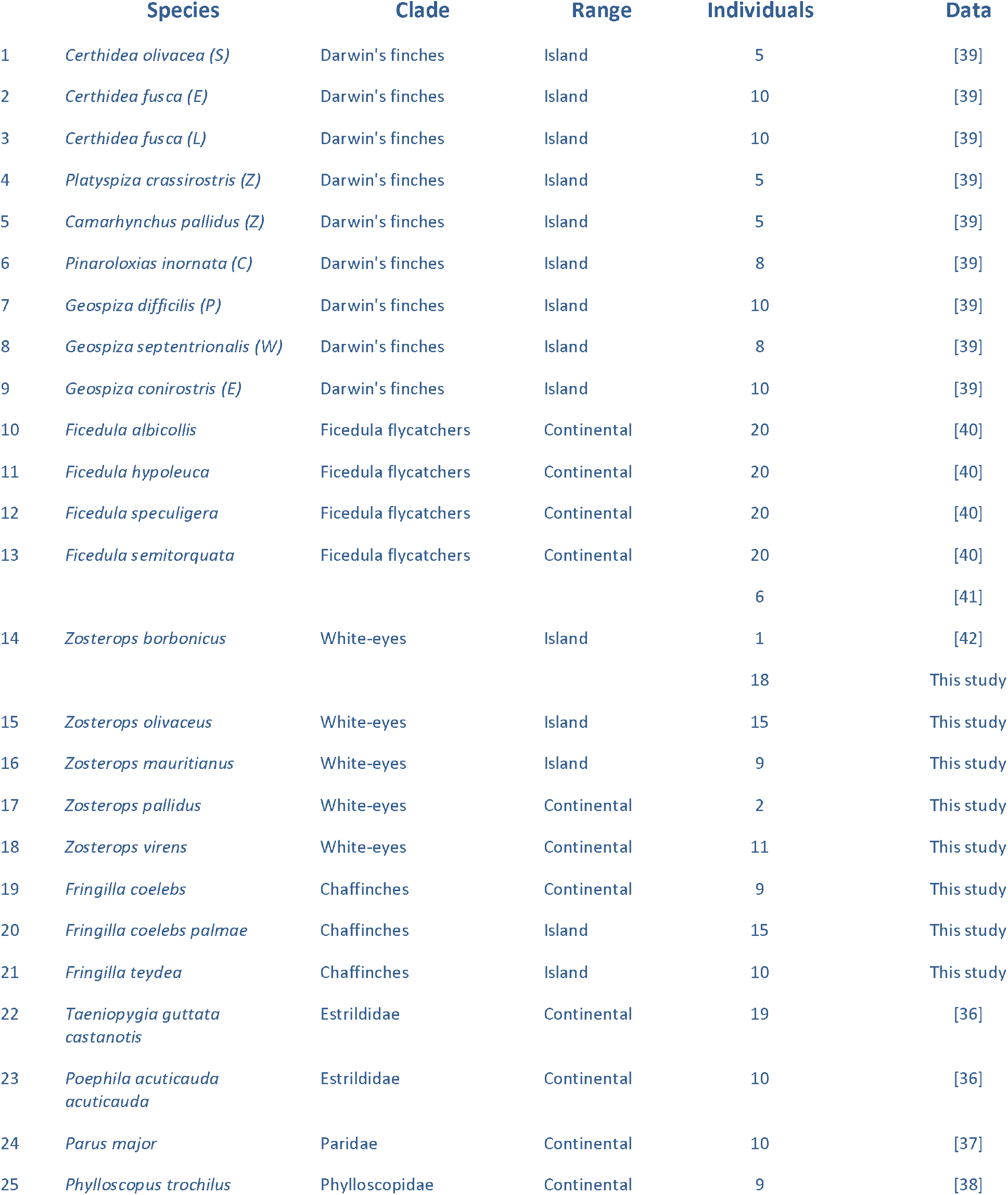
Sequencing data used in this study. The abbreviation in parenthesis following Darwin’s finches names indicate the island of origin (C=Coco, E=Española, L=San Cristobal, P=Pinta, S=Santiago, W=Wolf, Z=Santa Cruz, see also SI Methods S1)

|  | Species | Clade | Range | Individuals | Data |
| --- | --- | --- | --- | --- | --- |
| 1 | *Certhidea olivacea* (S) | Darwin's finches | Island | 5 | [39] |
| 2 | *Certhidea fusca* (E) | Darwin's finches | Island | 10 | [39] |
| 3 | *Certhidea fusca* (L) | Darwin's finches | Island | 10 | [39] |
| 4 | *Platyspiza crassirostris* (Z) | Darwin's finches | Island | 5 | [39] |
| 5 | *Camarhynchus pallidus* (Z) | Darwin's finches | Island | 5 | [39] |
| 6 | *Pinaroloxias inornata* (C) | Darwin's finches | Island | 8 | [39] |
| 7 | *Geospiza difficilis* (P) | Darwin's finches | Island | 10 | [39] |
| 8 | *Geospiza septentrionalis* (W) | Darwin's finches | Island | 8 | [39] |
| 9 | *Geospiza conirostris* (E) | Darwin's finches | Island | 10 | [39] |
| 10 | *Ficedula albicollis* | Ficedula flycatchers | Continental | 20 | [40] |
| 11 | *Ficedula hypoleuca* | Ficedula flycatchers | Continental | 20 | [40] |
| 12 | *Ficedula speculigera* | Ficedula flycatchers | Continental | 20 | [40] |
| 13 | *Ficedula semitorquata* | Ficedula flycatchers | Continental | 20 | [40] |
|  |  |  |  | 6 | [41] |
| 14 | *Zosterops borbonicus* | White-eyes | Island | 1 | [42] |
|  |  |  |  | 18 | This study |
| 15 | *Zosterops olivaceus* | White-eyes | Island | 15 | This study |
| 16 | *Zosterops mauritanus* | White-eyes | Island | 9 | This study |
| 17 | *Zosterops pallidus* | White-eyes | Continental | 2 | This study |
| 18 | *Zosterops virens* | White-eyes | Continental | 11 | This study |
| 19 | *Fringilla coelebs* | Chaffinches | Continental | 9 | This study |
| 20 | *Fringilla coelebs palmae* | Chaffinches | Island | 15 | This study |
| 21 | *Fringilla teydea* | Chaffinches | Island | 10 | This study |
| 22 | *Taeniopygia guttata castanotis* | Estrildidae | Continental | 19 | [36] |
| 23 | *Poephila acuticauda acuticauda* | Estrildidae | Continental | 10 | [36] |
| 24 | *Parus major* | Paridae | Continental | 10 | [37] |
| 25 | *Phylloscopus trochilus* | Phylloscopidae | Continental | 9 | [38] |

### Do island species exhibit genomic footprints consistent with low Ne?

Past effective population sizes were inferred using the Pairwise Sequentially Markovian Coalescent (PSMC) approach for one randomly selected individual from each species (Fig. S2) and were then averaged over the last one million years. The analyses confirmed that island species exhibit a significantly lower mean *Ne* than continental species over the last one million years (mean *Ne* = 362,456 and 94,944 for continental and island species respectively, Fig.1 A; log-transformed *Ne*, PGLS p-value=1.0 × 10^−4^). Specifically, inferred mean Ne values over the last million years range from 6.1 × 10^4^ for the Tenerife blue chaffinch (*Fringilla teydea*) to 1.2 × 10^6^ for a continental population of the common chaffinch (*F. coelebs*), representing a ∼20 fold difference (Fig. 1A & Table S2).

**Fig.1:**
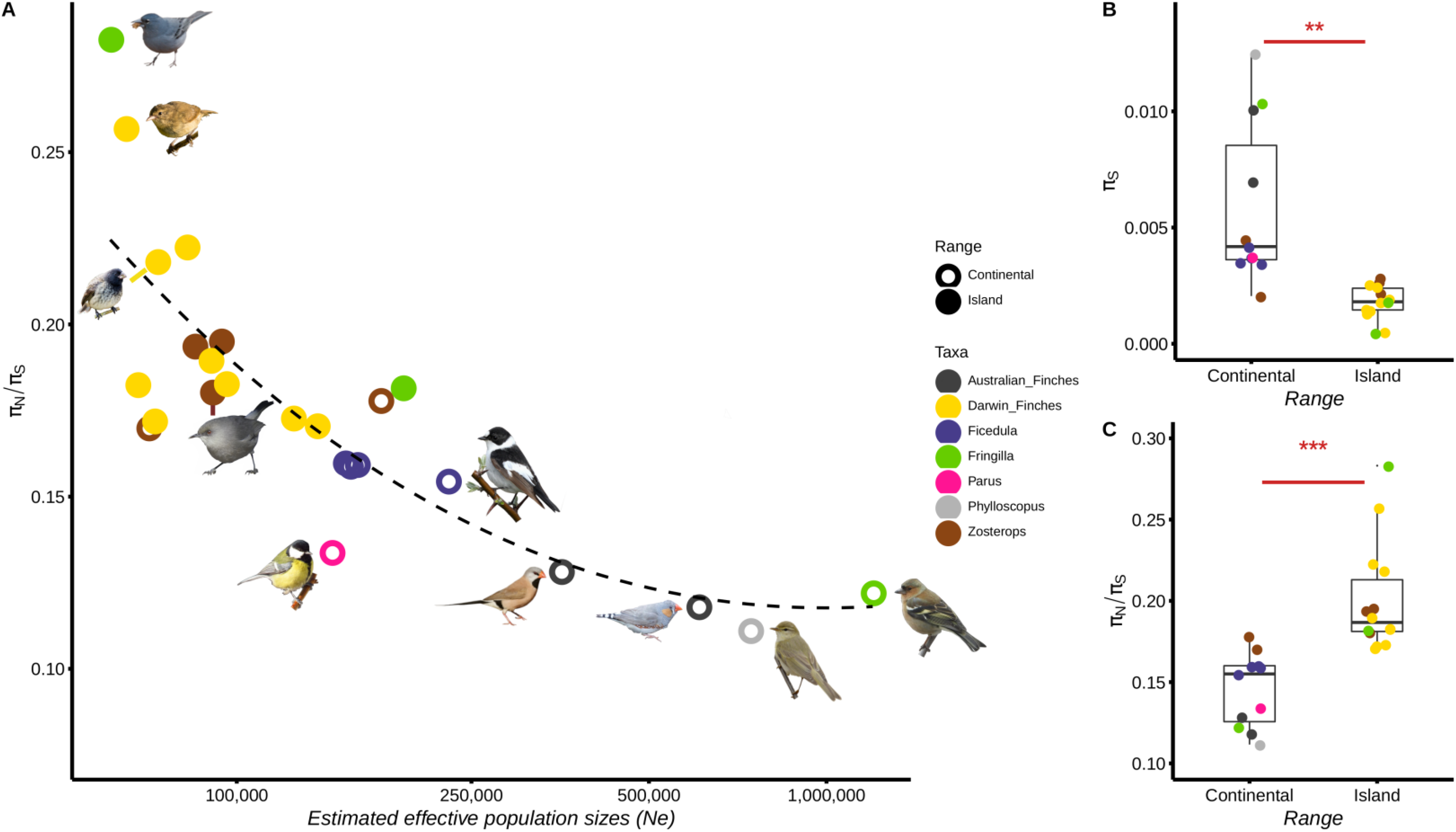
Island species as models for evolution in small effective population sizes. A. Local polynomial regression (LOESS with span=1.25) between the ratio of nonsynonymous to synonymous nucleotide diversity (π_N_/π_S_) and the mean effective population sizes over the last million years (Ne), as inferred using PSMC (see Fig. S3 for a log-log regression between π_N_/π_S_ and π_S_ estimates, respectively). B & C. Variation in nucleotide diversity (π_S_, B) and π_N_/π_S_ between island endemic and continental species (C).Photo credits: A. Chudý, F. Desmoulins, E. Giacone, G. Lasley, Lianaj, Y. Lyubchenko, B. Nabholz, J.D. Reynolds, K.Samodurov, A.Sarkisyan (iNaturalist.org); M. Gabrielli (personal communication).

Such long-term differences in Ne between insular and continental species are expected to generate differences in nucleotide diversity levels, because genetic variation is determined by both mutation rate and effective population size. By estimating nucleotide diversity at synonymous (π_S_) and at non-synonymous sites (π_N_), we find marked differences between island and continental species. Using 6,499 orthologous genes on average (range: 5,018-7,514, among 8,253 orthogroups [9]), we find that π_S_ varies from 0.07% in the Tenerife blue chaffinch to 1.25% in the willow warbler (*Phylloscopus trochilus*), representing a 17-fold difference between these island and continental species (Table S2 & Fig. S3). Island species exhibit significantly lower mean π_S_ than continental species (mean π_S_ = 0.59% and 0.18% for continental and island species respectively, Fig.1 B; PGLS, p-value=5.29 × 10^−3^).

In addition to strong evidence for lower Ne in island species, we also find lower census population sizes in the island species (island: 7 species, median: 1.1 × 10^4^ (range: 6.2 × 10^2^ - 3.0 × 10^5^); continental: 6 species, median: 2.5 × 10^8^ (range: 2.0 × 10^5^ - 5.7 × 10^8^); log-transformed census sizes, PGLS, p-value=8.29 × 10^−5^). Furthermore, both log_10_-transformed current census population sizes and geographical range in square kilometers are positively correlated with π_S_ (Fig.2A,C; PGLS, p-value < 0.01, Table S1). Taken all together, these results provide strong support for the view that long-term restrictions on census population sizes due to the limited surface area available to island species constrains the upper bound of effective population size.

### Are deleterious mutations segregating more in island species?

Based on the nearly neutral theory of molecular evolution, the higher level of genetic drift associated with lower Ne is expected to contribute to an accumulation of slightly deleterious mutations in island species relative to their continental counterparts. Using the ratio of non-synonymous to synonymous mutations (π_N_/π_S_) as a proxy for the proportion of these slightly deleterious mutations, we recover, on average, a 40% higher π_N_/π_S_ in island species than in continental species (Fig. 1 C; mean π_N_/π_S_ = 0.145 and 0.201 for continental and island species respectively, PGLS p-value=4.57 × 10^−3^).

In addition, we find substantial within-genome variation in the accumulation of slightly deleterious mutations, as well as in the levels of nucleotide diversity, in such a way that π_S_ and π_N_/π_S_ are respectively positively and negatively correlated to the GC content at the third codon position (hereafter, GC3, Note S2). These correlations are found to be stronger in island species relative to their continental counterparts, with a particularly pronounced difference in π_N_/π_S_ in genes exhibiting a low GC3 (Note S2). Recombination limits genetic interactions between selected mutations and can therefore improve the efficiency of selection [10-11]. Given that GC3 provides a robust proxy of recombination rate in birds [12-13], these results suggest that the intensity of the differences between island and continental species in the effectiveness of purifying selection relies heavily on the local genomic context.

We found strong negative correlations between (i) π_N_/π_S_ and the log_10_-transformed *Ne* averaged over the last one million years (Fig.1A; PGLS, p-value = 1.0 × 10^−4^) and (ii) the non-transformed π_N_ /π_S_ and π_S_ values (PGLS, p-value = 4.85 × 10^−5^). Log_10_ - transformed current census population sizes, as well as geographical range sizes, significantly correlate with π_N_/π_S_ (Fig.2). In contrast, the IUCN red list assessments have no effect on π_N_/π_S_ or π_S_ (Table S1, PGLS p-value >> 0.05). Taken together, our results provide strong empirical evidence that differences in census population sizes between island and continental species translate into differences in Ne, and that these differences have a marked influence on genetic diversity and the efficiency of natural selection. These findings fit remarkably well with the expectation from the nearly neutral theory.

**Fig. 2:**
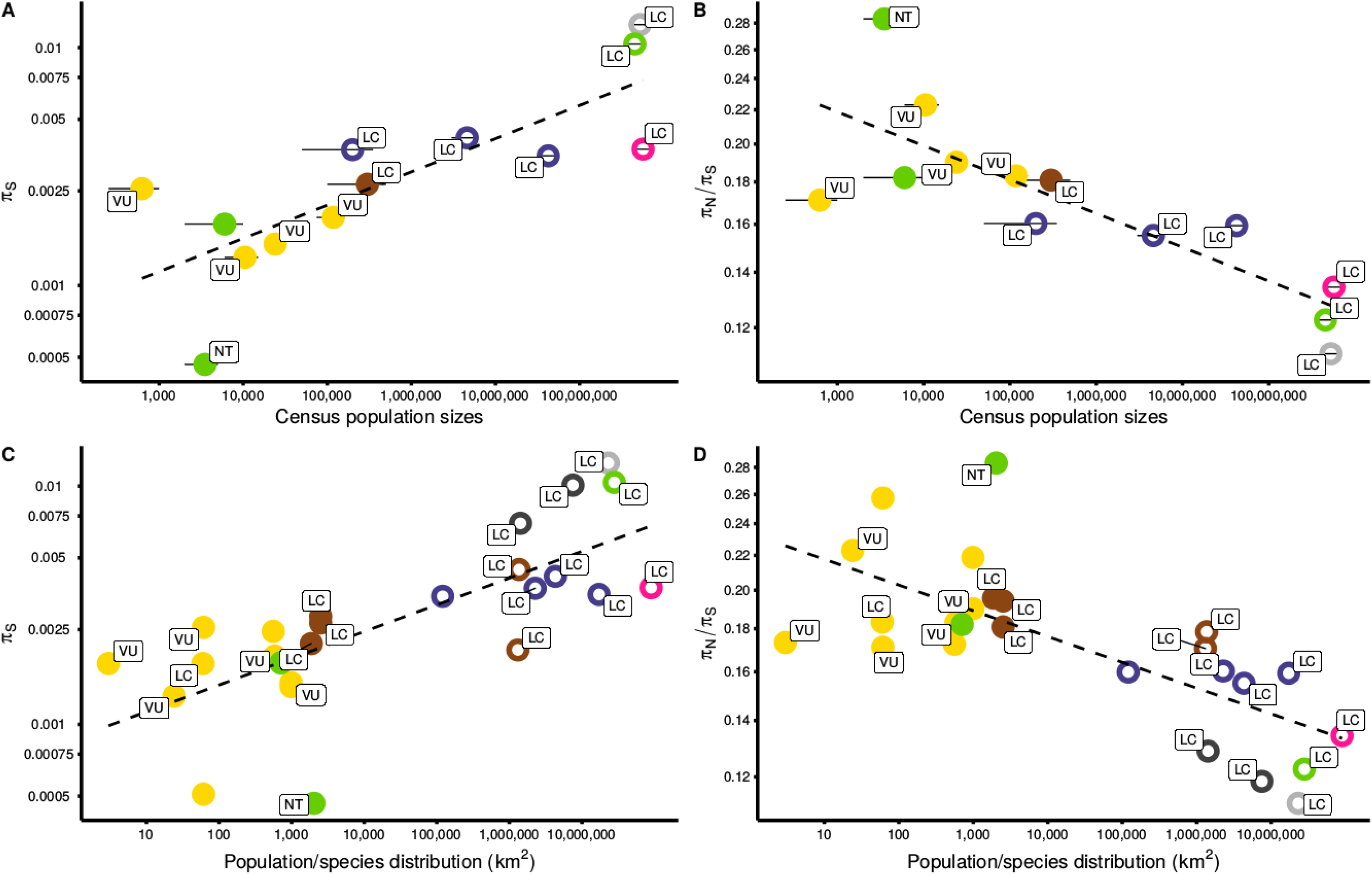
Ecological-evolutionary correlations based on the variables investigated in this study. π_S_ and π_N_/π_S_ are used as proxies of Ne and the efficiency of natural selection to remove deleterious variants and are correlated with both the median estimates of the current census population sizes (A & B) and the geographical range sizes (C & D). Both ecological and evolutionary parameters are log-transformed. Filled and open dots represent the island and the continental species, respectively (Fig. 1 for details). Only the 13 species with estimates of the current census population sizes are included for the panels A & B (with ranges shown with a thin black line). Where known, the IUCN conservation status of the investigated species is indicated (LC = least concerned, NT = near threatened, VU = vulnerable)

### Do insular species show lower adaptive potential?

Theory predicts that lower Ne in island species should lead to a lower rate of adaptive substitutions than in continental species, if adaptation is limited by the supply of new mutations [8] and/or if slightly advantageous mutations become effectively neutral in low *Ne species* [6]. For taxa with at least two species (*i*.*e*. all except *Parus* and *Phylloscopus*), we used the maximum likelihood method implemented in Grapes [14] to estimate non-adaptive rate of substitution (ω_NA_) and adaptive rate of substitution (ω_A_) with ω (i.e. d_N_/d_S_) being the sum of ω_NA_ + ω_A_. No significant difference in ω was observed between island and continental species (ω island = 0.194 and ω continental = 0.187). By contrast, island species showed a higher ω_NA_ (Δmean_continental *vs*. Island_ = 0.063) and a lower ω_A_ (Δmean_continental *vs*. Island_ = 0.056) (Fig. 3; see Sup Fig S4 for α estimates) than continental counterparts. However, these differences are only significant for tests that did not explicitly take phylogenetic structure into account (PGLS: p-value = 0.257 and p-value = 0.237; non-PGLS p-value = 0.014 and p-value = 0.002 for ω_A_ and ω_NA_ respectively, Table S1) and therefore they should be interpreted with caution.

**Fig. 3:**
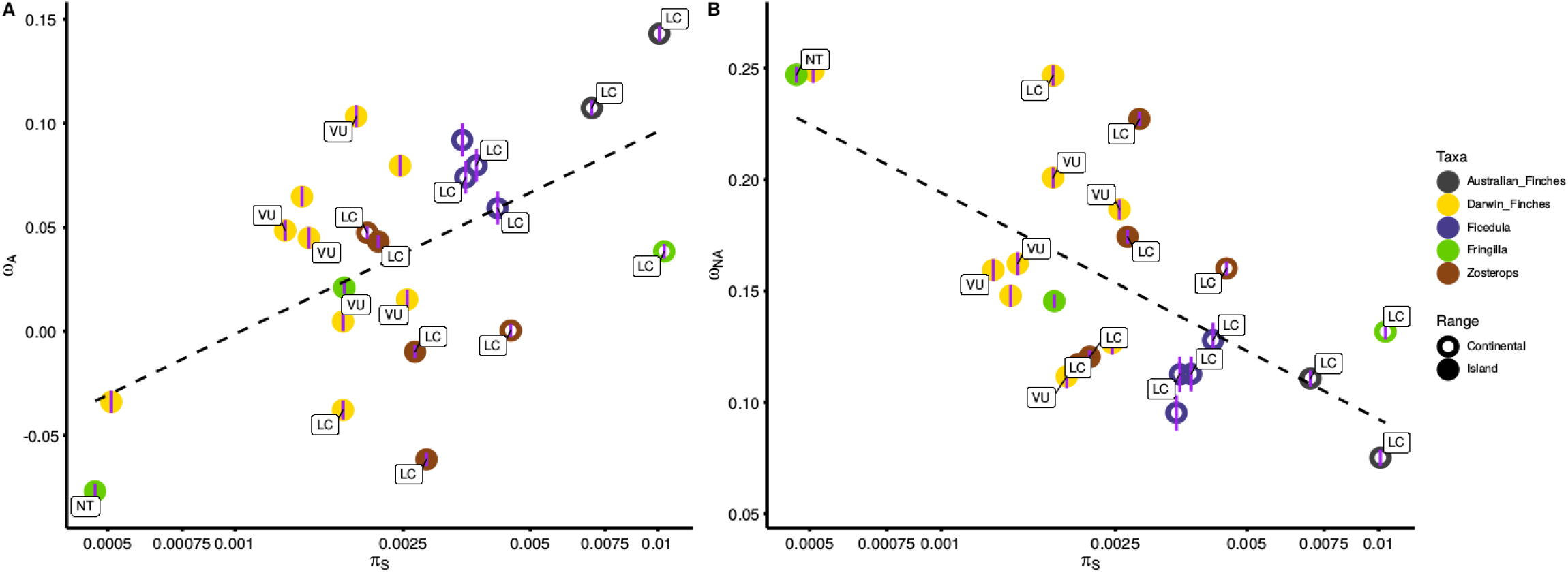
Proportion of adaptive (A) and non-adaptive (B) substitutions along the neutral genetic diversity gradient (π_S_) as estimated by comparing the observed and the expected d_N_/d_S_ under near neutrality assuming the polymorphism data using the DFE-⍰ method (α shown Fig. S4; with ω_A_ = α(d_N_/d_S_) & ω_NA_ = (1-α)(d_N_/d_S_)). Estimates were performed using all sites and the GammaExpo model. Error bars (purple line) represent the 95% confidence intervals of each estimate under this model. Where known, the IUCN conservation status of the investigated species is indicated (LC = least concerned, NT = near threatened, VU = vulnerable).

We found that ω_A_ was positively correlated with log_10_-transformed π_S_ (PGLS p-value = 0.029; Fig. 3A), and negatively correlated with the log_10_ -transformed π_N_/π_S_ (PGLS, p-value = 0.034; Table S1). Reciprocally, ω_NA_ is significantly negatively correlated with log10-transformed π_S_ (PGLS, p-value = 0.020; Fig. 3B) and positively with log_10_ -transformed π_N_/π_S_ (PGLS, p-value = 0.025; Fig. S4).

Overall, our analysis suggests that a lower *Ne* doubly affects island species relative to continental species, because (i) relatively fewer adaptive mutations can reach fixation, and (ii) the lower efficiency of natural selection allows a greater proportion of weakly deleterious variants to reach fixation in insular species.

## Discussion

Our analysis of whole-genome resequencing data has allowed us to find lower nucleotide diversity, a higher frequency of slightly deleterious mutations and lower adaptive substitution rates in the island species than in the continental ones. These results provide important insights for evolutionary biology and they also have major implications for the conservation of species with small populations.

### Island species as models for studying the evolutionary consequences of small Ne

The smaller land area available on oceanic islands should constrain the upper bound of both census and effective population sizes of insular species, to such an extent that demography affects the ability of purifying selection to remove weakly deleterious mutations. Our results are largely consistent with this general hypothesis and suggest that contemporary census sizes provide information on long-term *Ne* (but see also [15-16]). For most population genomic estimates we investigated, including π_S_, π_N_/π_S_ and PSMC-inferred *Ne*, we observed significant differences between continental and island species that are consistent with theoretical expectations.

Previous taxon-specific studies have reported low *Ne* in a diverse range of island organisms (e.g. Giant Galápagos tortoises [17], woolly mammoths [18], island foxes [19-20], *Corvus* [21]). Therefore, it is very likely that island species predominantly exhibit lower *Ne* than their more abundant, broadly distributed, mainland relatives, and this pattern may not be restricted to some specific animal clades such as birds or mammals, but may also be true for a large range of taxa (e.g. [22] for plants). More broadly, this result opens up new opportunities for using island species as models to understand the impact of *Ne* on genome evolution in natural populations, including genome size, or of natural selection on non-coding genomic regions.

### Broad support for the nearly neutral theory of molecular evolution

Fifty years after the introduction of the neutral theory of molecular evolution by Kimura [23] and King and Jukes [24], and after being extended into the nearly neutral theory [4], the neutralist–selectionist controversy remains one of the sharpest and most polarized debates in biology. Based on our large genome-scale empirical data, our results match theoretical expectations of the nearly neutral theory remarkably well. This is consistent with the strength of this theory in explaining patterns of DNA sequence evolution, allowing us to affirm that the nearly neutral theory is overwhelmingly supported by our dataset. Slightly deleterious mutations are frequent and become effectively neutral when the effect of genetic drift increases, as is typically observed in insular species.

Selective processes, including positive selection on beneficial alleles and background selection, play an important role in the sequence evolution of the investigated species, but cannot be used to reject the theory as a whole. Empirical investigations found that the proportion of adaptive substitutions does not overall scale with *Ne* when distant taxa are considered all together (e.g. [14]), but taxa-specific investigations were able to find such a relationship, with a lower proportion of adaptive substitutions in species with a lower *Ne*, as recently reported for several groups of animals [8]. Firstly, our analyses provide additional evidence for such a relationship in passerine birds. Secondly, we indeed observe that local recombination rates influence both local levels of nucleotide diversity and the number of deleterious mutations (Note S2), which is consistent with heterogeneous landscapes of *Ne* throughout genomes [11]. However, significant differences between island and continental species were similarly recovered in both lowly and highly recombining regions of the genome, supporting the claim that background selection does not fundamentally change the predictions that can be drawn from the theory.

### Ecological-evolutionary ties and perspectives

At the macroevolutionary scale, strong correlations between life-history traits and both levels of polymorphism and ratios of non-synonymous to synonymous mutations have been reported in the literature for both animals and plants [25-27], suggesting that determinants of genetic diversity are mostly ecologically-driven. We found that nucleotide diversity scales positively with species range, which therefore suggests a gradual transition between species restricted to small islands and species widely distributed over continents. Recently, Peart et al. [16] proposed that conservation priorities should be defined based on the ratio of census size to *Ne*. However, whether population genomic estimates of *Ne* are informative enough to assess conservation status is questionable. A general outcome is that animal species classified as threatened generally exhibit lower genetic diversity than those classified as non-threatened, including birds (at least at microsatellite loci [28-29]). Based on our whole-genome analyses, we can report no obvious contrast between the four island species classified as threatened (vulnerable status) and the species classified as non-threatened, neither for the levels of nucleotide diversity nor for their efficiency of natural selection (but see [30]). Díez-del-Molino et al. [15] were also unable to recover a significant effect of the IUCN assessment on the levels of nucleotide diversity in birds and mammals. Using 78 mammal species, Brüniche-Olsen et al. [31] only recovered this pattern when the animals’ diets were explicitly taken into account. Consequently, it seems that we still have a long way to go towards precisely describing whether these genomic features are completely independent or are correlated to some extent with the current conservation status.

Another open question is whether population genomics can provide information so that short-term IUCN objectives can be extended over a longer timeframe? Even if some island species accumulate slightly deleterious mutations [32], supposedly leading to increased maladaptation, we can question whether this burden of slightly deleterious mutations can lead to species extinction. This hypothesis holds true only if these deleterious mutations are neither purged nor opposed by compensatory or beneficial mutations [33]. Remarkably, the four species classified as threatened are not those exhibiting the lowest proportion of adaptive substitutions (mean ω_A_=0.053 compared to 0.036 for the 15 species with a Least Concern status). Recent macro-evolutionary investigations, however, provide support for this increased risk of (i) being endangered depending on the time since the species colonized the island [34] or (ii) becoming extinct depending on the island size [35]. Age-dependent processes such as ecological specialization were proposed, but the accumulation of deleterious mutations might explain this phenomenon as well. Rogers & Slatkin [18] propose that, after a tipping point, this mutational meltdown might contribute to the ultimate steps in the road to extinction. Endemic island species therefore represent taxa of high interest in the evaluation of the long-term consequences of evolution under low effective population sizes.

## Materials & Methods

### SUMMARY

#### Data used in this project

- Newly sequenced data (see also Table S3, sequencing data: BioProject PRJNA661201):

- 18 Zosterops borbonicus samples (Reunion Island, France)
- 15 Zosterops olivaceus samples (Reunion Island, France)
- 9 Zosterops mauritinaus samples (Mauritius)
- 1 Zosterops pallidus sample (South Africa)
- 11 Zosterops virens samples (South Africa)
- 9 continental Fringilla coelebs samples (continental Spain)
- 15 Fringilla coelebs palmae samples (La Palma, Canary Islands, Spain)
- 10 Fringilla teydae samples (Tenerife, Canary Islands, Spain)

- Publicly available data used in this study (see also Table S3):

- Taeniopygia & Poephila ([36], BioProject PRJEB10586)
- Parus ([37], BioProject PRJNA381923)
- Phylloscopus ([38], BioProject PRJNA319295)
- Darwin’s finches ([39], BioProject PRJNA263122)
- Ficedula ([40], BioProject PRJEB7359)
- Zosterops ([41-42, BioProjects PRJEB18566, PRJNA530916)

#### Bioinformatic softwares used

- Meraculous ([43], https://jgi.doe.gov/data-and-tools/meraculous/)
- HiRise ([44], https://github.com/DovetailGenomics/HiRise_July2015_GR)
- Trimmomatic ([45], http://www.usadellab.org/cms/?page=trimmomatic)
- BWA mem ([46], http://bio-bwa.sourceforge.net/)
- Picard ([47], http://broadinstitute.github.io/picard/)
- GATK (v. 3.7, [48], https://gatk.broadinstitute.org/hc/en-us)
- MitoFinder ([49], https://github.com/RemiAllio/MitoFinder)
- Macse ([50], https://bioweb.supagro.inra.fr/macse/index.php?menu=releases)
- IQTREE ([51], http://www.iqtree.org/) genBlastG ([52], http://genome.sfu.ca/genblast/download.html)
- HMMER toolkit ([53], http://hmmer.org/)
- PSMC ([54], https://github.com/lh3/psmc)
- Grapes ([14], https://github.com/BioPP/grapes)
- R (v.3.6.3, [55], https://cran.r-project.org/)
- Scripts used for this study (https://osf.io/uw6mb/)

## SPECIES

In this study, we both reanalyzed publicly available data and generated our own sequencing data from 25 passerine species (Table S3). By generating new sequencing data, our objective was to target taxa containing both island and continental relatives (chaffinches and white-eyes) in order to increase our statistical power. More broadly, our comparison is only based on species with relatively similar body-mass, longevity and clutch-size. This control was introduced to reduce the risk of some confounding factors that could correlate with *Ne* [27] in order to be able to truly assess the effect of insularity.

Species range sizes were obtained from BirdLife (http://datazone.birdlife.org/) or estimated based on the information shown on the IUCN-red list webpage using CalcMaps (https://www.calcmaps.com/map-area/). For endemic island species, we considered the total island area as a maximum bound for the population range. The IUCN red list conservation status is given at the species level and not at below-species level. As a consequence, we either considered this information to be missing for both populations (e.g. *Certhidea fusc*a E and *C. fusca* L) or we only used the status for the most widely distributed species (e.g. the Least Concern (LC) status for the population with a large continental distribution rather than for the island one as in *F. coelebs palmae*). For *Ficedula speculigera*, a species with a D_A_>0.002 (Methods S1) and recognized as a distinct species from *F. hypoleuca*, no information is yet available in the IUCN red list database.

## METHODS USED IN THIS STUDY

### DNA extraction and sequencing (*Zosterops* and *Fringilla* species)

All *Zosterops* and *Fringilla* individuals were captured using mist nets. With the exception of African *Zosterops* species (*Z. pallidus* and *Z. virens*, see below), we collected blood samples for each bird by venipuncture of the brachial vein and stored blood in absolute ethanol at −20°C until DNA extraction. For African species, *Z. pallidus* and *Z. virens* individuals, DNA was extracted from liver, muscle or blood. For these samples, voucher specimens are stored at the Museum National d’Histoire Naturelle (MNHN), Paris, France and a tissue duplicate is deposited in the National Museum Bloemfontein (South Africa). For all *Zosterops* and *Fringilla* samples, total genomic DNA was extracted using the DNeasy Blood and Tissue kit (QIAGEN, Valencia, CA) following the manufacturer’s instructions. Library preparation (1.0 µg DNA used per sample) and Illumina high-throughput sequencing using a paired-end 150 bp (PE150) strategy were performed at Novogene (Cambridge, UK) to a minimum sequencing yield of 18 Gb per sample (i.e., ∼15X coverage). Details on samples are available in Data S1. For these species, we used exactly the same approach as for the publicly available data for variant identification and sequence reconstruction strategies, as described in Methods S1. All newly sequenced raw reads are available under the SRA BioProject accession number PRJNA661201.

### Publicly available sequencing data

We collected publicly available raw sequencing data on SRA from a large range of studies (Table S3). The phylogenetic relationships among Darwin’s finches are not fully resolved [39;56-57], so we first evaluate the net divergence between all pairs of species to delimit 9 groups of species with a net divergence (D_A_ > 0.1%; see SI Methods S1). Within each group, we selected a single population based on the number of sequenced individuals that were publicly available [39]. Variant identification and sequence reconstruction steps are described in Methods S1.

### Variant identification

We used Trimmomatic (v.0.33;[45]) to remove adapters, stringently trim and filter reads using the following set of parameters: LEADING:3 TRAILING:3 SLIDINGWINDOW:4:15 MINLEN:50. All trimmed reads were then mapped against the reference genome for each clade (see above) with BWA mem (v. 0.7.12;[46]) using default settings. Unmapped reads and mapped reads with a quality (MQ) below 20 were then discarded. Potential PCR duplicates were then flagged using MarkDuplicates v. 1.140 (Picard tools, [47]). Variant calling was then performed using GATK (v. 3.7; [48]). First, we used HaplotypeCaller on single samples (gVCF) to call SNPs using default parameters. For each species, we then performed a joint genotyping (“GenotypeGVCFs”). To ensure high quality in our dataset, we filtered out low-quality SNPs using several settings: a quality by depth (QD) < 2.0, a Fisher Strand (FS) bias >60, a mapping quality (MQ) <40, a MQranksum < −2 or a ReadPosRankSum < −2 or a Raw Mapping Quality (Raw_MQ) < 45,000. SNPs satisfying one or more of these conditions were discarded. For every group of species, we performed principal component analyses (PCA) based on a random sampling of SNPs over the genome (50-200k) to capture additional levels of population structure or an unfortunate misnaming of an individual that could have occurred at some point between the bird sampling campaign and the analysis of the raw sequencing data.

### Gene models & orthology prediction

We used one reference genome for all species belonging to the same clade (Table S3). We used the genome and gene models of a medium ground-finch individual (*Geospiza fortis*; assembly GeoFor_1.0; GCF_000277835 [58]) for all Darwin’s finches, a collared flycatcher (*Ficedula albicollis*, GCF_000247815; assembly FicAlb_1.4 [59]) for all *Ficedula*, a zebra finch (*Taeniopygia guttata*; GCF_000151805; assembly taeGut3.2.4 [60]) for the Estrildidae, a Reunion grey white-eye (*Zosterops borbonicus*; GCA_007252995; assembly ZoBo_15179_v2.0 [42]) for all *Zosterops*. We also used the genome of the willow warbler (*Phylloscopus trochilus*; GCA_002305835; assembly ASM230583v1 [38]) and the great tit (*Parus major*; GCF_001522545.2; assembly Parus_major1.1 [61]). For all the investigated chaffinches, we used a newly generated assembly of *Fringilla coelebs* (Methods S1, version “HiRise” of [62]). For this latter species, as well as for the willow warbler (*P. trochilus*), no gene models were available and we therefore first performed a protein homology detection and intron resolution using genBlastG [52] (http://genome.sfu.ca/genblast/download.html) with the following options “-p genblastg - c 0.8 -r 3.0 -gff -e 1e-10”.

To analyse the same orthologous sequences in all species, we used the set of 8253 orthologs identified by Jarvis et al. [9] (http://gigadb.org/dataset/101041). Then, we added the sequence of our species to this set of orthogroups using the method described in Scornavacca et al. [63]. Briefly, each orthogroup was used to build an HMM profile using the HMMER toolkit [53]. Then, for each new sequence, *hmmscan* was used on the HMM database to get the best hits among the orthogroups. For each orthogroup, the most similar sequences for each species were then detected via *hmmsearch*. Outputs from *hmmsearch* and *hmmscan* were considered to be accurate if the first hit score was substantially better than the second best one (in order to limit the risk of paralogy), following a best-reciprocal-hit approach when the results of both programs were compared [63].

### Effective population size estimates

Historical demographic variations in *Ne* were estimated using the Pairwise Sequentially Markovian Coalescent (PSMC) model implemented in the software PSMC [48]. Fasta sequences were converted to the PSMC fasta format using a C++ program (Fasta2PSMCFasta: https://osf.io/uw6mb/) written using BIO++ library [64]. Only scaffolds longer than 500Kb were considered. We used block length of 100bp, with no more than 20% of missing data per block, as implemented in “fq2psmcfa” (https://github.com/lh3/psmc).

For each species, PSMC analyses were run using two randomly selected individuals. To identify suitable parameters, several -t and -p parameters were tested including -p “4+30*2+4+6+10” (as in [65]) and -p “4+25*2+4+6” (as in [66]) but also -p “4+10*3+4” and -p “5*1+25*2+6”. The best combination (t15 -r4 -p “5*1+25*2+6”) was manually chosen after excluding other parameter values leading to large differences between the two individuals from the same species. Then, we randomly selected one individual and excluded the first four atomic time intervals to exclude the noisy estimates generally generated by PSMC for very recent times and therefore strengthen the reliability of the average estimates of *Ne* over the last million years.

Time was scaled assuming a mutation rate of 4.6 × 10^−9^ mutation/site/generation as estimated [67] and a generation time of 2 years [65; 68]. Results were plotted in R (v3.6.3 [55]) using the function “psmc.results” [69; https://doi.org/10.5061/dryad.0618v/4] and with the R packages ggplot2 [70] and cowplot [71].

### Summary statistics of the polymorphic data

π_S_ and π_N_/π_S_ ratios were computed using *seq_stat_coding* from reconstructed fasta sequences (Methods S1) using a publicly available Bio++ script and a procedure previously described (42, https://osf.io/uw6mb/). We empirically validated that our π_S_ and π_N_/π_S_ estimates were not impacted by the variable number of samples per species (Note S2). In addition, we used the π_N_/π_S_ estimates based on the site frequency spectra at both non-synonymous and synonymous sites as described in Rousselle et al. [11] to check the accuracy of these estimates (see Note S2). Guanine-Cytosine (GC) content at third-codon positions of protein-coding genes (hereafter GC3), an excellent proxy of the local recombination rate in birds [72] was also computed under *seq_stat_coding*. To estimate the within-genome variation in the efficacy of selection (Note S2), we estimated π_N_/π_S_ on sets of genes representing a total concatenated coding alignment of 2 Mb, after sorting genes by ascending values of GC3. The last window corresponding to genes exhibiting the highest GC3 values was only considered if this window contained at least 1 Mb of coding sequence.

### Summary statistics of the divergence data

We used the method implemented by Galtier [14] (Grapes. v1.0) to estimate α, ω_A_ and ω_NA_ using the approach introduced by Eyre-Walker & Keightley [73]. Briefly, we fitted both a negative Gamma distribution and an exponential distribution to the synonymous and non-synonymous Site Frequency Spectrum (SFS) (the so-called GammaExpo model [14]) to model the distribution of fitness effect (DFE). Fitted parameters of the DFE were then used to compute the expected d_N_/d_SS_ under near neutrality (i.e. without adaptive substitutions but including weakly deleterious substitutions), which was compared to the observed d_N_/d_S_ to estimate the adaptive substitution rate (ω_A_) and the proportion of adaptive substitutions (α) [with ω_A_ = α(d_N_/d_S_) & ω_NA_ = (1-α)(d_N_/d_S_)]. Potential recent changes in population size that affect the SFS were taken into account via the use of nuisance parameters capturing distortions of the SFS optimized alongside the DFE parameters [74].

### Statistical analyses

All statistical analyses were performed using R [55]. We only considered models with a similar number of observations and compared these models based on the Akaike information criterion with a correction for small sample sizes (AICc). To test for the influence of the explanatory variables on π_S_ and π_N_/π_S_, we used Phylogenetic Generalized Least Square (PGLS) models. Explanatory variables were always log_10_-transformed as this violated less frequently the assumption of normality, heteroscedasticity and independence of the residuals using a simple linear model. For PGLS, we used the model implemented in the “nlme” package [75]. The mitochondrial phylogeny was considered as the species tree (see Methods S4) taken into account assuming a Brownian correlation structure (using “corBrownian” from the “ape” package; [76]). P-value and AICc were computed using the anova.gls function. The rationale of the phylogenetic control is to account for the shared polymorphisms (part of species similarity that is explained by the inheritance from a common ancestor). The level of polymorphism of a given species is dynamically controlled by drift, mutation rate and natural selection. As soon as two species do not share a significant fraction of their polymorphism, there is no need to account for their phylogenetic proximity because their polymorphisms evolved independently. Therefore, the results of all the tests including with and without phylogenetic controls, transformed and untransformed π_S_ and π_N_/π_S_ are presented in Table S1.

R plots were generated using a series of R packages: cowplot [71], ggplot2 [70], ggpubr [77], ggrepel [78] and ggtree [79].

## Acknowledgements

This research was funded by the French ANR (BirdIslandGenomic project, ANR-14-CE02-0002). The analyses benefited from the Montpellier Bioinformatics Biodiversity (MBB) platform services, the genotoul bioinformatics platform Toulouse Midi-Pyrenees (Bioinfo Genotoul) and the Biogenouest BiRD core facility (Université de Nantes). We are grateful to Quentin Rougemont for providing feedback on a previous version of the manuscript, colleagues from the phylogeny and molecular evolution team at ISEM Montpellier for fruitful discussions throughout this project and Neil McNair (University of Vienna) for proofreading the manuscript, as well as three anonymous reviewers for their constructive comments, which helped us to substantially improve the manuscript. A previous manuscript version also benefited from detailed discussions on Twitter, we would thus like to thank Graham Coop, Jenny James, Nicolas Rode, among others, for their helpful suggestions. The authors wish to thank all the farmers, private and national nature reserves where fieldwork was conducted. For South African samples, the handling and sampling protocols were approved by the Comité Cuvier (68-055 to JF). We are grateful to the provincial authorities in the Eastern Cape and Free State provinces of South Africa, and Eastern Cape Parks (Alan Southwood, Cathy Dreyer, Gavin Shaw, Sizwe Mkhulise) for granting permission to collect samples (permit numbers RA-190, CRO144/14CR and 01-24158). We would also like to acknowledge the Percy FitzPatrick Institute (University of Cape Town), the National Museum Bloemfontein (Free State), R.C.K. Bowie, P.-H. Fabre, E. Kolarova and G. Oatley for help during field work and logistical support. For Reunion Island samples, we thank the Reunion National Park for granting us permission to conduct fieldwork in Pas de Bellecombe, Reunion Island, France and the field station of Marelongue, funded by the P.O.E., Reunion National Park and OSU Reunion, for logistical support. For Spanish samples, the Canary government gave permission to perform the sampling work to JCI (permit 01-24158). JCI was funded by the Spanish Ministry of Science, Innovation and Universities (Ref.: PGC2018-097575-B-I00) and by a GRUPIN research grant from the Regional Government of Asturias (Ref.: IDI/2018/000151). BM and MR were partly funded by the Spanish Ministry of Science and innovation (grant PGC2018-098897-B-I00), and MR was supported by a doctoral fellowship from the Spanish Ministry of Education, Culture, and Sport (FPU16/05724). This is publication ISEM 2020-321.

## Author Contributions

T.L designed the research, conducted the analyses and wrote the paper. B.N. designed the research, contributed to the analyses, supervised the work and wrote the paper. M.R. contributed to the analysis of the divergence data. M-K.T. performed the wet lab work. A.E.C. performed preliminary investigations using the publicly available data and performed the analysis to identify the Darwin’s finch populations and species to consider for the paper. C.S. conducted the analyses of orthologous assignation. M.R.C., G. B. and B.M. provided the sequences data of *Fringilla coelebs* species and the reference *Fringilla* genome. J.C.I. provided the *Fringilla* samples used in this study. D.H.D.S. and J.F. helped for field work to collect mainland and island *Zosterops* samples. C.T and B.M. helped during filed works in Réunion and revised the manuscript. All the authors read and approved the manuscript.

## Notes

### Competing Interest Statement

The authors have declared no competing interest.

## References

1. Darwin, C., 1809-1882 (1859). On the origin of species by means of natural selection, or the preservation of favoured races in the struggle for life (Londonlll: John Murray, 1859)

2. Mayr, E. (1999). Systematics and the Origin of Species, from the Viewpoint of a Zoologist (Harvard University Press)

3. Ohta, T. (1973). Slightly Deleterious Mutant Substitutions in Evolution. Nature 246, 96–98.

4. Ohta, T. (1992). The Nearly Neutral Theory of Molecular Evolution. Annu. Rev. Ecol. Syst. 23, 263–286.

5. Gossmann, T.I., Keightley, P.D., and Eyre-Walker, A. (2012). The Effect of Variation in the Effective Population Size on the Rate of Adaptive Molecular Evolution in Eukaryotes. Genome Biology and Evolution 4, 658–667.

6. Lanfear, R., Kokko, H., and Eyre-Walker, A. (2014). Population size and the rate of evolution. Trends in Ecology & Evolution 29, 33–41.

7. Nam, K., Munch, K., Mailund, T., Nater, A., Greminger, M.P., Krützen, M., Marquès-Bonet, T., and Schierup, M.H. (2017). Evidence that the rate of strong selective sweeps increases with population size in the great apes. Proc Natl Acad Sci USA 114, 1613.

8. Rousselle, M., Simion, P., Tilak, M.-K., Figuet, E., Nabholz, B., and Galtier, N. (2020). Is adaptation limited by mutation? A timescale-dependent effect of genetic diversity on the adaptive substitution rate in animals. PLOS Genetics 16, e1008668.

9. Jarvis, E.D., Mirarab, S., Aberer, A.J., Li, B., Houde, P., Li, C., Ho, S.Y.W., Faircloth, B.C., Nabholz, B., Howard, J.T., et al. (2014). Whole-genome analyses resolve early branches in the tree of life of modern birds. Science 346, 1320.

10. Hill, W.G., and Robertson, A. (1966). The effect of linkage on limits to artificial selection. Genetical Research 8, 269–294.

11. Rousselle, M., Laverré, A., Figuet, E., Nabholz, B., and Galtier, N. (2018). Influence of Recombination and GC-biased Gene Conversion on the Adaptive and Nonadaptive Substitution Rate in Mammals versus Birds. Molecular Biology and Evolution 36, 458– 471.

12. Backström, N., Forstmeier, W., Schielzeth, H., Mellenius, H., Nam, K., Bolund, E., Webster, M.T., Ost, T., Schneider, M., Kempenaers, B., et al. (2010). The recombination landscape of the zebra finch Taeniopygia guttata genome. Genome Res 20, 485–495.

13. Kawakami, T., Smeds, L., Backström, N., Husby, A., Qvarnström, A., Mugal, C.F., Olason, P., and Ellegren, H. (2014). A high-density linkage map enables a second-generation collared flycatcher genome assembly and reveals the patterns of avian recombination rate variation and chromosomal evolution. Mol Ecol 23, 4035– 4058.

14. Galtier, N. (2016). Adaptive Protein Evolution in Animals and the Effective Population Size Hypothesis. PLOS Genetics 12, e1005774.

15. Díez-del-Molino, D., Sánchez-Barreiro, F., Barnes, I., Gilbert, M.T.P., and Dalén, L. (2018). Quantifying Temporal Genomic Erosion in Endangered Species. Trends in Ecology & Evolution 33, 176–185.

16. Peart, C.R., Tusso, S., Pophaly, S.D., Botero-Castro, F., Wu, C.-C., Aurioles-Gamboa, D., Baird, A.B., Bickham, J.W., Forcada, J., Galimberti, F., et al. (2020). Determinants of genetic variation across eco-evolutionary scales in pinnipeds. Nature Ecology & Evolution 4, 1095–1104.

17. Loire, E., Chiari, Y., Bernard, A., Cahais, V., Romiguier, J., Nabholz, B., Lourenço, J.M., and Galtier, N. (2013). Population genomics of the endangered giant Galápagos tortoise. Genome Biology 14, R136.

18. Rogers, R.L., and Slatkin, M. (2017). Excess of genomic defects in a woolly mammoth on Wrangel island. PLOS Genetics 13, e1006601.

19. Robinson, J.A., Brown, C., Kim, B.Y., Lohmueller, K.E., and Wayne, R.K. (2018). Purging of Strongly Deleterious Mutations Explains Long-Term Persistence and Absence of Inbreeding Depression in Island Foxes. Current Biology 28, 3487-3494.e4.

20. Robinson, J.A., Ortega-Del Vecchyo, D., Fan, Z., Kim, B.Y., vonHoldt, B.M., Marsden, C.D., Lohmueller, K.E., and Wayne, R.K. (2016). Genomic Flatlining in the Endangered Island Fox. Current Biology 26, 1183–1189.

21. Kutschera, V.E., Poelstra, J.W., Botero-Castro, F., Dussex, N., Gemmell, N.J., Hunt, G.R., Ritchie, M.G., Rutz, C., Wiberg, R.A.W., and Wolf, J.B.W. (2019). Purifying Selection in Corvids Is Less Efficient on Islands. Molecular Biology and Evolution 37, 469–474.

22. Hamabata, T., Kinoshita, G., Kurita, K., Cao, P.-L., Ito, M., Murata, J., Komaki, Y., Isagi, Y., and Makino, T. (2019). Endangered island endemic plants have vulnerable genomes. Communications Biology 2, 244.

23. Kimura, M. (1968). Evolutionary Rate at the Molecular Level. Nature 217, 624–626.

24. King, J.L., and Jukes, T.H. (1969). Non-Darwinian Evolution. Science 164, 788.

25. Chen, J., Glémin, S., and Lascoux, M. (2017). Genetic Diversity and the Efficacy of Purifying Selection across Plant and Animal Species. Molecular Biology and Evolution 34, 1417–1428.

26. Plomion, C., Aury, J.-M., Amselem, J., Leroy, T., Murat, F., Duplessis, S., Faye, S., Francillonne, N., Labadie, K., Le Provost, G., et al. (2018). Oak genome reveals facets of long lifespan. Nature Plants 4, 440–452.

27. Romiguier, J., Gayral, P., Ballenghien, M., Bernard, A., Cahais, V., Chenuil, A., Chiari, Y., Dernat, R., Duret, L., Faivre, N., et al. (2014). Comparative population genomics in animals uncovers the determinants of genetic diversity. Nature 515, 261–263.

28. Doyle, J.M., Hacking, C.C., Willoughby, J.R., Sundaram, M., and DeWoody, J.A. (2015). Mammalian genetic diversity as a function of habitat, body size, trophic class, and conservation status. Journal of Mammalogy 96, 564–572.

29. Willoughby, J.R., Sundaram, M., Wijayawardena, B.K., Kimble, S.J.A., Ji, Y., Fernandez, N.B., Antonides, J.D., Lamb, M.C., Marra, N.J., and DeWoody, J.A. (2015). The reduction of genetic diversity in threatened vertebrates and new recommendations regarding IUCN conservation rankings. Biological Conservation 191, 495–503.

30. Brüniche-Olsen, A., Kellner, K.F., and DeWoody, J.A. (2019). Island area, body size and demographic history shape genomic diversity in Darwin’s finches and related tanagers. Molecular Ecology 28, 4914–4925.

31. Brüniche-Olsen, A., Kellner, K.F., Anderson, C.J., and DeWoody, J.A. (2018). Runs of homozygosity have utility in mammalian conservation and evolutionary studies. Conservation Genetics 19, 1295– 1307.

32. Kondrashov, A.S. (1988). Deleterious mutations and the evolution of sexual reproduction. Nature 336, 435–440.

33. Lynch, M., and Gabriel, W. (1990). Mutation load and the survival of small populations. Evolution 44, 1725–1737.

34. Warren, B.H., Hagen, O., Gerber, F., Thébaud, C., Paradis, E., and Conti, E. (2018). Evaluating alternative explanations for an association of extinction risk and evolutionary uniqueness in multiple insular lineages. Evolution 72, 2005–2024.

35. Valente, L., Phillimore, A.B., Melo, M., Warren, B.H., Clegg, S.M., Havenstein, K., Tiedemann, R., Illera, J.C., Thébaud, C., Aschenbach, T., et al. (2020). A simple dynamic model explains the diversity of island birds worldwide. Nature 579, 92–96

36. Singhal, S., Leffler, E.M., Sannareddy, K., Turner, I., Venn, O., Hooper, D.M., Strand, A.I., Li, Q., Raney, B., Balakrishnan, C.N., et al. (2015). Stable recombination hotspots in birds. Science 350, 928.

37. Corcoran, P., Gossmann, T.I., Barton, H.J., Great Tit HapMap Consortium, Slate, J., and Zeng, K. (2017). Determinants of the Efficacy of Natural Selection on Coding and Noncoding Variability in Two Passerine Species. Genome Biol Evol 9, 2987–3007.

38. Lundberg, M., Liedvogel, M., Larson, K., Sigeman, H., Grahn, M., Wright, A., Åkesson, S., and Bensch, S. (2017). Genetic differences between willow warbler migratory phenotypes are few and cluster in large haplotype blocks. Evolution Letters 1, 155– 168.

39. Lamichhaney, S., Berglund, J., Almén, M.S., Maqbool, K., Grabherr, M., Martinez-Barrio, A., Promerová, M., Rubin, C.-J., Wang, C., Zamani, N., et al. (2015). Evolution of Darwin’s finches and their beaks revealed by genome sequencing. Nature 518, 371–375.

40. Burri, R., Nater, A., Kawakami, T., Mugal, C.F., Olason, P.I., Smeds, L., Suh, A., Dutoit, L., Bureš, S., Garamszegi, L.Z., et al. (2015). Linked selection and recombination rate variation drive the evolution of the genomic landscape of differentiation across the speciation continuum of Ficedula flycatchers. Genome Research. gr.196485.115

41. Bourgeois, Y.X.C., Delahaie, B., Gautier, M., Lhuillier, E., Malé, P.-J.G., Bertrand, J.A.M., Cornuault, J., Wakamatsu, K., Bouchez, O., Mould, C., et al. A novel locus on chromosome 1 underlies the evolution of a melanic plumage polymorphism in a wild songbird. Royal Society Open Science 4, 160805.

42. Leroy, T., Anselmetti, Y., Tilak, M.-K., Bérard, S., Csukonyi, L., Gabrielli, M., Scornavacca, C., Milá, B., Thébaud, C., and Nabholz, B. (2019). A bird’s white-eye view on neosex chromosome evolution. BioRxiv, 505610, ver. 4 peer-reviewed and recommended by Peer Community in Evolutionary Biology.

43. Chapman, J.A., Ho, I., Sunkara, S., Luo, S., Schroth, G.P., and Rokhsar, D.S. (2011). Meraculous: De Novo Genome Assembly with Short Paired-End Reads. PLOS ONE 6, e23501.

44. Putnam, N.H., O’Connell, B.L., Stites, J.C., Rice, B.J., Blanchette, M., Calef, R., Troll, C.J., Fields, A., Hartley, P.D., Sugnet, C.W., et al. (2016). Chromosome-scale shotgun assembly using an in vitro method for long-range linkage. Genome Research 26, 342–350.

45. Bolger, A.M., Lohse, M., and Usadel, B. (2014). Trimmomatic: a flexible trimmer for Illumina sequence data. Bioinformatics 30, 2114–2120.

46. Li, H. (2013). Aligning sequence reads, clone sequences and assembly contigs with BWA-MEM. arXiv.

47. Broad Institute (2019). Picard Toolkit (http://broadinstitute.github.io/picard/)

48. McKenna, A., Hanna, M., Banks, E., Sivachenko, A., Cibulskis, K., Kernytsky, A., Garimella, K., Altshuler, D., Gabriel, S., Daly, M., et al. (2010). The Genome Analysis Toolkit: a MapReduce framework for analyzing next-generation DNA sequencing data. Genome Res 20, 1297–1303.

49. Allio, R., Schomaker-Bastos, A., Romiguier, J., Prosdocimi, F., Nabholz, B., and Delsuc, F. (in press). MitoFinder: efficient automated large-scale extraction of mitogenomic data in target enrichment phylogenomics. Mol Ecol Resour.

50. Ranwez, V., Douzery, E.J.P., Cambon, C., Chantret, N., and Delsuc, F. (2018). MACSE v2: Toolkit for the Alignment of Coding Sequences Accounting for Frameshifts and Stop Codons. Molecular Biology and Evolution 35, 2582–2584.

51. Nguyen, L.-T., Schmidt, H.A., von Haeseler, A., and Minh, B.Q. (2015). IQ-TREE: A Fast and Effective Stochastic Algorithm for Estimating Maximum-Likelihood Phylogenies. Molecular Biology and Evolution 32, 268–274.

52. She, R., Chu, J.S.-C., Uyar, B., Wang, J., Wang, K., and Chen, N. (2011). genBlastG: using BLAST searches to build homologous gene models. Bioinformatics 27, 2141–2143.

53. Eddy, S.R. (2011). Accelerated Profile HMM Searches. PLoS Comput Biol 7, e1002195–e1002195.

54. Li, H., and Durbin, R. (2011). Inference of human population history from individual whole-genome sequences. Nature 475, 493– 496.

55. R Core Team (2018). R: A Language and Environment for Statistical Computing (Vienna, Austria: R Foundation for Statistical Computing)

56. Lamichhaney, S., Han, F., Berglund, J., Wang, C., Almén, M.S., Webster, M.T., Grant, B.R., Grant, P.R., and Andersson, L. (2016). A beak size locus in Darwin’s finches facilitated character displacement during a drought. Science 352, 470.

57. Zink, R.M., and Vázquez-Miranda, H. (2018). Species Limits and Phylogenomic Relationships of Darwin’s Finches Remain Unresolved: Potential Consequences of a Volatile Ecological Setting. Systematic Biology 68, 347–357.

58. Parker, P., Li, B., Li, H., and Wang, J. (2012). The genome of Darwin’s Finch (Geospiza fortis). GigaScience.

59. Ellegren, H., Smeds, L., Burri, R., Olason, P.I., Backström, N., Kawakami, T., Künstner, A., Mäkinen, H., Nadachowska-Brzyska, K., Qvarnström, A., et al. (2012). The genomic landscape of species divergence in Ficedula flycatchers. Nature 491, 756.

60. Warren, W.C., Clayton, D.F., Ellegren, H., Arnold, A.P., Hillier, L.W., Künstner, A., Searle, S., White, S., Vilella, A.J., Fairley, S., et al. (2010). The genome of a songbird. Nature 464, 757.

61. Laine, V.N., Gossmann, T.I., Schachtschneider, K.M., Garroway, C.J., Madsen, O., Verhoeven, K.J.F., de Jager, V., Megens, H.-J., Warren, W.C., Minx, P., et al. (2016). Evolutionary signals of selection on cognition from the great tit genome and methylome. Nature Communications 7, 10474.

62. Recuerda, M., Vizueta, J., Cuevas-Caballé, C., Blanco, G., Rozas, J., and Milá, B. (2020). Chromosome-level genome assembly of the common chaffinch (Aves: Fringilla coelebs): a valuable resource for evolutionary biology. bioRxiv, 2020.11.30.404061.

63. Scornavacca, C., Belkhir, K., Lopez, J., Dernat, R., Delsuc, F., Douzery, E.J.P., and Ranwez, V. (2019). OrthoMaM v10: Scaling-Up Orthologous Coding Sequence and Exon Alignments with More than One Hundred Mammalian Genomes. Molecular Biology and Evolution 36, 861–862.

64. Guéguen, L., Gaillard, S., Boussau, B., Gouy, M., Groussin, M., Rochette, N.C., Bigot, T., Fournier, D., Pouyet, F., Cahais, V., et al. (2013). Bio++: efficient extensible libraries and tools for computational molecular evolution. Mol Biol Evol 30, 1745– 1750.

65. Nadachowska-Brzyska, K., Burri, R., Olason, P.I., Kawakami, T., Smeds, L., and Ellegren, H. (2013). Demographic Divergence History of Pied Flycatcher and Collared Flycatcher Inferred from Whole-Genome Re-sequencing Data. PLOS Genetics 9, e1003942.

66. Kim, S., Cho, Y.S., Kim, H.-M., Chung, O., Kim, H., Jho, S., Seomun, H., Kim, J., Bang, W.Y., Kim, C., et al. (2016). Comparison of carnivore, omnivore, and herbivore mammalian genomes with a new leopard assembly. Genome Biology 17, 211.

67. Smeds, L., Qvarnström, A., and Ellegren, H. (2016). Direct estimate of the rate of germline mutation in a bird. Genome Res 26, 1211–1218.

68. Brommer, J.E., Gustafsson, L., Pietiäinen, H., and Merilä, J. (2004). Single-generation estimates of individual fitness as proxies for long-term genetic contribution. Am Nat 163, 505–517.

69. Liu, S., and Hansen, M.M. (2017). PSMC (pairwise sequentially Markovian coalescent) analysis of RAD (restriction site associated DNA) sequencing data. Molecular Ecology Resources 17, 631–641.

70. Wickham, H. (2016). ggplot2: Elegant Graphics for Data Analysis (Springer-Verlag New York)

71. Wilke, C. (2016). cowplot: Streamlined Plot Theme and Plot Annotations for “ggplot2”

72. Bolívar, P., Mugal, C.F., Nater, A., and Ellegren, H. (2016). Recombination Rate Variation Modulates Gene Sequence Evolution Mainly via GC-Biased Gene Conversion, Not Hill-Robertson Interference, in an Avian System. Mol Biol Evol 33, 216– 227.

73. Eyre-Walker, A., and Keightley, P.D. (2009). Estimating the Rate of Adaptive Molecular Evolution in the Presence of Slightly Deleterious Mutations and Population Size Change. Molecular Biology and Evolution 26, 2097–2108.

74. Eyre-Walker, A., Woolfit, M., and Phelps, T. (2006). The distribution of fitness effects of new deleterious amino acid mutations in humans. Genetics 173, 891–900.

75. Pinheiro, J., Bates, D., DebRoy, S., Sarkar, D., and R Core Team (2020). nlme: Linear and Nonlinear Mixed Effects Models

76. Paradis, E., and Schliep, K. (2018). ape 5.0: an environment for modern phylogenetics and evolutionary analyses in R. Bioinformatics 35, 526–528.

77. Kassambara, A. (2018). “ggplot2” Based Publication Ready Plots

78. Slowikowski, K. (2019). ggrepel: Automatically Position Non-Overlapping Text Labels with “ggplot2”

79. Yu, G., Smith, D.K., Zhu, H., Guan, Y., and Lam, T.T.-Y. (2017). ggtree: an r package for visualization and annotation of phylogenetic trees with their covariates and other associated data. Methods in Ecology and Evolution 8, 28–36.

