## Supplementary_Information for "Island songbirds as windows into evolution in small populations"

#### Table of contents:

Supplementary Tables S1-S3 — 2

Supplementary Figures (S1-S4) — 6

Supplementary Note S1 (Extended Introduction) — 17

Supplementary Note S2 (Within-genome variation in the efficacy of purifying selection) — 21

Supplementary Methods S1 (Extended Materials & Methods) — 26

References (SI only) — 30

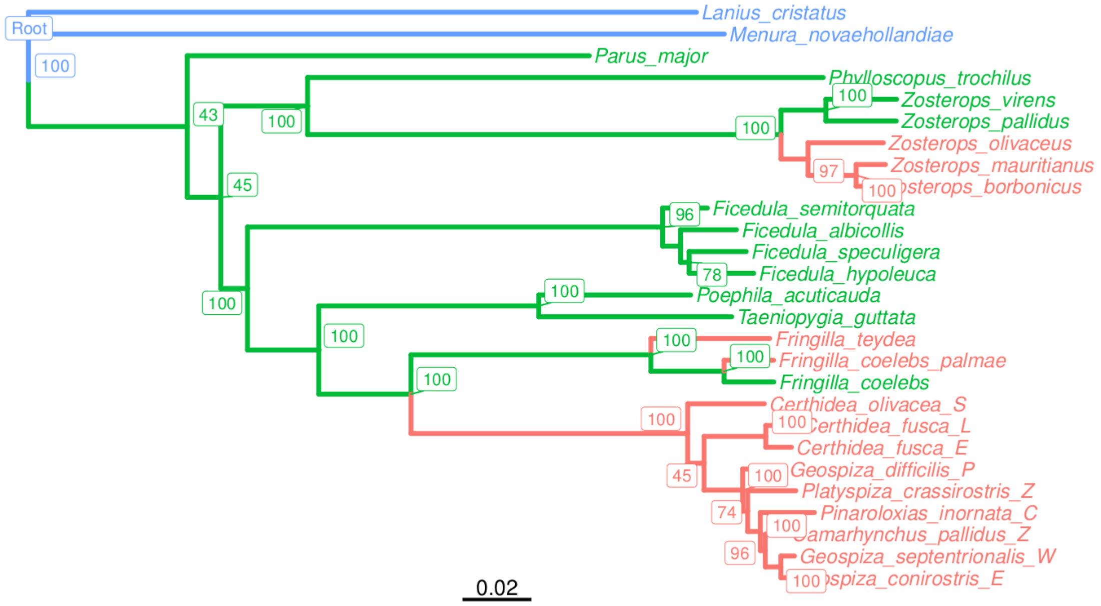


***Fig. S1: Phylogenetic relationship estimated using complete mitochondrial sequences and a maximum likelihood approach.*** *Island, continental and outgroup species are shown in red, green and blue, respectively. Numbers shown in boxes indicated ultrafast bootstrap supports estimated by IQTREE.*

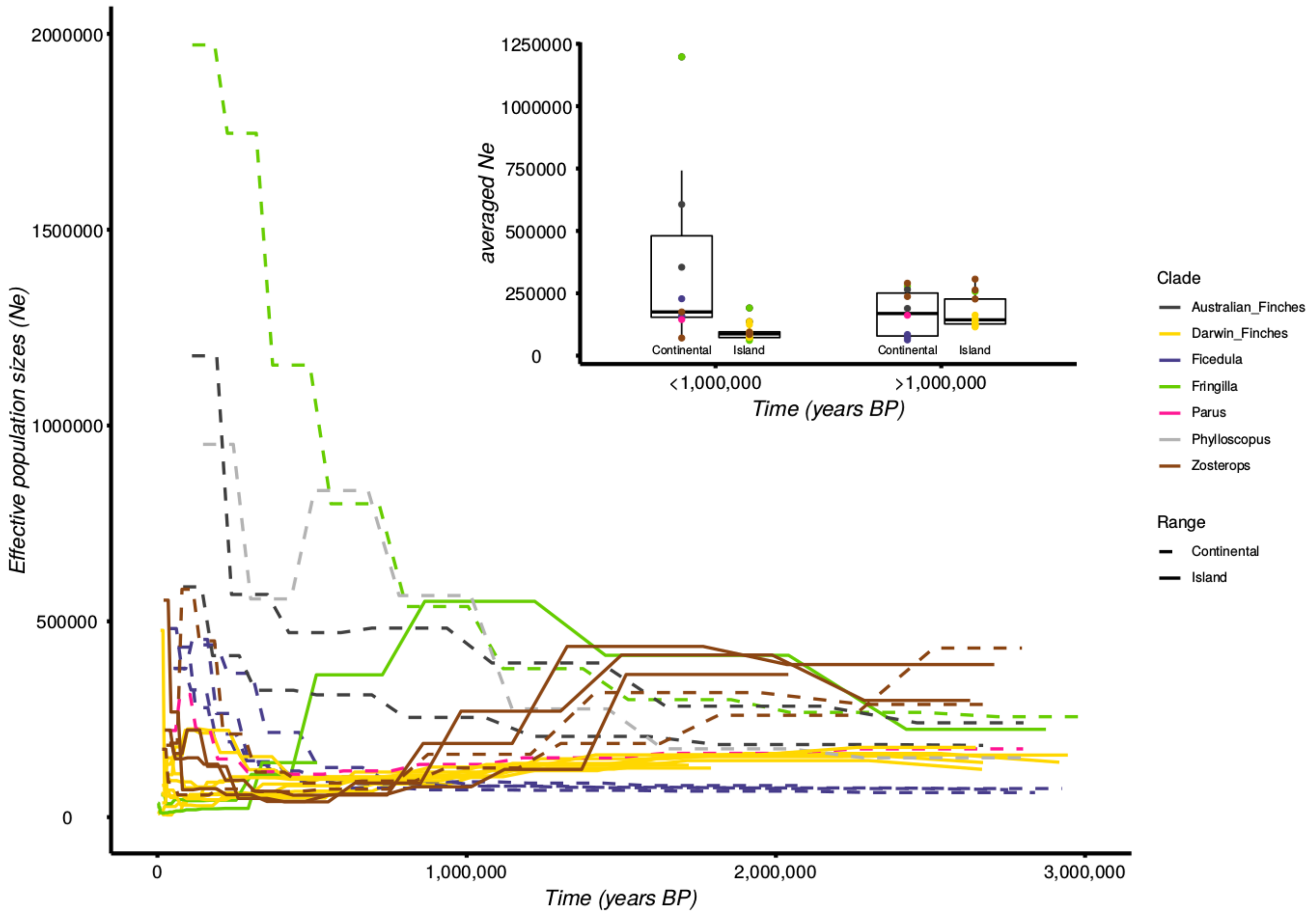


***Fig. S2: PSMC estimates of the changes in effective population size (Ne) over time for each species (outer plot).*** *Each line represents the historical changes of Ne for one individual randomly selected among all individuals of each species. The inner boxplot shows the averaged mean Ne for all island and continental species over the last million years or estimates for older times. Estimates for the last million year exclude the last 4 estimates (see the STAR Methods), because PSMC can generate inaccurate estimates for present-day or very recent Ne.*

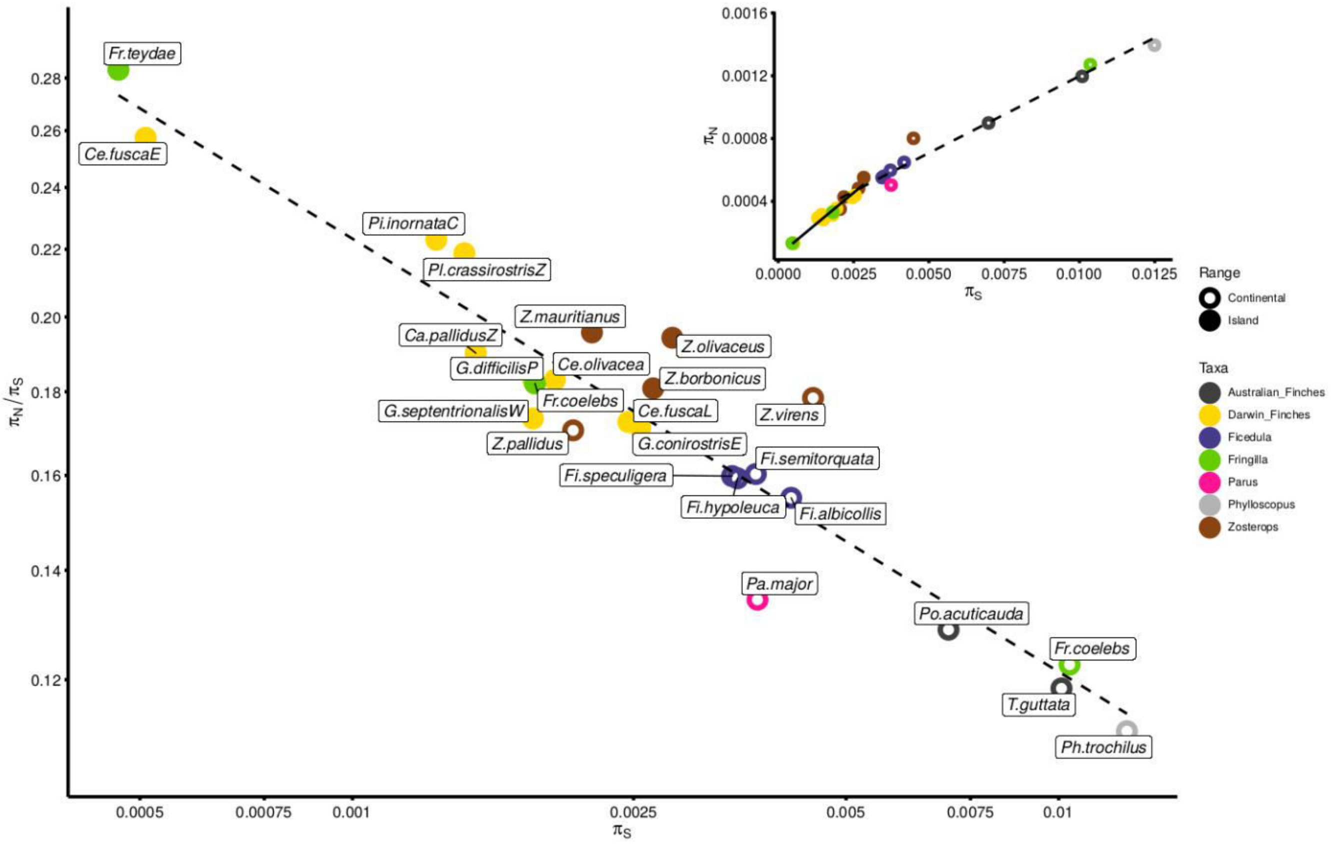


***Fig. S3: Linear regression between the log-transformed ratio of nonsynonymous to synonymous nucleotide diversity ($\pi_N/\pi_S$) and the log-transformed levels of nucleotide diversity ($\pi_S$), used as an indicator of effective population sizes (outer plot)***. *Species genera are: Ca=Camarhynchus, Ce=Certhidea, Fi=Ficedula, Fr=Fringilla, G=Geospiza, Pa=Parus, Ph=Phylloscopus, Pi=Pinaroloxias, Pl=Platyspiza, Po= Poephila, T=Taeniopygia, Z= Zosterops. Inner plot: linear regressions between $\pi_N$ and $\pi_S$.for insular species (solid line) vs. continental species (dotted line).*

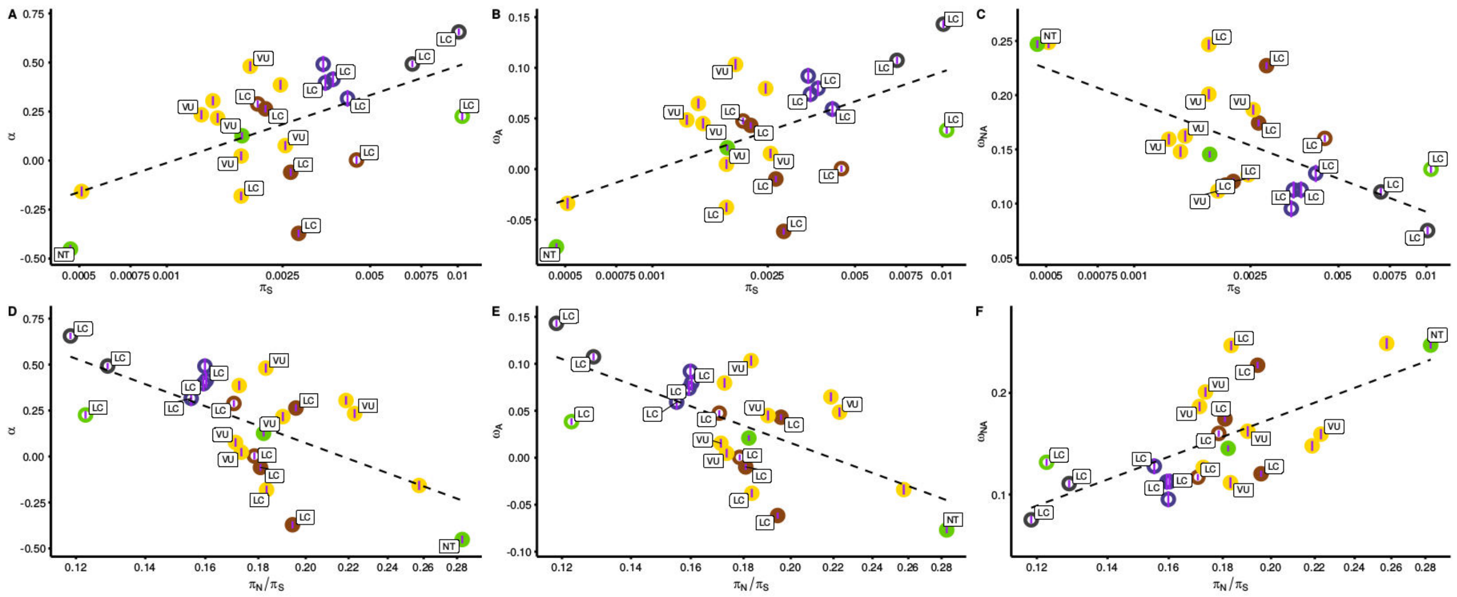

***Fig S4: Scatterplots of proportion of amino-acid substitutions that result from positive selection (α, A and C), adaptive ratio of nonsynonymous over synonymous ($\omega_A$, B and E) & non-adaptive ratio of non-synonymous over synonymous substitutions ($\omega_{NA}$, C and F) with regards to the levels of nucleotide diversity ($\pi_S$; A, B and C) and the observed ratios of nonsynonymous to synonymous mutations in the polymorphism data ($\pi_N/\pi_S$; D, E and F).***

***Table S1: Linear models and* Phylogenetic Generalized Least Squared (*PGLS) method-based tests performed.*** *To perform accurate comparisons, model selections were performed using the maximum number of species (N) for which all information was available (i.e. 13 species with census size estimates, 21 species with IUCN status, and 25 with both insularity status & range sizes). IUCN status was considered as a binary variable "Threatened", when the status is "Vulnerable", or "Non-threatened" otherwise (i.e. "Least Concerned or Near-threatened). For each comparison, the best model is indicated in bold. For the models with PGLS, the p-value of the model was obtained from a Likelihood ratio test using the anova.lme function (nlme package, Pinheiro et al. 2020).* ▲*: for these tests, assumptions of normality, heteroscedasticity and independence are not all respected (Shapiro, Breusch-Pagan and Durbin-Watson tests)*

| Dependent variable | Explanatory variable | Test | Model | N | $R^2$ | p-val model* | p-value variable 1 | p-value variable 2 | AICc | DeltaAICc |
|---|---|---|---|---|---|---|---|---|---|---|
| $\pi_S$ | Insularity *vs.* census size | Linear models | ps~Insularity | 13 | 0.448 | 0.012 | -- | -- | -109.83 | 3.44 |
| | | | **ps~census_log** | **13** | **0.576** | **0.003** | -- | -- | **-113.27** | **0** |
| | | | ps~census_log+Insularity▲ | 13 | 0.578 | 0.013 | 0.109 | 0.829 | -109.00 | 4.27 |
| | | PGLS | **ps~Insularity** | **13** | **--** | **1.06e$^{-06}$** | **--** | **--** | **-120.13** | **0** |
| | | | ps~census_log | 13 | -- | 0.001 | -- | -- | -106.63 | 13.51 |
| | | | ps~census_log+Insularity | 13 | -- | 9.70e$^{-07}$ | 0.73 | 0.002 | -115.96 | 4.17 |
| $\log_{10}(\pi_S)$ | Insularity *vs.* census size | Linear models | ps_log~Insularity | 13 | 0.546 | 0.003 | -- | -- | 7.67 | 1.72 |
| | | | **ps_log~census_log** | **13** | **0.63** | **0.001** | -- | -- | **5.95** | **0** |
| | | | ps_log~census_log+Insularity | 13 | 0.656 | 0.005 | 0.162 | 0.405 | 9.34 | 3.38 |
| | | PGLS | **ps_log~Insularity** | **13** | -- | 1.00e$^{-04}$ | -- | -- | **10.66** | **0** |
| | | | ps_log~census_log | 13 | -- | 0.015 | -- | -- | 19.91 | 9.25 |
| | | | ps_log~census_log+Insularity | 13 | -- | 3.00e$^{-04}$ | 0.365 | 0.006 | 13.87 | 3.21 |
| $\pi_S$ | Insularity *vs.* species range area | Linear models | ps ~ Insularity | 25 | 0.445 | 2.73e$^{-04}$ | -- | -- | -226.10 | 2.02 |
| | | | **ps ~ range_kmsq_log** | **25** | **0.488** | **1.04e$^{-04}$** | -- | -- | **-228.12** | **0** |
| | | | ps ~ Insularity + range_kmsq_log▲ | 25 | 0.49 | 6.06e$^{-04}$ | 0.175 | 0.751 | -225.38 | 2.74 |
| | | PGLS | **ps ~ Insularity** | **21** | **--** | **3.54e$^{-05}$** | -- | -- | -234.07 | 0.06 |
| | | | ps ~ range_kmsq_log | 21 | -- | 8.28e$^{-05}$ | -- | -- | -232.46 | 1.67 |
| | | | ps ~ Insularity + range_kmsq_log | 21 | -- | 4.50e$^{-05}$ | 0.049 | 0.113 | **-234.13** | **0** |
| $\log_{10}(\pi_S)$ | Insularity *vs.* species range | Linear models | ps_log ~ Insularity | 25 | 0.528 | 3.94e$^{-05}$ | -- | -- | 5.59 | 0.52 |
| | | | **ps_log ~ range_kmsq_log** | **25** | **0.537** | **3.08e$^{-05}$** | -- | -- | **5.07** | **0** |

| | | | | | | | | | | |
|---|---|---|---|---|---|---|---|---|---|---|
| | area | | ps_log ~ Insularity + range_kmsq_log | 25 | 0.552 | $1.45e^{-04}$ | 0.282 | 0.398 | 7.10 | 2.03 |
| | | PGLS | ps_log ~ Insularity | 25 | -- | 0.005 | -- | -- | 21.52 | 2.15 |
| | | | **ps_log ~ range_kmsq_log** | **25** | -- | **0.002** | -- | -- | **19.36** | **0** |
| | | | ps_log ~ range_kmsq_log+Insularity | 25 | -- | 0.005 | 0.43 | 0.116 | 21.50 | 2.13 |
| $\pi_S$ | Insularity *vs.* IUCN | Linear models | **ps~Insularity** | **21** | **0.444** | **0.001** | -- | -- | **-186.33** | **0** |
| | | | ps~IUCN | 21 | 0.106 | 0.132 | -- | -- | -176.96 | 9.37 |
| | | | ps~IUCN+Insularity▲ | 21 | 0.444 | 0.005 | 0.96 | 0.005 | -183.24 | 3.09 |
| | | PGLS | **ps~Insularity** | **25** | -- | **0.004** | -- | -- | **-194.88** | **0** |
| | | | ps~IUCN | 25 | -- | 0.926 | -- | -- | -186.62 | 8.26 |
| | | | ps~IUCN+Insularity | 25 | -- | 0.016 | 0.938 | 0.009 | -191.80 | 3.08 |
| $\log_{10}(\pi_S)$ | Insularity *vs.* IUCN | Linear models | **ps_log~Insularity** | **21** | **0.547** | **$1.28e^{-04}$** | -- | -- | **4.13** | **0** |
| | | | ps_log~IUCN | 21 | 0.133 | 0.104 | -- | -- | 17.75 | 13.62 |
| | | | ps_log~IUCN+Insularity | 21 | 0.548 | 0.001 | 0.83 | 0.001 | 7.16 | 3.03 |
| | | PGLS | **ps_log~Insularity** | **21** | -- | **0.007** | -- | -- | **10.83** | **0** |
| | | | ps_log~IUCN | 21 | -- | 0.942 | -- | -- | 18.21 | 7.38 |
| | | | ps_log~IUCN+Insularity | 21 | -- | 0.024 | 0.804 | 0.013 | 13.85 | 3.01 |
| $\pi_N/\pi_S$ | Insularity *vs.* census size | Linear models | pnps~Insularity | 13 | 0.514 | 0.006 | -- | -- | -45.84 | 1.76 |
| | | | **pnps~census_log** | **13** | **0.576** | **0.003** | -- | -- | **-47.60** | **0** |
| | | | pnps~census_log+Insularity | 13 | 0.595 | 0.011 | 0.19 | 0.512 | -43.85 | 3.75 |
| | | PGLS | **pnps~Insularity** | **13** | -- | **0.003** | -- | -- | **-39.44** | **0** |
| | | | pnps~census_log | 13 | -- | 0.047 | -- | -- | -34.84 | 4.60 |
| | | | pnps~census_log+Insularity | 13 | -- | 0.011 | 0.599 | 0.056 | -35.48 | 3.96 |
| $\log_{10}(\pi_N/\pi_S)$ | Insularity *vs.* census size | Linear models | pnps_log~Insularity | 13 | 0.577 | 0.003 | -- | -- | -25.01 | 3.02 |
| | | | **pnps_log~census_log** | **13** | **0.665** | **$6.80e^{-04}$** | -- | -- | **-28.03** | **0** |
| | | | pnps_log~census_log+Insularity | 13 | 0.681 | 0.003 | 0.102 | 0.496 | 24.33 | 52.36 |
| | | PGLS | **pnps_log~Insularity** | **13** | -- | **$5.72e^{-04}$** | -- | -- | **-20.26** | **0** |
| | | | pnps_log~census_log | 13 | -- | 0.021 | -- | -- | -13.72 | 6.54 |
| | | | pnps_log~census_log+Insularity | 13 | -- | 0.002 | 0.557 | 0.023 | -16.40 | 3.86 |

| | | | | | | | | | | |
|---|---|---|---|---|---|---|---|---|---|---|
| $\pi_N/\pi_S$ | Insularity *vs.* species range area | Linear models | **pnps~Insularity** | **25** | **0.484** | **$1.13e^{-04}$** | -- | -- | **-100.15** | **0** |
| | | | pnps~range_kmsq_log | 25 | 0.451 | $2.38e^{-04}$ | -- | -- | -98.59 | 1.55 |
| | | | pnps~Insularity+range_kmsq_log▲ | 25 | 0.489 | $6.24e^{-04}$ | 0.658 | 0.216 | -97.52 | 2.63 |
| | | PGLS | **pnps~Insularity** | **21** | -- | **0.01683009** | -- | -- | **-83.79** | **0** |
| | | | pnps~range_kmsq_log | 21 | -- | 0.02243272 | -- | -- | -83.28 | 0.50 |
| | | | pnps~Insularity+range_kmsq_log | 21 | -- | 0.04064072 | 0.311 | 0.44 | -81.62 | 2.17 |
| $\log_{10}(\pi_N/\pi_S)$ | Insularity *vs.* species range area | Linear models | **pnps_log~Insularity** | **25** | **0.529** | **$3.76e^{-05}$** | -- | -- | **-58.25** | **0** |
| | | | pnps_log~range_kmsq_log | 25 | 0.513 | 0.5126 | -- | -- | -57.37 | 0.88 |
| | | | pnps_log~Insularity+range_kmsq_log | 25 | 0.541 | $1.90e^{-04}$ | 0.461 | 0.254 | -56.03 | 2.23 |
| | | PGLS | **pnps_log ~ Insularity** | **25** | -- | **0.005** | -- | -- | **-45.82** | **0** |
| | | | pnps_log ~ range_kmsq_log | 25 | -- | 0.009 | -- | -- | -44.61 | 1.20 |
| | | | pnps_log ~ range_kmsq_log + Insularity | 25 | -- | 0.012 | 0.191 | 0.41 | -43.75 | 2.07 |
| $\pi_N/\pi_S$ | Insularity *vs.* IUCN | Linear models | **pnps~Insularity** | **21** | **0.513** | **$2.59e^{-04}$** | -- | -- | **-84.52** | **0** |
| | | | pnps~IUCN | 21 | 0.349 | 0.04632 | -- | -- | -70.39 | 14.13 |
| | | | pnps~IUCN+Insularity▲ | 21 | 0.552 | $7.02e^{-04}$ | 0.228 | 0 | -83.17 | 1.34 |
| | | PGLS | **pnps~Insularity** | **25** | -- | **0.0037** | -- | -- | **-78.90** | **0** |
| | | | pnps~IUCN | 25 | -- | 0.6279 | -- | -- | -70.69 | 8.21 |
| | | | pnps~IUCN+Insularity | 25 | -- | 0.0107 | 0.466 | 0.007 | -76.45 | 2.45 |
| $\log_{10}(\pi_N/\pi_S)$ | Insularity *vs.* IUCN | Linear models | **pnps_log~Insularity** | **21** | **0.552** | **$1.13e^{-04}$** | -- | -- | **-48.26** | **0** |
| | | | pnps_log~IUCN | 21 | 0.067 | 0.2559 | -- | -- | -32.84 | 15.41 |
| | | | pnps_log~IUCN+Insularity | 21 | 0.5784 | $4.21e^{-04}$ | 0.305 | 0 | -46.43 | 1.83 |
| | | PGLS | **pnps_log~Insularity** | **21** | -- | **0.0027** | -- | -- | **-44.56** | **0** |
| | | | pnps_log~IUCN | 21 | -- | 0.6671 | -- | -- | -35.76 | 8.80 |
| | | | pnps_log~IUCN+Insularity | 21 | -- | 0.0085 | 0.496 | 0.005 | -42.03 | 2.53 |
| $\omega_A$ | all | Linear models | omega_A ~ Insularity | 23 | 0.255 | 0.014 | -- | -- | -67.91 | 5.62 |
| | | | omega_A ~ ps | 23 | 0.234 | 0.020 | -- | -- | -67.25 | 6.29 |
| | | | omega_A ~ pnps | 23 | 0.416 | $8.88\ e^{-04}$ | -- | -- | -73.51 | 0.025 |

| | | | | | | | | | | |
|---|---|---|---|---|---|---|---|---|---|---|
| | | | omega_A ~ ps_log | 23 | 0.332 | 0.004 | -- | -- | -70.40 | 3.13 |
| | | | **omega_A ~ pnps_log** | **23** | **0.417** | **$8.80\ e^{-04}$** | -- | -- | **-73.54** | **0** |
| | | | omega_A ~ PSMC_Ne | 23 | 0.165 | 0.054 | -- | -- | -65.29 | 8.25 |
| | | | omega_A ~ range_kmsq_log | 23 | 0.212 | 0.027 | -- | -- | -66.60 | 6.94 |
| | | PGLS | omega_A ~ Insularity | 23 | -- | 0.237 | -- | -- | -56.42 | 3.74 |
| | | | omega_A ~ ps | 23 | -- | 0.220 | -- | -- | -56.52 | 3.64 |
| | | | **omega_A ~ pnps** | **23** | -- | **0.023** | -- | -- | **-60.16** | **0** |
| | | | omega_A ~ ps_log | 23 | -- | 0.029 | -- | -- | -59.76 | 0.39 |
| | | | omega_A ~ pnps_log | 23 | -- | 0.034 | -- | -- | -59.50 | 0.65 |
| | | | omega_A ~ PSMC_Ne | 23 | -- | 0.590 | -- | -- | -55.31 | 4.85 |
| | | | omega_A ~ range_kmsq_log | 23 | -- | 0.054 | -- | -- | -58.72 | 1.43 |
| $\omega_{NA}$ | all | Linear models | omega_NA ~ Insularity | 23 | 0.384 | 0.002 | -- | -- | -76.74 | 3.64 |
| | | | omega_NA ~ ps | 23 | 0.279 | 0.010 | -- | -- | -73.12 | 7.26 |
| | | | **omega_NA ~ pnps** | **23** | **0.474** | **$2.28e^{-04}$** | -- | -- | **-80.39** | **0** |
| | | | omega_NA ~ ps_log | 23 | 0.44 | $5.56e^{-04}$ | -- | -- | -78.96 | 1.43 |
| | | | omega_NA ~ pnps_log | 23 | 0.463 | $3.54e^{-04}$ | -- | -- | -79.89 | 0.49 |
| | | | omega_NA ~ PSMC_Ne | 23 | 0.213 | 0.027 | -- | -- | -71.10 | 9.29 |
| | | | omega_NA ~ range_kmsq_log | 23 | 0.387 | 0.002 | -- | -- | -76.87 | 3.51 |
| | | PGLS | omega_NA ~ Insularity | 23 | -- | 0.257 | -- | -- | -56.08 | 4.53 |
| | | | omega_NA ~ ps | 23 | -- | 0.214 | -- | -- | -56.34 | 4.27 |
| | | | **omega_NA ~ pnps** | **23** | -- | **0.016** | -- | -- | **-60.61** | **0** |
| | | | omega_NA ~ ps_log | 23 | -- | 0.020 | -- | -- | -60.24 | 0.37 |
| | | | omega_NA ~ pnps_log | 23 | -- | 0.025 | -- | -- | -59.80 | 0.81 |
| | | | omega_NA ~ PSMC_Ne | 23 | -- | 0.550 | -- | -- | -55.31 | 5.30 |
| | | | omega_NA ~ range_kmsq_log | 23 | -- | 0.054 | -- | -- | -58.50 | 2.11 |

***Table S2: $\pi_N$, $\pi_S$ and $\pi_N/\pi_S$ as estimated by the diversity-based (meth.1) and the SFS-based (meth.2) methods.*** *For the method 2, P. major and P. trochilus were not excluded for the computations (no phylogenetically relevant outgroup information available).*

| **Groups** | **species** | **outgroup** | **Range** | **πS (meth1)** | **πN (meth1)** | **πN/πS (meth1)** | **πS (meth2)** | **πN (meth2)** | **πN/πS (meth2)** |
|---|---|---|---|---|---|---|---|---|---|
| *Zosterops* | *Z.borbonicus* | *Z.pallidus* | Island | 0.00267 | 0.00048 | 0.18086 | 0.00049 | 0.00239 | 0.20353 |
| *Zosterops* | *Z.olivaceus* | *Z.virens* | Island | 0.00284 | 0.00055 | 0.19422 | 0.00061 | 0.00268 | 0.22600 |
| *Zosterops* | *Z.mauritianus* | *Z.virens* | Island | 0.00218 | 0.00043 | 0.19569 | 0.00038 | 0.00166 | 0.22898 |
| *Zosterops* | *Z.pallidus* | *Z.borbonicus* | Continental | 0.00205 | 0.00035 | 0.17048 | 0.00029 | 0.00141 | 0.20435 |
| *Zosterops* | *Z.virens* | *Z.borbonicus* | Continental | 0.004490 | 0.00080 | 0.17844 | 0.00097 | 0.00463 | 0.20842 |
| *Ficedula* | *Fi.albicollis* | *Fi.speculigera* | Continental | 0.00418 | 0.00065 | 0.15502 | 0.00151 | 0.00830 | 0.18185 |
| *Ficedula* | *Fi.semitorquata* | *Fi.speculigera* | Continental | 0.00372 | 0.00060 | 0.16031 | 0.00269 | 0.01384 | 0.19446 |
| *Ficedula* | *Fi.hypoleuca* | *Fi.speculigera* | Continental | 0.00351 | 0.00056 | 0.15935 | 0.00175 | 0.00904 | 0.19362 |
| *Ficedula* | *Fi.speculigera* | *Fi.albicollis* | Continental | 0.00345 | 0.00055 | 0.15985 | 0.00097 | 0.00493 | 0.19631 |
| Darwin Finches | *Ce.olivacea* | *Ce.fuscaE* | Island | 0.00193 | 0.00035 | 0.18304 | 0.00027 | 0.00118 | 0.22832 |
| Darwin Finches | *G.septentrionalis W* | *Ce.olivacea* | Island | 0.00180 | 0.00031 | 0.17333 | 0.00033 | 0.00149 | 0.21897 |
| Darwin Finches | *Ce.fuscaE* | *Ce.olivacea* | Island | 0.00051 | 0.00013 | 0.25737 | 0.00019 | 0.00057 | 0.33101 |
| Darwin Finches | *Pl.crassirostrisZ* | *Ce.olivacea* | Island | 0.00144 | 0.00032 | 0.21875 | 0.00022 | 0.00079 | 0.27822 |
| Darwin Finches | *Pi.inornataC* | *Ce.olivacea* | Island | 0.00131 | 0.00029 | 0.22298 | 0.00036 | 0.00133 | 0.27235 |
| Darwin Finches | *Ce.fuscaL* | *Ce.olivacea* | Island | 0.00246 | 0.00042 | 0.17250 | 0.00070 | 0.00327 | 0.21465 |
| Darwin Finches | *Ca.pallidusZ* | *Ce.olivacea* | Island | 0.00149 | 0.00028 | 0.19009 | 0.00010 | 0.00041 | 0.23349 |
| Darwin Finches | *G.conirostrisE* | *Ce.olivacea* | Island | 0.00255 | 0.00044 | 0.17110 | 0.00041 | 0.00195 | 0.20801 |
| Darwin Finches | *G.difficilisP* | *Ce.olivacea* | Island | 0.00180 | 0.00033 | 0.18333 | 0.00038 | 0.00167 | 0.22543 |
| Parus | *Pa.major* | NA | Continental | 0.00375 | 0.00050 | 0.13431 | NA | NA | NA |
| Phylloscopus | *Ph.trochilus* | NA | Continental | 0.01249 | 0.00139 | 0.1116 | NA | NA | NA |
| Australian Finches | *T.guttata castanodis* | *Po.acuticauda* | Continental | 0.01009 | 0.00120 | 0.11853 | 0.00109 | 0.00856 | 0.12702 |
| Australian Finches | *Po.acuticauda* | *T.guttata castanodis* | Continental | 0.00698 | 0.00090 | 0.12876 | 0.00076 | 0.00526 | 0.14493 |
| Fringilla | *Fr.coelebs* | *Fr.teydae* | Continental | 0.01036 | 0.00127 | 0.12258 | 0.00119 | 0.00869 | 0.13703 |
| Fringilla | *Fr.coelebs* | *Fr.teydae* | Island | 0.00181 | 0.00033 | 0.18212 | 0.00032 | 0.00151 | 0.21456 |
| Fringilla | *Fr.teydae* | *Fr.coelebs(M)* | Island | 0.00047 | 0.00013 | 0.28326 | 0.00010 | 0.00029 | 0.34870 |

**Table S3: Complete list of all accessions used in the study.** More detailed sampling information is given for all in-house whole genome sequencing data.

| Species | Reference | SRA | code | Sampling location | Year | DNA extraction | Sequencing | Morph | sex | Longitude | Latitude | Elevation (m) | MNCN_ID |
|---|---|---|---|---|---|---|---|---|---|---|---|---|---|
| *Zosterops borbonicus* | this study | SRR12615530 | 11_0809 | Trois Citernes , Réunion | 2011 | Montpellier ISEM | Novogene | GHB | M | -21.15885 | 55.76181 | 846 | -- |
| *Zosterops borbonicus* | this study | SRR12615529 | 11_1020 | Coulée 2007, Réunion | 2012 | Montpellier ISEM | Novogene | BNB | M | -21.27819 | 55.79164 | 194 | -- |
| *Zosterops borbonicus* | this study | SRR12615520 | 264 | Basse Vallée, Réunion | 2007 | Montpellier ISEM | Novogene | BNB | M | -21.34094 | 55.70902 | 687 | -- |
| *Zosterops borbonicus* | this study | SRR12615519 | 996 | Ermitage, Réunion | 2009 | Montpellier ISEM | Novogene | LBHB | M | -21.07334 | 55.23271 | 60 | -- |
| *Zosterops borbonicus* | this study | SRR12615518 | 1602 | Piton de l'Entonnoir, | 2010 | Montpellier ISEM | Novogene | BNB | M | -21.35665 | 55.61450 | 430 | -- |
| *Zosterops borbonicus* | this study | SRR12615517 | 17-670 | Bébour, Réunion | -- | Montpellier ISEM | Novogene | HBHB | -- | -21.09650 | 55.54945 | 1540 | -- |
| *Zosterops borbonicus* | This study | SRR12615516 | 487 | Saint-Leu, Reunion | 2007 | Toulouse, LBBE | GenoToul | LBHB | M | -21.13736 | 55.29531 | 509 | -- |
| *Zosterops borbonicus* | This study | SRR12615515 | 604 | L'Etang-salé les Bains, | 2008 | Toulouse, LBBE | GenoToul | LBHB | M | -21.25848 | 55.33786 | 68 | -- |
| *Zosterops borbonicus* | This study | SRR12615514 | 1243 | Tévelave, Reunion | 2009 | Toulouse, LBBE | GenoToul | HBHB | M | -21.16891 | 55.38707 | 1962 | -- |
| *Zosterops borbonicus* | This study | SRR12615513 | 1544 | Moka, Reunion | 2010 | Toulouse, LBBE | GenoToul | GHB | M | -20.92789 | 55.51502 | 219 | -- |
| *Zosterops borbonicus* | This study | SRR12615528 | 1681 | Ravine d'Abord, Reunion | -- | Toulouse, LBBE | GenoToul | BNB | M | -21.32539 | 55.49492 | 160 | -- |
| *Zosterops borbonicus* | This study | SRR12615527 | 2062 | Rivière du Mât les Bas, | 2013 | Toulouse, LBBE | GenoToul | GHB | F | -21.97940 | 55.68230 | 42 | -- |
| *Zosterops borbonicus* | This study | SRR12615526 | 2078 | Sentier Ste-Marguerite, | 2013 | Toulouse, LBBE | GenoToul | GHB | M | -21.10876 | 55.68827 | 550 | -- |
| *Zosterops borbonicus* | This study | SRR12615525 | 17-687 | Grand Matarum, Reunion | -- | Toulouse, LBBE | GenoToul | HBHB | -- | -21.12443 | 55.47868 | 1460 | -- |
| *Zosterops borbonicus* | This study | SRR12615524 | 17-703 | Gros Galet; Reunion | -- | Toulouse, LBBE | GenoToul | HBHB | -- | -21.17890 | 55.48360 | 945 | -- |
| *Zosterops borbonicus* | This study | SRR12615523 | 17-717 | Le Pavillon, Reunion | -- | Toulouse, LBBE | GenoToul | GRY | -- | -21.18860 | 55.44940 | 400 | -- |
| *Zosterops borbonicus* | This study | SRR12615522 | JB41 | Grande Chaloupe, Reunion | 2008 | Toulouse, LBBE | GenoToul | GHB | M | -20.90052 | 55.37582 | 35 | -- |
| *Zosterops borbonicus* | This study | SRR12615521 | JB58 | Petit Bernica, Reunion | 2008 | Toulouse, LBBE | GenoToul | LBHB | -- | -21.02543 | 55.28032 | 306 | -- |
| *Zosterops borbonicus* | Leroy et al. | SRX5667783, | 15-179 | -- | -- | -- | -- | -- | -- | -- | -- | -- | -- |
| *Zosterops borbonicus* | Bourgeois et | ERR1753737 | 314 | -- | -- | -- | -- | -- | -- | -- | -- | -- | -- |
| *Zosterops borbonicus* | Bourgeois et | ERR1753738 | 317 | -- | -- | -- | -- | -- | -- | -- | -- | -- | -- |
| *Zosterops borbonicus* | Bourgeois et | ERR1753742 | 1430 | -- | -- | -- | -- | -- | -- | -- | -- | -- | -- |
| *Zosterops borbonicus* | Bourgeois et | ERR1753743 | 1434 | -- | -- | -- | -- | -- | -- | -- | -- | -- | -- |
| *Zosterops borbonicus* | Bourgeois et | ERR1753746 | 1337 | -- | -- | -- | -- | -- | -- | -- | -- | -- | -- |
| *Zosterops borbonicus* | Bourgeois et | ERR1753747 | 1588 | -- | -- | -- | -- | -- | -- | -- | -- | -- | -- |
| *Zosterops mauritianus* | This study | SRR12604653 | 461 | Bel Ombre Forest, Mauritius | 2007 | Toulouse, LBBE | GenoToul | -- | M | -20.47353 | 57.42066 | 278 | -- |
| *Zosterops mauritianus* | This study | SRR12604652 | 1281 | Le Bouchon, Mauritius | - | Toulouse, LBBE | GenoToul | -- | M | -20.47091 | 57.67862 | 15 | -- |
| *Zosterops mauritianus* | This study | SRR12604651 | 1312 | Cap Malheureux, Mauritius | 2009 | Toulouse, LBBE | GenoToul | -- | F | -19.98935 | 57.62839 | 21 | -- |
| *Zosterops mauritianus* | This study | SRR12604650 | 1310 | Roches Noires forest, | - | Toulouse, LBBE | GenoToul | -- | M | -20.11532 | 57.73568 | 15 | -- |
| *Zosterops mauritianus* | This study | SRR12604649 | 391 | Yemen, Mauritius | 2007 | Toulouse, LBBE | GenoToul | -- | F | -20.34319 | 57.41384 | 164 | -- |
| *Zosterops mauritianus* | This study | SRR12604648 | 1326 | Macchabé–Brise Fer forest, | 2009 | Toulouse, LBBE | GenoToul | -- | M | -20.37866 | 57.44069 | 600 | -- |
| *Zosterops mauritianus* | This study | SRR12604647 | 398 | Black River Gorges, Mauri- | 2007 | Toulouse, LBBE | GenoToul | -- | M | -20.38356 | 57.41987 | 117 | -- |
| *Zosterops mauritianus* | This study | SRR12604646 | 1319 | Le Pouce Mt, Mauritius | - | Toulouse, LBBE | GenoToul | -- | M | -20.19902 | 57.52384 | 605 | -- |
| *Zosterops mauritianus* | This study | SRR12604645 | 1295 | Le Morne Brabant, Mauri- | - | Toulouse, LBBE | GenoToul | -- | M | -20.45976 | 57.33227 | 9 | -- |
| *Zosterops olivaceus* | This study | SRR12717624 | 11-1034 |  | - | Toulouse, LBBE | GenoToul | -- | -- | -21.18372 | 55.83008 | 55 | -- |
| *Zosterops olivaceus* | This study | SRR12717630 | 236 | Basse Vallée , Reunion | - | Toulouse, LBBE | GenoToul | -- | -- | -21.34094 | 55.70902 | 687 | -- |
| *Zosterops olivaceus* | This study | SRR12717628 | 322 | Maïdo, Reunion | - | Toulouse, LBBE | GenoToul | -- | -- | -21.07276 | 55.37742 | 2062 | -- |
| *Zosterops olivaceus* | This study | SRR12717631 | 354 | Roche Verre Bouteille , | - | Toulouse, LBBE | GenoToul | -- | -- | -20.98591 | 55.39621 | 1134 | -- |
| *Zosterops olivaceus* | This study | SRR12717625 | 1113 | Sentier Ste-Marguerite, | - | Toulouse, LBBE | GenoToul | -- | -- | -21.10876 | 55.68827 | 550 | -- |
| *Zosterops olivaceus* | This study | SRR12717632 | 1418 | Canot, Reunion | - | Toulouse, LBBE | GenoToul | -- | -- | -21.15376 | 55.30325 | 518 | -- |
| *Zosterops olivaceus* | This study | SRR12717627 | 1550 | Moka, Reunion | 2010 | Toulouse, LBBE | GenoToul | -- | -- | -20.92789 | 55.51502 | 219 | -- |
| *Zosterops olivaceus* | This study | SRR12717623 | 1649 | Bébour, Réunion | 2010 | Toulouse, LBBE | GenoToul | -- | -- | -21.09650 | 55.54945 | 1540 | -- |
| *Zosterops olivaceus* | This study | SRR12717626 | 2217 | Nez de Bœuf, Reunion | - | Toulouse, LBBE | GenoToul | -- | -- | -21.20615 | 55.61893 | 2070 | -- |
| *Zosterops olivaceus* | This study | SRR12717633 | 15-026 | Ravine Petit St-Pierre, | 2015 | Montpellier ISEM | Novogene | -- | -- | -21.18495 | 55.67216 | 2083 | -- |
| *Zosterops olivaceus* | This study | SRR12717622 | 15-250 | Nez de Bœuf, Reunion | 2015 | Montpellier ISEM | Novogene | -- | -- | -21.20615 | 55.61893 | 2070 | -- |
| *Zosterops olivaceus* | This study | SRR12717621 | 15-258 | Trois citernes, Réunion | 2015 | Montpellier ISEM | Novogene | -- | -- | -21.15885 | 55.76181 | 846 | -- |
| *Zosterops olivaceus* | This study | SRR12717635 | 15-261 | Trois citernes, Réunion | 2015 | Montpellier ISEM | Novogene | -- | -- | -21.15885 | 55.76181 | 846 | -- |
| *Zosterops olivaceus* | This study | SRR12717634 | 15-262 | Trois citernes, Réunion | 2015 | Montpellier ISEM | Novogene | -- | -- | -21.15885 | 55.76181 | 846 | -- |
| *Zosterops olivaceus* | This study | SRR12717629 | 17-691 | Grand Matarum, Reunion | 2017 | Toulouse, LBBE | GenoToul | -- | -- | -21.12443 | 55.47868 | 1460 | -- |
| *Zosterops pallidus* | Leroy et al. | SRX5651175 | BN72 MNHN | Free state province, | 2015 | Montpellier ISEM | Novogene | -- | F | -29.75508 | 25.17733 | -- | -- |
| *Zosterops pallidus* | This study | SRX9239935 | BN63 | Free state province, | 2015 | Montpellier ISEM | Novogene | -- | M | -29.75508 | 25.17733 | -- | -- |
| *Zosterops virens* | This study | SRR12728740 | MNHN Uncat. | Doodsklip Camp, Bavi- | 2014 | Montpellier ISEM | Novogene | -- | M | -33.65875 | 24.43164 | -- | -- |

| | | | | | | | | | | | | | |
|---|---|---|---|---|---|---|---|---|---|---|---|---|---|
| *Zosterops virens* | This study | SRR12728739 | MNHN Uncat. | close to cottage, | 2014 | Montpellier ISEM | Novogene | -- | -- | -- | -- | -- | -- |
| *Zosterops virens* | This study | SRR12728736 | MNHN ZO | Upper Glass Nevin Farm, | 2014 | Montpellier ISEM | Novogene | -- | M | -- | -- | -- | -- |
| *Zosterops virens* | This study | SRR12728735 | MNHN Uncat. | close to Research Center, | 2014 | Montpellier ISEM | Novogene | -- | F | -- | -- | -- | -- |
| *Zosterops virens* | This study | SRR12728734 | MNHN Uncat. | Upper Glass Nevin Farm, | 2014 | Montpellier ISEM | Novogene | -- | M | -- | -- | -- | -- |
| *Zosterops virens* | This study | SRR12728733 | MNHN Uncat. | close to Research Center, | 20014 | Montpellier ISEM | Novogene | -- | M | -- | -- | -- | -- |
| *Zosterops virens* | This study | SRR12728732 | MNHN Uncat. | Fort Fordyce, close to | 2014 | Montpellier ISEM | Novogene | -- | F | -- | -- | -- | -- |
| *Zosterops virens* | This study | SRR12728731 | MNHN ZO | Upper Glass Nevin Farm, | 2014 | Montpellier ISEM | Novogene | -- | M | -- | -- | -- | -- |
| *Zosterops virens* | This study | SRR12728730 | MNHN Uncat. | Research Center, Great Fish | 2014 | Montpellier ISEM | Novogene | -- | M | -- | -- | -- | -- |
| *Zosterops virens* | This study | SRR12728729 | RCKB_1917_2 | Upper Glass Nevin Farm, | 2014 | Montpellier ISEM | Novogene | -- | M | -- | -- | -- | -- |
| *Zosterops virens* | This study | SRR12728738 | MNHN Uncat. | Upper Glass Nevin Farm, | 2014 | Montpellier ISEM | Novogene | -- | M | -- | -- | -- | -- |
| *Zosterops virens* | This study | SRR12728737 | MNHN ZO | Doodsklip Camp, Bavi- | 2014 | Montpellier ISEM | Novogene | -- | -- | -33.65875 | 24.43164 | -- | -- |
| *Fringilla coelebs* | This study | SRR12816841 | CHASE9 | Segovia, Iberian Peninsula | 2018 | Madrid, CSIC | Novogene | -- | F | -4.26463 | 40.81017 | 971 | 1112352 |
| *Fringilla coelebs* | This study | SRR12816842 | CHASE8 | Segovia, Iberian Peninsula | 2018 | Madrid, CSIC | Novogene | -- | M | -4.26463 | 40.81017 | 971 | 1112343 |
| *Fringilla coelebs* | This study | SRR12816843 | CHASE7 | Segovia, Iberian Peninsula | 2018 | Madrid, CSIC | Novogene | -- | M | -4.26463 | 40.81017 | 971 | 1112342 |
| *Fringilla coelebs* | This study | SRR12816844 | CHASE6 | Segovia, Iberian Peninsula | 2018 | Madrid, CSIC | Novogene | -- | M | -4.26463 | 40.81017 | 971 | 1112341 |
| *Fringilla coelebs* | This study | SRR12816846 | CHASE5 | Segovia, Iberian Peninsula | 2018 | Madrid, CSIC | Novogene | -- | M | -4.26463 | 40.81017 | 971 | 1112340 |
| *Fringilla coelebs* | This study | SRR12816847 | CHASE4 | Segovia, Iberian Peninsula | 2018 | Madrid, CSIC | Novogene | -- | M | -4.26463 | 40.81017 | 971 | 1112339 |
| *Fringilla coelebs* | This study | SRR12816848 | CHASE3 | Segovia, Iberian Peninsula | 2016 | Madrid, CSIC | Novogene | -- | M | -4.26463 | 40.81017 | 971 | 1112273 |
| *Fringilla coelebs* | This study | SRR12816849 | CHASE2 | Segovia, Iberian Peninsula | 2016 | Madrid, CSIC | Novogene | -- | M | -4.26463 | 40.81017 | 971 | 1112308 |
| *Fringilla coelebs* | This study | SRR12816850 | CHASE1 | Segovia, Iberian Peninsula | 2016 | Madrid, CSIC | Novogene | -- | M | -4.26463 | 40.81017 | 971 | 1112303 |
| *Fringilla coelebs palmae* | This study | SRR12816834 | CHAFU10 | La palma, Canary island | 2016 | Madrid, CSIC | Novogene | -- | M | -17.83315 | 28.51912 | 1110 | 1112398 |
| *Fringilla coelebs palmae* | This study | SRR12816835 | CHAFU9 | La palma, Canary island | 2016 | Madrid, CSIC | Novogene | -- | M | -17.83315 | 28.51912 | 1110 | 1112393 |
| *Fringilla coelebs palmae* | This study | SRR12816836 | CHAFU8 | La palma, Canary island | 2016 | Madrid, CSIC | Novogene | -- | M | -17.83315 | 28.51912 | 1110 | 1112408 |
| *Fringilla coelebs palmae* | This study | SRR12816837 | CHAFU7 | La palma, Canary island | 2016 | Madrid, CSIC | Novogene | -- | M | -17.83315 | 28.51912 | 1110 | 1112407 |
| *Fringilla coelebs palmae* | This study | SRR12816838 | CHAFU6 | La palma, Canary island | 2016 | Madrid, CSIC | Novogene | -- | M | -17.83315 | 28.51912 | 1110 | 1112406 |
| *Fringilla coelebs palmae* | This study | SRR12816839 | CHAFU5 | La palma, Canary island | 2016 | Madrid, CSIC | Novogene | -- | M | -17.83315 | 28.51912 | 1110 | 1112405 |
| *Fringilla coelebs palmae* | This study | SRR12816840 | CHAFU4 | La palma, Canary island | 2016 | Madrid, CSIC | Novogene | -- | M | -17.83315 | 28.51912 | 1110 | 1112404 |
| *Fringilla coelebs palmae* | This study | SRR12816845 | CHAFU3 | La palma, Canary island | 2016 | Madrid, CSIC | Novogene | -- | M | -17.83315 | 28.51912 | 1110 | 1112396 |
| *Fringilla coelebs palmae* | This study | SRR12816851 | CHASE12 | La palma, Canary island | 2018 | Madrid, CSIC | Novogene | -- | M | -17.83315 | 28.51912 | 1110 | 1112397 |
| *Fringilla coelebs palmae* | This study | SRR12816852 | CHASE11 | La palma, Canary island | 2018 | Madrid, CSIC | Novogene | -- | M | -17.83315 | 28.51912 | 1110 | 1112596 |
| *Fringilla coelebs palmae* | This study | SRR12816853 | CHASE10 | La palma, Canary island | 2018 | Madrid, CSIC | Novogene | -- | M | -17.83315 | 28.51912 | 1110 | 1112595 |
| *Fringilla coelebs palmae* | This study | SRR12816854 | CHAFU12 | La palma, Canary island | 2016 | Madrid, CSIC | Novogene | -- | M | -17.83315 | 28.51912 | 1110 | 1112411 |
| *Fringilla coelebs palmae* | This study | SRR12816855 | CHAFU11 | La palma, Canary island | 2016 | Madrid, CSIC | Novogene | -- | M | -17.83315 | 28.51912 | 1110 | 1112400 |
| *Fringilla coelebs palmae* | This study | SRR12816856 | CHAFU2 | La palma, Canary island | 2016 | Madrid, CSIC | Novogene | -- | M | -17.83315 | 28.51912 | 1110 | 1112387 |
| *Fringilla coelebs palmae* | This study | SRR12816857 | CHAFU1 | La palma, Canary island | 2016 | Madrid, CSIC | Novogene | -- | M | -17.83315 | 28.51912 | 1110 | 1112386 |
| *Fringilla teydea* | This study | SRR12728685 | B2A446882 | Mña Cedro, Tenerife | 2016 | Madrid, CSIC | Novogene | -- | F | -- | -- | -- | -- |
| *Fringilla teydea* | This study | SRR12728686 | B2A446881 | Mña Cedro, Tenerife | 2016 | Madrid, CSIC | Novogene | -- | M | -- | -- | -- | -- |
| *Fringilla teydea* | This study | SRR12728687 | B2A446880 | El Portillo, Tenerife | 2016 | Madrid, CSIC | Novogene | -- | M | -- | -- | -- | -- |
| *Fringilla teydea* | This study | SRR12728688 | B2A446879 | El Portillo, Tenerife | 2016 | Madrid, CSIC | Novogene | -- | M | -- | -- | -- | -- |
| *Fringilla teydea* | This study | SRR12728689 | B2A446875 | Mña Cedro, Tenerife | 2016 | Madrid, CSIC | Novogene | -- | M | -- | -- | -- | -- |
| *Fringilla teydea* | This study | SRR12728690 | B2A446874 | Mña Cedro, Tenerife | 2016 | Madrid, CSIC | Novogene | -- | F | -- | -- | -- | -- |
| *Fringilla teydea* | This study | SRR12728691 | B2A446873 | Mña Cedro, Tenerife | 2016 | Madrid, CSIC | Novogene | -- | F | -- | -- | -- | -- |
| *Fringilla teydea* | This study | SRR12728692 | B2A446872 | Mña Cedro, Tenerife | 2016 | Madrid, CSIC | Novogene | -- | M | -- | -- | -- | -- |
| *Fringilla teydea* | This study | SRR12728693 | B2A446871 | Mña Cedro, Tenerife | 2016 | Madrid, CSIC | Novogene | -- | F | -- | -- | -- | -- |
| *Fringilla teydea* | This study | SRR12728694 | B2A446870 | El Portillo, Tenerife | 2016 | Madrid, CSIC | Novogene | -- | F | -- | -- | -- | -- |
| *Taeniopygia guttata cas-* | Singhal et al. | ERR1013161 | 26462 | -- | -- | -- | -- | -- | -- | -- | -- | -- | -- |
| *Taeniopygia guttata cas-* | Singhal et al. | ERR1013162 | 28339 | -- | -- | -- | -- | -- | -- | -- | -- | -- | -- |
| *Taeniopygia guttata cas-* | Singhal et al. | ERR1013163 | 28353 | -- | -- | -- | -- | -- | -- | -- | -- | -- | -- |
| *Taeniopygia guttata cas-* | Singhal et al. | ERR1013164 | 26721 | -- | -- | -- | -- | -- | -- | -- | -- | -- | -- |
| *Taeniopygia guttata cas-* | Singhal et al. | ERR1013165 | 28456 | -- | -- | -- | -- | -- | -- | -- | -- | -- | -- |
| *Taeniopygia guttata cas-* | Singhal et al. | ERR1013166 | 28402 | -- | -- | -- | -- | -- | -- | -- | -- | -- | -- |
| *Taeniopygia guttata cas-* | Singhal et al. | ERR1013167 | 26516 | -- | -- | -- | -- | -- | -- | -- | -- | -- | -- |
| *Taeniopygia guttata cas-* | Singhal et al. | ERR1013168 | 28404 | -- | -- | -- | -- | -- | -- | -- | -- | -- | -- |
| *Taeniopygia guttata cas-* | Singhal et al. | ERR1013169 | 26820 | -- | -- | -- | -- | -- | -- | -- | -- | -- | -- |
| *Taeniopygia guttata cas-* | Singhal et al. | ERR1013170 | 26733 | -- | -- | -- | -- | -- | -- | -- | -- | -- | -- |
| *Taeniopygia guttata cas-* | Singhal et al. | ERR1013171 | 28481 | -- | -- | -- | -- | -- | -- | -- | -- | -- | -- |
| *Taeniopygia guttata cas-* | Singhal et al. | ERR1013172 | 26881 | -- | -- | -- | -- | -- | -- | -- | -- | -- | -- |

| | | | | | | | | | | | | | |
|---|---|---|---|---|---|---|---|---|---|---|---|---|---|
| *Taeniopygia guttata cas-* | Singhal et al. | ERR1013173 | 26781 | -- | -- | -- | -- | -- | -- | -- | -- | -- | -- |
| *Taeniopygia guttata cas-* | Singhal et al. | ERR1013174 | 26896 | -- | -- | -- | -- | -- | -- | -- | -- | -- | -- |
| *Taeniopygia guttata cas-* | Singhal et al. | ERR1013175 | 26792 | -- | -- | -- | -- | -- | -- | -- | -- | -- | -- |
| *Taeniopygia guttata cas-* | Singhal et al. | ERR1013176 | 28016 | -- | -- | -- | -- | -- | -- | -- | -- | -- | -- |
| *Taeniopygia guttata cas-* | Singhal et al. | ERR1013177 | 26795 | -- | -- | -- | -- | -- | -- | -- | -- | -- | -- |
| *Taeniopygia guttata cas-* | Singhal et al. | ERR1013178 | 28078 | -- | -- | -- | -- | -- | -- | -- | -- | -- | -- |
| *Taeniopygia guttata cas-* | Singhal et al. | ERR1013179 | 28313 | -- | -- | -- | -- | -- | -- | -- | -- | -- | -- |
| *Poephila acuticauda acuti-* | Singhal et al. | ERR1013135 | G111 | -- | -- | -- | -- | -- | -- | -- | -- | -- | -- |
| *Poephila acuticauda acuti-* | Singhal et al. | ERR1013136 | G118 | -- | -- | -- | -- | -- | -- | -- | -- | -- | -- |
| *Poephila acuticauda acuti-* | Singhal et al. | ERR1013137 | G163 | -- | -- | -- | -- | -- | -- | -- | -- | -- | -- |
| *Poephila acuticauda acuti-* | Singhal et al. | ERR1013138 | G169 | -- | -- | -- | -- | -- | -- | -- | -- | -- | -- |
| *Poephila acuticauda acuti-* | Singhal et al. | ERR1013139 | G183 | -- | -- | -- | -- | -- | -- | -- | -- | -- | -- |
| *Poephila acuticauda acuti-* | Singhal et al. | ERR1013140 | G250 | -- | -- | -- | -- | -- | -- | -- | -- | -- | -- |
| *Poephila acuticauda acuti-* | Singhal et al. | ERR1013141 | G276 | -- | -- | -- | -- | -- | -- | -- | -- | -- | -- |
| *Poephila acuticauda acuti-* | Singhal et al. | ERR1013142 | G294 | -- | -- | -- | -- | -- | -- | -- | -- | -- | -- |
| *Poephila acuticauda acuti-* | Singhal et al. | ERR1013143 | W2703 | -- | -- | -- | -- | -- | -- | -- | -- | -- | -- |
| *Poephila acuticauda acuti-* | Singhal et al. | ERR1013144 | W2994 | -- | -- | -- | -- | -- | -- | -- | -- | -- | -- |
| *Parus major* | Corcoran et | SRR5423293 | TR44666 | -- | -- | -- | -- | -- | -- | -- | -- | -- | -- |
| *Parus major* | Corcoran et | SRR5423294 | 943 | -- | -- | -- | -- | -- | -- | -- | -- | -- | -- |
| *Parus major* | Corcoran et | SRR5423295 | 917 | -- | -- | -- | -- | -- | -- | -- | -- | -- | -- |
| *Parus major* | Corcoran et | SRR5423296 | 61 | -- | -- | -- | -- | -- | -- | -- | -- | -- | -- |
| *Parus major* | Corcoran et | SRR5423297 | 318 | -- | -- | -- | -- | -- | -- | -- | -- | -- | -- |
| *Parus major* | Corcoran et | SRR5423298 | 249 | -- | -- | -- | -- | -- | -- | -- | -- | -- | -- |
| *Parus major* | Corcoran et | SRR5423299 | 167 | -- | -- | -- | -- | -- | -- | -- | -- | -- | -- |
| *Parus major* | Corcoran et | SRR5423300 | 15 | -- | -- | -- | -- | -- | -- | -- | -- | -- | -- |
| *Parus major* | Corcoran et | SRR5423301 | 1485 | -- | -- | -- | -- | -- | -- | -- | -- | -- | -- |
| *Parus major* | Corcoran et | SRR5423302 | 1280 | -- | -- | -- | -- | -- | -- | -- | -- | -- | -- |
| *Phylloscopus trochilus* | Lundberg et | SRX1764074 to | 00G03 | -- | -- | -- | -- | -- | -- | -- | -- | -- | -- |
| *Phylloscopus trochilus* | Lundberg et | SRX1764080 to | 00G04 | -- | -- | -- | -- | -- | -- | -- | -- | -- | -- |
| *Phylloscopus trochilus* | Lundberg et | SRX1764086 to | 00G10 | -- | -- | -- | -- | -- | -- | -- | -- | -- | -- |
| *Phylloscopus trochilus* | Lundberg et | SRX1764092 to | 00J01 | -- | -- | -- | -- | -- | -- | -- | -- | -- | -- |
| *Phylloscopus trochilus* | Lundberg et | SRX1764146 to | 01P02 | -- | -- | -- | -- | -- | -- | -- | -- | -- | -- |
| *Phylloscopus trochilus* | Lundberg et | SRX1764152 to | 03K06 | -- | -- | -- | -- | -- | -- | -- | -- | -- | -- |
| *Phylloscopus trochilus* | Lundberg et | SRX1764158 to | 96A01 | -- | -- | -- | -- | -- | -- | -- | -- | -- | -- |
| *Phylloscopus trochilus* | Lundberg et | SRX1764164 to | 96B07 | -- | -- | -- | -- | -- | -- | -- | -- | -- | -- |
| *Phylloscopus trochilus* | Lundberg et | SRX1764170 to | 97A12 | -- | -- | -- | -- | -- | -- | -- | -- | -- | -- |
| *Certhidea olivacea (S)* | Lamichhaney | SRX728428 to SRX728430 | Co13 | -- | -- | -- | -- | -- | -- | -- | -- | -- | -- |
| *Certhidea olivacea (S)* | Lamichhaney | SRX728431 to SRX728435 | Co14 | -- | -- | -- | -- | -- | -- | -- | -- | -- | -- |
| *Certhidea olivacea (S)* | Lamichhaney | SRX728436; SRX728437 | Co15 | -- | -- | -- | -- | -- | -- | -- | -- | -- | -- |
| *Certhidea olivacea (S)* | Lamichhaney | SRX728438; SRX728439 | Co16 | -- | -- | -- | -- | -- | -- | -- | -- | -- | -- |
| *Certhidea olivacea (S)* | Lamichhaney | SRX728440; SRX728441 | Co20 | -- | -- | -- | -- | -- | -- | -- | -- | -- | -- |
| *Certhidea fusca (E)* | Lamichhaney | SRX728406; SRX728407 | CfE9 | -- | -- | -- | -- | -- | -- | -- | -- | -- | -- |
| *Certhidea fusca (E)* | Lamichhaney | SRX728404; SRX728405 | CfE8 | -- | -- | -- | -- | -- | -- | -- | -- | -- | -- |
| *Certhidea fusca (E)* | Lamichhaney | SRX728402; SRX728403 | CfE7 | -- | -- | -- | -- | -- | -- | -- | -- | -- | -- |
| *Certhidea fusca (E)* | Lamichhaney | SRX728400; SRX728401 | CfE6 | -- | -- | -- | -- | -- | -- | -- | -- | -- | -- |
| *Certhidea fusca (E)* | Lamichhaney | SRX728398; SRX728399 | CfE5 | -- | -- | -- | -- | -- | -- | -- | -- | -- | -- |
| *Certhidea fusca (E)* | Lamichhaney | SRX728396; SRX728397 | CfE4 | -- | -- | -- | -- | -- | -- | -- | -- | -- | -- |
| *Certhidea fusca (E)* | Lamichhaney | SRX728394; SRX728395 | CfE3 | -- | -- | -- | -- | -- | -- | -- | -- | -- | -- |
| *Certhidea fusca (E)* | Lamichhaney | SRX728392; SRX728393 | CfE2 | -- | -- | -- | -- | -- | -- | -- | -- | -- | -- |
| *Certhidea fusca (E)* | Lamichhaney | SRX728390; SRX728391 | CfE10 | -- | -- | -- | -- | -- | -- | -- | -- | -- | -- |
| *Certhidea fusca (E)* | Lamichhaney | SRX728388; SRX728389 | CfE1 | -- | -- | -- | -- | -- | -- | -- | -- | -- | -- |
| *Certhidea fusca (L)* | Lamichhaney | SRX728386; SRX728387 | Cfc9 | -- | -- | -- | -- | -- | -- | -- | -- | -- | -- |
| *Certhidea fusca (L)* | Lamichhaney | SRX728384; SRX728385 | Cfc8 | -- | -- | -- | -- | -- | -- | -- | -- | -- | -- |
| *Certhidea fusca (L)* | Lamichhaney | SRX728382; SRX728383 | Cfc7 | -- | -- | -- | -- | -- | -- | -- | -- | -- | -- |
| *Certhidea fusca (L)* | Lamichhaney | SRX728380; SRX728381 | Cfc6 | -- | -- | -- | -- | -- | -- | -- | -- | -- | -- |
| *Certhidea fusca (L)* | Lamichhaney | SRX728378; SRX728379 | Cfc5 | -- | -- | -- | -- | -- | -- | -- | -- | -- | -- |
| *Certhidea fusca (L)* | Lamichhaney | SRX728376; SRX728377 | Cfc4 | -- | -- | -- | -- | -- | -- | -- | -- | -- | -- |

| | | | | | | | | | | | | | |
|---|---|---|---|---|---|---|---|---|---|---|---|---|---|
| *Certhidea fusca (L)* | Lamichhaney | SRX728374; SRX728375 | Cfc3 | -- | -- | -- | -- | -- | -- | -- | -- | -- | -- |
| *Certhidea fusca (L)* | Lamichhaney | SRX728372; SRX728373 | Cfc2 | -- | -- | -- | -- | -- | -- | -- | -- | -- | -- |
| *Certhidea fusca (L)* | Lamichhaney | SRX728370; SRX728371 | Cfc10 | -- | -- | -- | -- | -- | -- | -- | -- | -- | -- |
| *Certhidea fusca (L)* | Lamichhaney | SRX728368; SRX728369 | Cfc1 | -- | -- | -- | -- | -- | -- | -- | -- | -- | -- |
| *Platyspiza crassirostris (Z)* | Lamichhaney | SRX728584; SRX728585 | PL9 | -- | -- | -- | -- | -- | -- | -- | -- | -- | -- |
| *Platyspiza crassirostris (Z)* | Lamichhaney | SRX728582; SRX728583 | PL7 | -- | -- | -- | -- | -- | -- | -- | -- | -- | -- |
| *Platyspiza crassirostris (Z)* | Lamichhaney | SRX728580; SRX728581 | PL4 | -- | -- | -- | -- | -- | -- | -- | -- | -- | -- |
| *Platyspiza crassirostris (Z)* | Lamichhaney | SRX728577; SRX728578; | PL16 | -- | -- | -- | -- | -- | -- | -- | -- | -- | -- |
| *Platyspiza crassirostris (Z)* | Lamichhaney | SRX728575; SRX728576 | PL15 | -- | -- | -- | -- | -- | -- | -- | -- | -- | -- |
| *Camarhynchus pallidus (Z)* | Lamichhaney | SRX728545; SRX728546 | PAL5 | -- | -- | -- | -- | -- | -- | -- | -- | -- | -- |
| *Camarhynchus pallidus (Z)* | Lamichhaney | SRX728543; SRX728544 | PAL4 | -- | -- | -- | -- | -- | -- | -- | -- | -- | -- |
| *Camarhynchus pallidus (Z)* | Lamichhaney | SRX728541; SRX728542 | PAL3 | -- | -- | -- | -- | -- | -- | -- | -- | -- | -- |
| *Camarhynchus pallidus (Z)* | Lamichhaney | SRX728539; SRX728540 | PAL2 | -- | -- | -- | -- | -- | -- | -- | -- | -- | -- |
| *Camarhynchus pallidus (Z)* | Lamichhaney | SRX728537; SRX728538 | PAL1 | -- | -- | -- | -- | -- | -- | -- | -- | -- | -- |
| *Pinaroloxias inornata (C)* | Lamichhaney | SRX728572 to SRX728574 | PIN8 | -- | -- | -- | -- | -- | -- | -- | -- | -- | -- |
| *Pinaroloxias inornata (C)* | Lamichhaney | SRX728567 to SRX728571 | PIN7 | -- | -- | -- | -- | -- | -- | -- | -- | -- | -- |
| *Pinaroloxias inornata (C)* | Lamichhaney | SRX728565; SRX728566 | PIN6 | -- | -- | -- | -- | -- | -- | -- | -- | -- | -- |
| *Pinaroloxias inornata (C)* | Lamichhaney | SRX728563; SRX728564 | PIN5 | -- | -- | -- | -- | -- | -- | -- | -- | -- | -- |
| *Pinaroloxias inornata (C)* | Lamichhaney | SRX728561; SRX728562 | PIN4 | -- | -- | -- | -- | -- | -- | -- | -- | -- | -- |
| *Pinaroloxias inornata (C)* | Lamichhaney | SRX728559; SRX728560 | PIN3 | -- | -- | -- | -- | -- | -- | -- | -- | -- | -- |
| *Pinaroloxias inornata (C)* | Lamichhaney | SRX728557; SRX728558 | PIN2 | -- | -- | -- | -- | -- | -- | -- | -- | -- | -- |
| *Pinaroloxias inornata (C)* | Lamichhaney | SRX728555; SRX728556 | PIN1 | -- | -- | -- | -- | -- | -- | -- | -- | -- | -- |
| *Geospiza difficilis (P)* | Lamichhaney | SRX728478; SRX728479 | DP9 | -- | -- | -- | -- | -- | -- | -- | -- | -- | -- |
| *Geospiza difficilis (P)* | Lamichhaney | SRX728473 to SRX728477 | DP7 | -- | -- | -- | -- | -- | -- | -- | -- | -- | -- |
| *Geospiza difficilis (P)* | Lamichhaney | SRX728471; SRX728472 | DP6 | -- | -- | -- | -- | -- | -- | -- | -- | -- | -- |
| *Geospiza difficilis (P)* | Lamichhaney | SRX728469; SRX728470 | DP5 | -- | -- | -- | -- | -- | -- | -- | -- | -- | -- |
| *Geospiza difficilis (P)* | Lamichhaney | SRX728467; SRX728468 | DP4 | -- | -- | -- | -- | -- | -- | -- | -- | -- | -- |
| *Geospiza difficilis (P)* | Lamichhaney | SRX728465; SRX728466 | DP3 | -- | -- | -- | -- | -- | -- | -- | -- | -- | -- |
| *Geospiza difficilis (P)* | Lamichhaney | SRX728463; SRX728464 | DP2 | -- | -- | -- | -- | -- | -- | -- | -- | -- | -- |
| *Geospiza difficilis (P)* | Lamichhaney | SRX728461; SRX728462 | DP13 | -- | -- | -- | -- | -- | -- | -- | -- | -- | -- |
| *Geospiza difficilis (P)* | Lamichhaney | SRX728459; SRX728460 | DP11 | -- | -- | -- | -- | -- | -- | -- | -- | -- | -- |
| *Geospiza difficilis (P)* | Lamichhaney | SRX728457; SRX728458 | DP10 | -- | -- | -- | -- | -- | -- | -- | -- | -- | -- |
| *Geospiza septentrionalis (W)* | Lamichhaney | SRX728499; SRX728500 | DW9 | -- | -- | -- | -- | -- | -- | -- | -- | -- | -- |
| *Geospiza septentrionalis (W)* | Lamichhaney | SRX728496 to SRX728498 | DW8 | -- | -- | -- | -- | -- | -- | -- | -- | -- | -- |
| *Geospiza septentrionalis (W)* | Lamichhaney | SRX728494; SRX728495 | DW7 | -- | -- | -- | -- | -- | -- | -- | -- | -- | -- |
| *Geospiza septentrionalis (W)* | Lamichhaney | SRX728492; SRX728493 | DW6 | -- | -- | -- | -- | -- | -- | -- | -- | -- | -- |
| *Geospiza septentrionalis (W)* | Lamichhaney | SRX728490; SRX728491 | DW5 | -- | -- | -- | -- | -- | -- | -- | -- | -- | -- |
| *Geospiza septentrionalis (W)* | Lamichhaney | SRX728488; SRX728489 | DW3 | -- | -- | -- | -- | -- | -- | -- | -- | -- | -- |
| *Geospiza septentrionalis (W)* | Lamichhaney | SRX728485 to SRX728487 | DW2 | -- | -- | -- | -- | -- | -- | -- | -- | -- | -- |
| *Geospiza septentrionalis (W)* | Lamichhaney | SRX728483;SRX728484 | DW1 | -- | -- | -- | -- | -- | -- | -- | -- | -- | -- |
| *Geospiza conirostris (E)* | Lamichhaney | SRX728365; SRX728367 | CE9 | -- | -- | -- | -- | -- | -- | -- | -- | -- | -- |
| *Geospiza conirostris (E)* | Lamichhaney | SRX728363; SRX728364 | CE8 | -- | -- | -- | -- | -- | -- | -- | -- | -- | -- |
| *Geospiza conirostris (E)* | Lamichhaney | SRX728361; SRX728362 | CE7 | -- | -- | -- | -- | -- | -- | -- | -- | -- | -- |
| *Geospiza conirostris (E)* | Lamichhaney | SRX728358 to SRX728360 | CE6 | -- | -- | -- | -- | -- | -- | -- | -- | -- | -- |
| *Geospiza conirostris (E)* | Lamichhaney | SRX728355 to SRX728357 | CE5 | -- | -- | -- | -- | -- | -- | -- | -- | -- | -- |
| *Geospiza conirostris (E)* | Lamichhaney | SRX728352 to SRX728354 | CE4 | -- | -- | -- | -- | -- | -- | -- | -- | -- | -- |
| *Geospiza conirostris (E)* | Lamichhaney | SRX728347 to SRX728351 | CE3 | -- | -- | -- | -- | -- | -- | -- | -- | -- | -- |
| *Geospiza conirostris (E)* | Lamichhaney | SRX728344 to SRX728346 | CE2 | -- | -- | -- | -- | -- | -- | -- | -- | -- | -- |
| *Geospiza conirostris (E)* | Lamichhaney | SRX728342; SRX728343 | CE10 | -- | -- | -- | -- | -- | -- | -- | -- | -- | -- |
| *Geospiza conirostris (E)* | Lamichhaney | SRX728339 to SRX728341 | CE1 | -- | -- | -- | -- | -- | -- | -- | -- | -- | -- |
| *Ficedula albicollis* | Burri et al. | ERR637361 | OC_2 | -- | -- | -- | -- | -- | -- | -- | -- | -- | -- |
| *Ficedula albicollis* | Burri et al. | ERR637363 | OC_4 | -- | -- | -- | -- | -- | -- | -- | -- | -- | -- |
| *Ficedula albicollis* | Burri et al. | ERR637368 | OC_10 | -- | -- | -- | -- | -- | -- | -- | -- | -- | -- |
| *Ficedula albicollis* | Burri et al. | ERR637372 | OC_HB4 | -- | -- | -- | -- | -- | -- | -- | -- | -- | -- |
| *Ficedula albicollis* | Burri et al. | ERR637373 | OC_HB5 | -- | -- | -- | -- | -- | -- | -- | -- | -- | -- |
| *Ficedula albicollis* | Burri et al. | ERR637509 | I_E30-3 | -- | -- | -- | -- | -- | -- | -- | -- | -- | -- |
| *Ficedula albicollis* | Burri et al. | ERR637511 | I_F16-2 | -- | -- | -- | -- | -- | -- | -- | -- | -- | -- |

| | | | | | | | | | | | | | |
|---|---|---|---|---|---|---|---|---|---|---|---|---|---|
| *Ficedula albicollis* | Burri et al. | ERR637512 | I_F9-1 | -- | -- | -- | -- | -- | -- | -- | -- | -- | -- |
| *Ficedula albicollis* | Burri et al. | ERR637517 | I_LC-2 | -- | -- | -- | -- | -- | -- | -- | -- | -- | -- |
| *Ficedula albicollis* | Burri et al. | ERR637518 | I_LX-2 | -- | -- | -- | -- | -- | -- | -- | -- | -- | -- |
| *Ficedula albicollis* | Burri et al. | ERR692007 | H_117 | -- | -- | -- | -- | -- | -- | -- | -- | -- | -- |
| *Ficedula albicollis* | Burri et al. | ERR692008 | H_118 | -- | -- | -- | -- | -- | -- | -- | -- | -- | -- |
| *Ficedula albicollis* | Burri et al. | ERR692013 | H_367 | -- | -- | -- | -- | -- | -- | -- | -- | -- | -- |
| *Ficedula albicollis* | Burri et al. | ERR692014 | H_377 | -- | -- | -- | -- | -- | -- | -- | -- | -- | -- |
| *Ficedula albicollis* | Burri et al. | ERR692019 | H_454 | -- | -- | -- | -- | -- | -- | -- | -- | -- | -- |
| *Ficedula albicollis* | Burri et al. | ERR692022 | H_53 | -- | -- | -- | -- | -- | -- | -- | -- | -- | -- |
| *Ficedula albicollis* | Burri et al. | ERR692051 | CZC_419135 | -- | -- | -- | -- | -- | -- | -- | -- | -- | -- |
| *Ficedula albicollis* | Burri et al. | ERR692052 | CZC_419251 | -- | -- | -- | -- | -- | -- | -- | -- | -- | -- |
| *Ficedula albicollis* | Burri et al. | ERR692055 | CZC_48837 | -- | -- | -- | -- | -- | -- | -- | -- | -- | -- |
| *Ficedula albicollis* | Burri et al. | ERR692056 | CZC_73077 | -- | -- | -- | -- | -- | -- | -- | -- | -- | -- |
| *Ficedula hypoleuca* | Burri et al. | ERR637485 | SP_11 | -- | -- | -- | -- | -- | -- | -- | -- | -- | -- |
| *Ficedula hypoleuca* | Burri et al. | ERR637488 | SP_14 | -- | -- | -- | -- | -- | -- | -- | -- | -- | -- |
| *Ficedula hypoleuca* | Burri et al. | ERR637495 | SP_SV10 | -- | -- | -- | -- | -- | -- | -- | -- | -- | -- |
| *Ficedula hypoleuca* | Burri et al. | ERR637499 | SP_SV4 | -- | -- | -- | -- | -- | -- | -- | -- | -- | -- |
| *Ficedula hypoleuca* | Burri et al. | ERR637502 | SP_SV7 | -- | -- | -- | -- | -- | -- | -- | -- | -- | -- |
| *Ficedula hypoleuca* | Burri et al. | ERR693842 | CZP_312467 | -- | -- | -- | -- | -- | -- | -- | -- | -- | -- |
| *Ficedula hypoleuca* | Burri et al. | ERR693844 | CZP_312472 | -- | -- | -- | -- | -- | -- | -- | -- | -- | -- |
| *Ficedula hypoleuca* | Burri et al. | ERR693848 | CZP_522953 | -- | -- | -- | -- | -- | -- | -- | -- | -- | -- |
| *Ficedula hypoleuca* | Burri et al. | ERR693851 | CZP_522956 | -- | -- | -- | -- | -- | -- | -- | -- | -- | -- |
| *Ficedula hypoleuca* | Burri et al. | ERR693856 | CZP_522963 | -- | -- | -- | -- | -- | -- | -- | -- | -- | -- |
| *Ficedula hypoleuca* | Burri et al. | ERR699270 | OP_CA91255 | -- | -- | -- | -- | -- | -- | -- | -- | -- | -- |
| *Ficedula hypoleuca* | Burri et al. | ERR699277 | OP_CK53268 | -- | -- | -- | -- | -- | -- | -- | -- | -- | -- |
| *Ficedula hypoleuca* | Burri et al. | ERR699283 | OP_CK53891 | -- | -- | -- | -- | -- | -- | -- | -- | -- | -- |
| *Ficedula hypoleuca* | Burri et al. | ERR699284 | OP_CK53898 | -- | -- | -- | -- | -- | -- | -- | -- | -- | -- |
| *Ficedula hypoleuca* | Burri et al. | ERR699286 | OP_CL76147 | -- | -- | -- | -- | -- | -- | -- | -- | -- | -- |
| *Ficedula hypoleuca* | Burri et al. | ERR699406 | E_F_1 | -- | -- | -- | -- | -- | -- | -- | -- | -- | -- |
| *Ficedula hypoleuca* | Burri et al. | ERR699407 | E_F_1 | -- | -- | -- | -- | -- | -- | -- | -- | -- | -- |
| *Ficedula hypoleuca* | Burri et al. | ERR699418 | E_M_40 | -- | -- | -- | -- | -- | -- | -- | -- | -- | -- |
| *Ficedula hypoleuca* | Burri et al. | ERR699420 | E_M_57 | -- | -- | -- | -- | -- | -- | -- | -- | -- | -- |
| *Ficedula hypoleuca* | Burri et al. | ERR699557 | E_M_97 | -- | -- | -- | -- | -- | -- | -- | -- | -- | -- |
| *Ficedula speculigera* | Burri et al. | ERR700462 | FSP08-001 | -- | -- | -- | -- | -- | -- | -- | -- | -- | -- |
| *Ficedula speculigera* | Burri et al. | ERR700463 | FSP08-002 | -- | -- | -- | -- | -- | -- | -- | -- | -- | -- |
| *Ficedula speculigera* | Burri et al. | ERR700464 | FSP08-006 | -- | -- | -- | -- | -- | -- | -- | -- | -- | -- |
| *Ficedula speculigera* | Burri et al. | ERR700465 | FSP08-007 | -- | -- | -- | -- | -- | -- | -- | -- | -- | -- |
| *Ficedula speculigera* | Burri et al. | ERR700466 | FSP08-009 | -- | -- | -- | -- | -- | -- | -- | -- | -- | -- |
| *Ficedula speculigera* | Burri et al. | ERR700467 | FSP08-010 | -- | -- | -- | -- | -- | -- | -- | -- | -- | -- |
| *Ficedula speculigera* | Burri et al. | ERR700468 | FSP08-011 | -- | -- | -- | -- | -- | -- | -- | -- | -- | -- |
| *Ficedula speculigera* | Burri et al. | ERR700469 | FSP08-012 | -- | -- | -- | -- | -- | -- | -- | -- | -- | -- |
| *Ficedula speculigera* | Burri et al. | ERR700470 | FSP08-014 | -- | -- | -- | -- | -- | -- | -- | -- | -- | -- |
| *Ficedula speculigera* | Burri et al. | ERR700471 | FSP08-015 | -- | -- | -- | -- | -- | -- | -- | -- | -- | -- |
| *Ficedula speculigera* | Burri et al. | ERR700472 | FSP08-017 | -- | -- | -- | -- | -- | -- | -- | -- | -- | -- |
| *Ficedula speculigera* | Burri et al. | ERR700473 | FSP08-018 | -- | -- | -- | -- | -- | -- | -- | -- | -- | -- |
| *Ficedula speculigera* | Burri et al. | ERR700474 | FSP08-019 | -- | -- | -- | -- | -- | -- | -- | -- | -- | -- |
| *Ficedula speculigera* | Burri et al. | ERR700475 | FSP08-020 | -- | -- | -- | -- | -- | -- | -- | -- | -- | -- |
| *Ficedula speculigera* | Burri et al. | ERR700476 | FSP08-021 | -- | -- | -- | -- | -- | -- | -- | -- | -- | -- |
| *Ficedula speculigera* | Burri et al. | ERR700477 | FSP08-022 | -- | -- | -- | -- | -- | -- | -- | -- | -- | -- |
| *Ficedula speculigera* | Burri et al. | ERR700478 | FSP08-023 | -- | -- | -- | -- | -- | -- | -- | -- | -- | -- |
| *Ficedula speculigera* | Burri et al. | ERR700479 | FSP08-024 | -- | -- | -- | -- | -- | -- | -- | -- | -- | -- |
| *Ficedula speculigera* | Burri et al. | ERR700480 | FSP08-025 | -- | -- | -- | -- | -- | -- | -- | -- | -- | -- |
| *Ficedula speculigera* | Burri et al. | ERR700481 | FSP08-026 | -- | -- | -- | -- | -- | -- | -- | -- | -- | -- |
| *Ficedula semitorquata* | Burri et al. | ERR700020 | FST_11 | -- | -- | -- | -- | -- | -- | -- | -- | -- | -- |
| *Ficedula semitorquata* | Burri et al. | ERR700021 | FST_16 | -- | -- | -- | -- | -- | -- | -- | -- | -- | -- |
| *Ficedula semitorquata* | Burri et al. | ERR700022 | FST_19 | -- | -- | -- | -- | -- | -- | -- | -- | -- | -- |
| *Ficedula semitorquata* | Burri et al. | ERR700023 | FST_1 | -- | -- | -- | -- | -- | -- | -- | -- | -- | -- |

| | | | | | | | | | | | | | |
|---|---|---|---|---|---|---|---|---|---|---|---|---|---|
| *Ficedula semitorquata* | Burri et al. | ERR700024 | FST_20 | -- | -- | -- | -- | -- | -- | -- | -- | -- | -- |
| *Ficedula semitorquata* | Burri et al. | ERR700025 | FST_21 | -- | -- | -- | -- | -- | -- | -- | -- | -- | -- |
| *Ficedula semitorquata* | Burri et al. | ERR700026 | FST_22 | -- | -- | -- | -- | -- | -- | -- | -- | -- | -- |
| *Ficedula semitorquata* | Burri et al. | ERR700027 | FST_24 | -- | -- | -- | -- | -- | -- | -- | -- | -- | -- |
| *Ficedula semitorquata* | Burri et al. | ERR700028 | FST_27 | -- | -- | -- | -- | -- | -- | -- | -- | -- | -- |
| *Ficedula semitorquata* | Burri et al. | ERR700029 | FST_2 | -- | -- | -- | -- | -- | -- | -- | -- | -- | -- |
| *Ficedula semitorquata* | Burri et al. | ERR700030 | FST_32 | -- | -- | -- | -- | -- | -- | -- | -- | -- | -- |
| *Ficedula semitorquata* | Burri et al. | ERR700031 | FST_34 | -- | -- | -- | -- | -- | -- | -- | -- | -- | -- |
| *Ficedula semitorquata* | Burri et al. | ERR700032 | FST_35 | -- | -- | -- | -- | -- | -- | -- | -- | -- | -- |
| *Ficedula semitorquata* | Burri et al. | ERR700033 | FST_36 | -- | -- | -- | -- | -- | -- | -- | -- | -- | -- |
| *Ficedula semitorquata* | Burri et al. | ERR700034 | FST_37 | -- | -- | -- | -- | -- | -- | -- | -- | -- | -- |
| *Ficedula semitorquata* | Burri et al. | ERR700035 | FST_39 | -- | -- | -- | -- | -- | -- | -- | -- | -- | -- |
| *Ficedula semitorquata* | Burri et al. | ERR700036 | FST_41 | -- | -- | -- | -- | -- | -- | -- | -- | -- | -- |
| *Ficedula semitorquata* | Burri et al. | ERR700037 | FST_4 | -- | -- | -- | -- | -- | -- | -- | -- | -- | -- |
| *Ficedula semitorquata* | Burri et al. | ERR700038 | FST_5 | -- | -- | -- | -- | -- | -- | -- | -- | -- | -- |
| *Ficedula semitorquata* | Burri et al. | ERR700039 | FST_7 | -- | -- | -- | -- | -- | -- | -- | -- | -- | -- |

#### Note S1: Extended introduction

Effective population size is a key parameter quantifying the magnitude of genetic drift. Endemic island species are expected to have smaller population sizes and therefore expected to be more prone to increased inbreeding and enhanced effects of genetic drift (Frankham, 2002). Indeed, islands represent limited spaces and consequently, species inhabiting islands likely represent taxa with lower census and, supposedly, lower *Ne* than continental species. If this hypothesis is true and assuming the nearly neutral theory of molecular evolution (Ohta 1973; 1992), such a lower *Ne* is expected to limit the adaptive potential of the endemic island species, because smaller populations produce lower numbers of mutations per generation at the species level and therefore carry fewer alleles that may become beneficial after an environmental change (Lanfear et al. 2014, Nam et al 2017, Gossman et al 2012, Rousselle et al. 2020). This is particularly important, because island species are expected to face higher demographic and environmental stochasticity, in such a way that a low proportion of adaptive mutations may limit the capacity to buffer or trade off these variations. In addition, *Ne* is also expected to influence the efficacy of natural selection to remove deleterious alleles, with higher effectiveness in large populations assuming the nearly neutral models of genome evolution (Ohta 1973, Charlesworth 2009; Eyre-Walker et al. 2002; Lanfear et al. 2014). Following the nearly neutral theory, slightly deleterious alleles are supposed to be accumulated at a higher rate in species with small *Ne* (as expected on islands) than in species with large *Ne* (as typically expected for species with large continental distributions), leading to an increasing deleterious mutational burden over time. Some species-specific evidence has accumulated supporting this theory for island species (Loire et al. 2013 for Giant Galápagos tortoises; Rogers and Slatkin 2017 for woolly mammoths; Robinson et al. 2016; 2018 for island foxes and Kutschera et al. 2019 for crows), but no effect of island colonization on mitochondrial molecular evolution was reported in mammals. This absence of evidence could be due to a lack of power to detect variation of *Ne* between continental and island mammals with a single locus (James et al. 2016). More generally, the strength of the relation between census size and *Ne* remains debated. Weak positive correlations between census size and nuclear genetic diversity were observed in birds and in mammals (Díez-del-Molino et al. 2018). In pinnipeds, in particular, strong correlation was recovered (Peart et al. 2020). In contrast, no relationship between range size and the level of nuclear genetic diversity were observed based on the transciptome data of 76 animal species

(Romiguier et al. 2014), but such a positive correlation was recovered using mitochondrial DNA markers (James & Eyre-Walker 2020). The absence of a general conclusion about the correlation between the effective and census population sizes was one of our main motivations for this study.

Despite its central importance, estimating *Ne* is challenging in practice. *Ne* is indeed well-known to fluctuate rapidly over short periods of time, in such a way that the current size may not reflect past population size that has shaped present-day genetic diversity (e.g. Atkinson et al. 2008; Schiffels & Durbin, 2014 for humans). For example, effective population sizes that have determined the dynamics of ancestral alleles detected using between-species (divergence) data might be several orders of magnitude different from recent *Ne* estimates using present-day within-species (polymorphism) data (Eyre-Walker 2002, Rousselle et al. 2018). In animals or plants, life-history traits such as body mass or longevity are known to provide useful proxies for long-term *Ne* (Popadin et al. 2007, Romiguier et al. 2014, Figuet et al. 2016, Chen et al. 2017). However, there is a fundamental limitation to the use of life-history traits as a proxy of *Ne* since it only holds as a result of the correlations between body size and population density (White et al. 2007). Long-term *Ne* can be more directly estimated using population genomic variables. The level of genetic drift is inversely proportional to *Ne* (Kimura & Crow, 1964; Kimura et al. 1983) and consequently, the observed nucleotide diversity levels at synonymous sites ($\pi_S$) can be used as a predictor of the long-term *Ne* because it provides an estimate of the population mutation rate ($\theta$) = $4Ne\mu$ (e.g. Romiguier et al. 2014). During last decade, methods based on the rate of coalescence such as PSMC, MSMC or SMC++ have rapidly become popular to infer historical changes in *Ne* and provide information about the demographic trajectories of a given species, including *Ne* estimates in the more recent past, which is especially important for conservation-related issues.

The impact of *Ne* on the efficacy of positive selection still remains highly debated among evolutionary biologists (e.g. Kern & Hahn, 2018 and Jensen et al., 2019 for recent publications), despite the significant progress in both the methods and the fundamental knowledge gained in this field over the last decade (e.g. Keightley and Eyre-Walker 2010; Gossmann et al. 2010; 2012; Galtier 2016; Chen et al. 2017; Rousselle et al. 2020). By combining information at both the within-species (polymorphism) and between-species (divergence) levels, the proportion of adaptive substitutions can be estimated based on the comparisons of the ratios of non-synonymous and synonymous mutations in the polymorphism and substitution data ($\pi_N/\pi_S$ & $d_N/d_S$, McDonald & Kreitman, 1991). Over the last decade, several methods inspired by the seminal work of McDonald and Kreitman (1991) have been developed to take into account short-term demographic variation and the presence of slightly deleterious mutations. These methods used the Site Frequency Spectra (SFS) at both synonymous and non-synonymous sites to estimate the Distribution of Fitness Effects (DFE) of non-synonymous mutations (Keightley & Eyre-Walker 2007; Eyre-Walker & Keightley 2009; Galtier 2016; Tataru et al. 2017; see also Moutinho et al. 2019 for a review). Using these methods, can we expect different proportions of adaptive mutations between island and continental species? On the one hand, we can hypothesize that the populations with larger effective population sizes will exhibit the higher rates of adaptive evolution, because of a greater number of *de novo* mutations produced per generation and because of the greater standing genetic variability available. On the other hand, assuming that the species with small *Ne* exhibits a higher proportion of deleterious mutations, therefore generating proteins that contribute to pulling away from their fitness optima, we can hypothesize the production of more frequent compensatory adaptive mutations in these species.

Passerine birds represent an excellent group of species to investigate the effect of insularity on evolutionary dynamic for a series of reasons. First, a lot of genomic resources are available for passerine birds. Since the first genome in 2010 (Warren et al. 2010), a lot of passerine bird species have recently joined the list (e.g. Ellegren et al. 2012; Zhang et al. 2014; Cornetti et al. 2015; Laine et al. 2016; Lundberg et al. 2017; Leroy et al. 2019). Beyond genome assemblies, several excellent population genomics studies have focused on passerine birds and the data are now available in the public domain (e.g. Burri et al. 2015; Lamichhaney et al. 2015, Lundberg et al. 2017 and Delmore *et al.* 2020 among others). Second, the large community of bird-enthusiasts, including ornithologists but also numerous voluntary birdwatchers, provides excellent monitoring of the species presence, distribution and abundance. Third, probably thanks to their good dispersal capability, a quite large diversity of birds was able to colonize islands and became endemic species (17% of the world's bird species, Johnson & Stattersfield 1990). It is important to mention that endemic islands birds have however experienced a dramatic species loss over the last five centuries (Ricketts et al. 2005). Island ecosystems are indeed more threatened than continents, especially given the deleterious consequences of human activities and the introduction of non-native species (Steadman 1995). Fourth, both the chromosome architectures and the recombination landscapes are stable in birds (Ellegren 2010; Singhal et al. 2015). Recombination landscape variation is indeed an important factor to consider because recombination rate is well known to contribute strongly to the local levels of nucleotide diversity through the effects of linked selection. Unlike mammals, bird genomes lack the *PRDM9* gene (Baker et al. 2017), a gene which explains the rapid turnover in recombination hotspots in mammals (Baudat et al. 2010; Parvanov et al. 2010). As a consequence, the correlation between the recombination rates and the G+C content is stronger in passerine birds than in mammal species for instance. This relationship is due to the GC-biased gene conversion (gBGC), a recombination-associated segregation bias that favors G and C over A and T alleles (for details, see Duret & Galtier, 2009). In birds, the genomic GC content, or more specifically GC-content at third codon position (hereafter GC3) for coding regions, are therefore a good predictor of the local recombination rate. Passerida is a clade of songbirds that originated 20 to 30 Million years ago (Barker et al. 2004; Prum et al. 2015, Oliveros et al. 2019) and represents the most species-rich avian clade. In this study, we considered songbird species with relatively similar body-mass, longevity and clutch-size to reduce the risk of some confounding factors that could correlate with *Ne*.

#### <u>Note S2:</u> Within-genome variation in the efficacy of purifying selection

In birds, the GC-content at third codon position (GC3) is known to be highly correlated with recombination rates (Hillier et al. 2004; Backström et al. 2010; Singhal et al. 2015). After ordering genes according to GC3 (see Materials and Methods), we observed negative slopes between ($\pi_N/\pi_S$) and GC3 (Fig. 1.A, B of this Note). These results were equally supported when using the estimates of $\pi_N/\pi_S$ based on the method described in Rousselle and collaborators (2019) which considers the site frequency spectra at both non-synonymous and synonymous (“method 2” in Fig. 2 & 3 of this Note). The relationship between $\pi_N/\pi_S$ and GC3 is therefore supported, and consistent with an increased efficiency of natural selection with higher local recombination rate.

By comparing sets of genes exhibiting the lowest and highest GC3 (2Mb of total concatenated sequences used for the computations), we reported significant differences in mean $\pi_N/\pi_S$ between island and mainland species in both GC-rich (t=3.316 ,p=0.003) and GC-poor (t=7.300, p=2.1x10$^{-07}$) regions, but with more marked $\pi_N/\pi_S$ differences in genes exhibiting low GC3 ($\Delta mean_{insular\ vs.\ mainland}$= 0.107, 95%CI: 0.077-0.194) than in those exhibiting high GC3 (Δmean= 0.029, 95%CI: 0.011-0.048) (Fig. 1. C, D of this Note). The genetic advantage of recombination, which is captured by the slope between the $\pi_N/\pi_S$ and the GC3 is itself highly correlated with the levels of nucleotide diversity (Fig. 1E of this Note), consistent with a growing importance of the local recombination rate for the efficiency of purifying selection with respect to the effective population size, which is especially important for the endemic island species (as shown by the steeper slopes in Fig. 1E of this note). The stronger effect of recombination for island species is captured by the significant interaction between GC3 and insularity in the linear model : $\pi_N/\pi_S$ ~ GC3+insularity+GC3:insularity ($R^2$ = 0.72, *p model* <2.2x10$^{-16}$, *p GC3* <2.2x10$^{-16}$, *p insularity* = 2.8x10$^{-10}$, *p interaction* = 1.33x10$^{-05}$).

To take into account the potential impact of gBGC on the estimations of $\pi_S$ and $\pi_N$, we used the same approach as in Rousselle et al. 2019 and computed $\pi_N/\pi_S$ based on the Site Frequency Spectrum (SFS) computed for GC-conservative sites only (i.e., A↔T and G↔C mutations), a part of the total variants which is unaffected by gBGC. We also recovered significant differences in $\pi_N/\pi_S$ between island and mainland species at GC-conservative sites

(Figs. 1 & 4 of this Note) (GC-poor: Δmean=0.138 with 95%CI: 0.050-0.225, t=3.284, p=0.004; GC-rich: Δmean=0.050 with 95%CI: 0.018-0.082, t=3.284, p=0.005). Genetic recombination has a higher effect on purifying selection in the island (GC-conservative $\pi_N/\pi_S$, $\Delta mean_{GC\text{-}rich\ vs.\ GC\text{-}poor}$=0.239) than in the mainland species (Δmean=0.151), leading to a weaker difference in $\pi_N/\pi_S$ in highly recombining region. However, this advantage is still insufficient to entirely compensate for the increase in frequency of deleterious alleles due to the increased drift effects in species in small populations, as typically represented by the island Passerida species used in our study

.

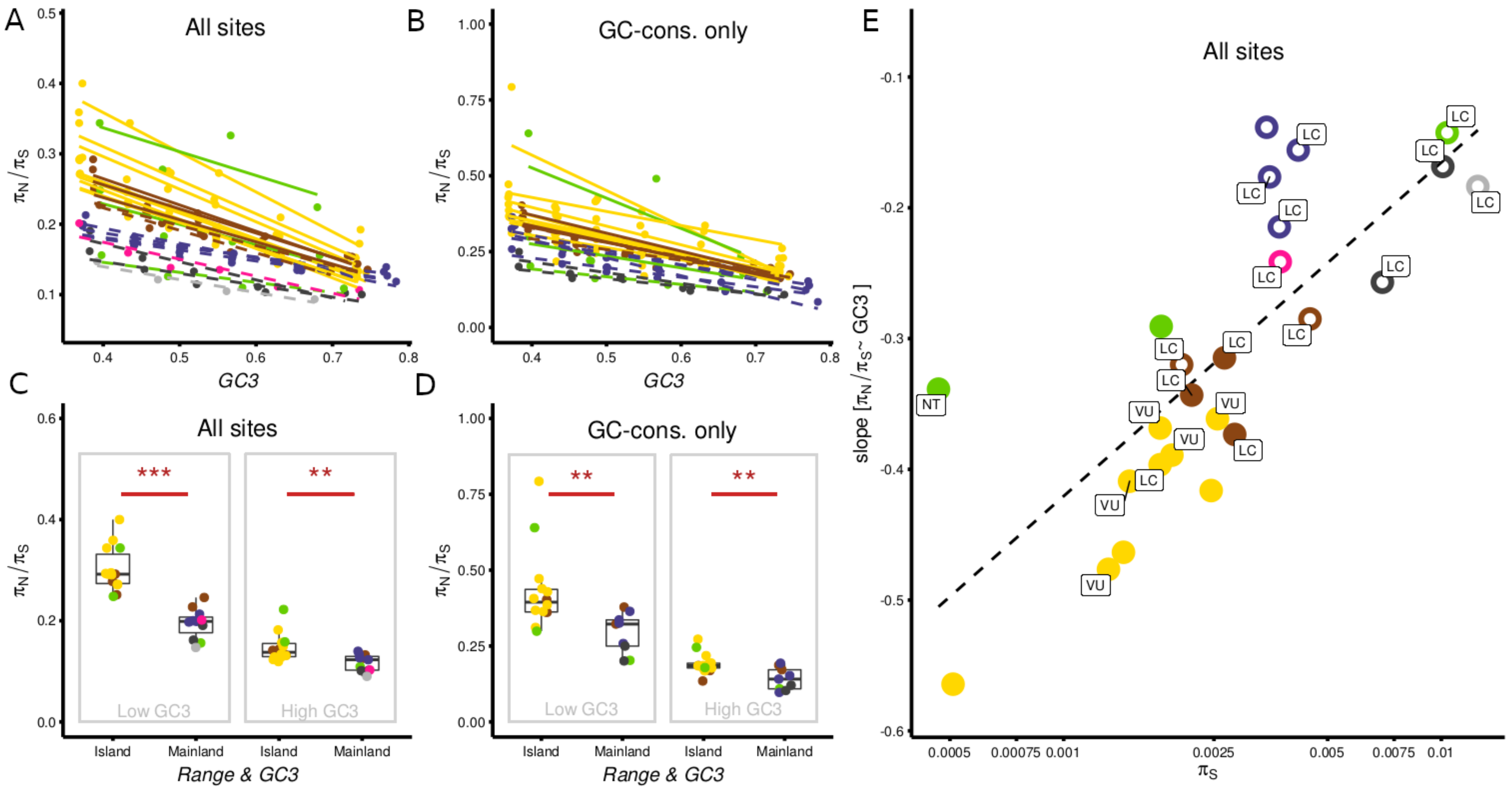


***Fig. 1 of the Note S2: Efficiency of natural selection of insular and mainland Passerida bird species to remove potentially deleterious mutations depending on the local recombination rates.*** *A & B. Linear regressions between* $\pi_N/\pi_S$ *ratios and the third codon position (GC3), a proxy of local recombination rates for all sites (A) and GC-conservative sites only (B). C & D. Observed variation in* $\pi_N/\pi_S$ *ratios at GC-rich and GC-poor genes between island and mainland species for all sites (C) and for GC-conservative sites only (D). E. Slopes of the linear regressions between* $\pi N/\pi S$ *and GC3 (i.e. as shown in the panel A) depending on the synonymous nucleotide diversity. The dotted line indicates the best-fitting linear model.*

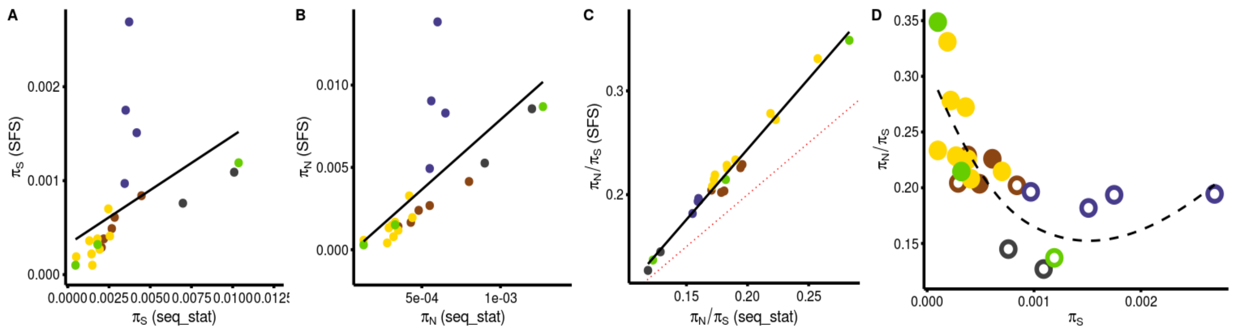


***Fig. 2 of the Note S2: Comparisons between the nucleotide diversity estimates.*** *$\pi_S$ (A), $\pi_N$ (B) and $\pi_N/\pi_S$ (C) as estimated by the diversity-based (meth.1, x-axis) and the SFS-based (meth.2, y-axis) methods. D) Same plot than for the Fig. 1 assuming the SFS-based estimates (dotted black line = loess with span=1.25).*

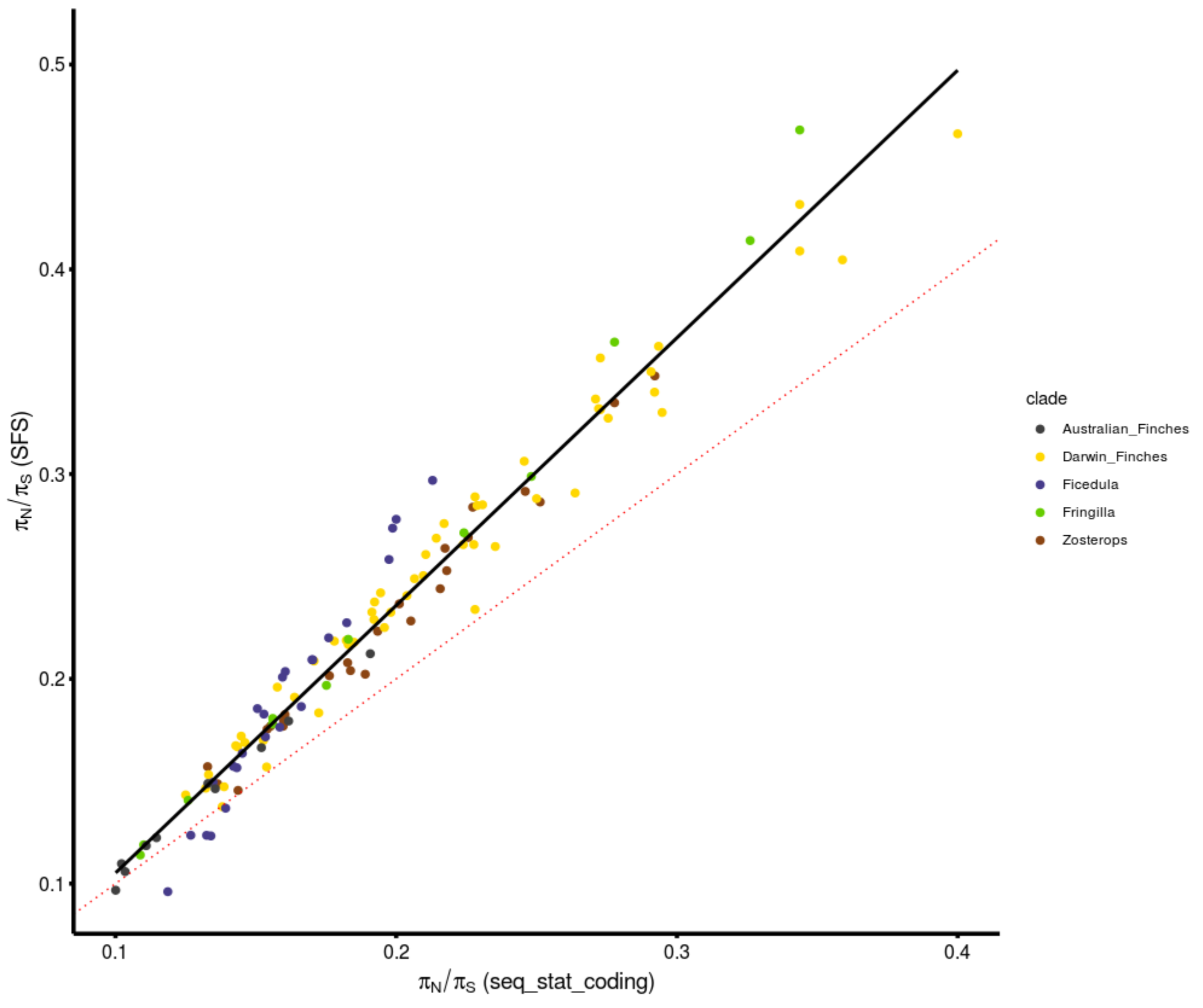


***Fig. 3 of the Note S2: Comparisons of $\pi_N/\pi_S$ estimates using the sequence-based (method 1) and Site-Frequency Spectrum (SFS)-based (method 2) methods based on the same sets of variants*** *(sets are based on the same bins of GC3 than shown in Fig. 2A). Linear regression (black): slope:1.307, $R^2$=0.966, p-value < 2.2 $e^{-16}$. Dotted red line: intercept=0, slope=1*

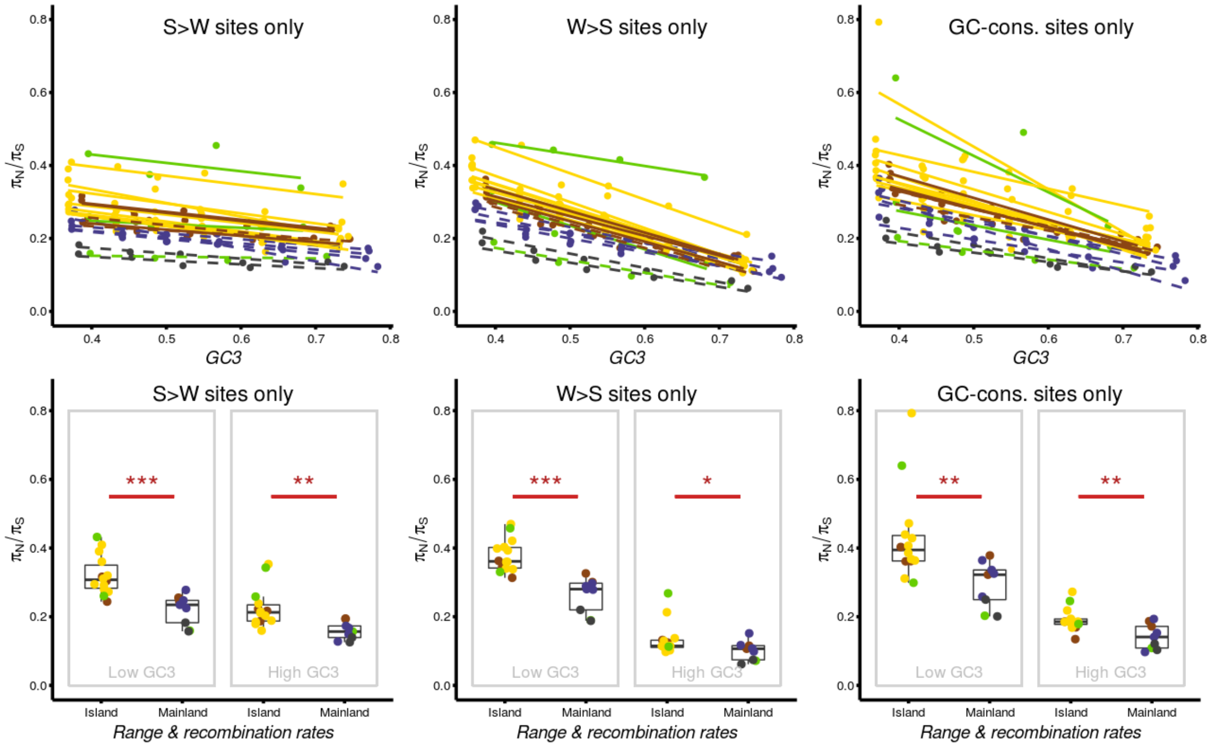


***Fig. 4 of the Note S2**: A,C,E: Linear regressions between $\pi_N/\pi_S$ ratios and the third codon position (GC3), a proxy of local recombination rates for S>W (A), W>S (C) and GC-conservative mutations only (E). B, D & F: Observed variation in $\pi_N/\pi_S$ ratios at GC-rich and GC-poor genes between island and continental species for S>W (B), W>S (D) and GC-conservative mutations only (F).*

### <u>Methods S1: Extended Materials & Methods</u>

##### Darwin's Finch and flycatcher species used in this study

The taxonomy of the Darwin's finches is still debated (Zink & Vázquez-Miranda, 2019). Several species previously described with morphological markers were found to be polyphyletic at the molecular level (Almén et al. 2016; Lamichhaney et al. 2015). Lamichhaney et al. (2015) detected extensive sharing of genetic variation among populations particularly among ground (*Geospiza*) and tree finches (*Camarhynchus*). In order to avoid paraphyly and to limit the level of shared polymorphism among the populations included in our analysis, we excluded populations that are too closely related genetically. In practice, we computed the statistic Da which is the average divergence corrected for within-species diversity: Da = Dxy – (P1+P2)/2 where Dxy is the absolute average divergence between population 1 and 2 and P1 and P2 is the within-species genetic diversity of the population 1 and 2 respectively. Da was computed on a 40-Mb sequence randomly selected across the genome. We computed this statistic between all pairs of Darwin's finch populations and between all pairs of the *Ficedula* flycatcher species. *Ficedula* flycacthers were used as reference because they are considered indisputable species (Ellegren et al. 2012) with reproductive isolation and reduction in fitness in hybrids (Qvarnström et al. 2010).

In flycatchers, observed Da values were close to 0.002 (min Da = 0.0018 between *F. albicollis* and *F. hypoleuca* and max Da = 0.0023 between *F. albicollis* and *F. speculigera*) (Fig 1a of this Note). Based on this observation, we chose a threshold of Da = 0.001 to separate the Darwin's finches populations/species. Based on 61 pairs of populations/species of animals with variable levels of divergence, Roux et al. (2016) have shown that animal populations with a Da above 0.001 have a very low probability of ongoing gene-flow. As expected, Darwin's finches showed much more variation in Da from less than 0.0001 between *G. fuliginosa*_Z (Santa Cruz) and *G. fortis*_M (Daphné Major) to more than 0.0052 between *Certhidea fusca* and *Pinaroloxias inornata*. Applying this threshold, we can delimit 9 lineages (Fig 1b of this Note) from which we selected only one population for further analysis (focusing on the population with the highest sample size) : *Certhidea olivacea* S (Santiago), *C. fusca* E (Española), *C. fusca* L (San Cristobal), *Pinaroloxias crassirostris* Z (Santa Cruz), *P. inornata* C (Coco), *Camarhynchus pallidus* Z (Santa Cruz), *Geospiza difficilis* P (Pinta),

*G. difficilis* W (Wolf, now Geospiza septentrionalis) and *G. conirostris* E (Española). It must be noted that this threshold of Da = 0.001 is probably conservative for birds (Peñalba et al. 2019).

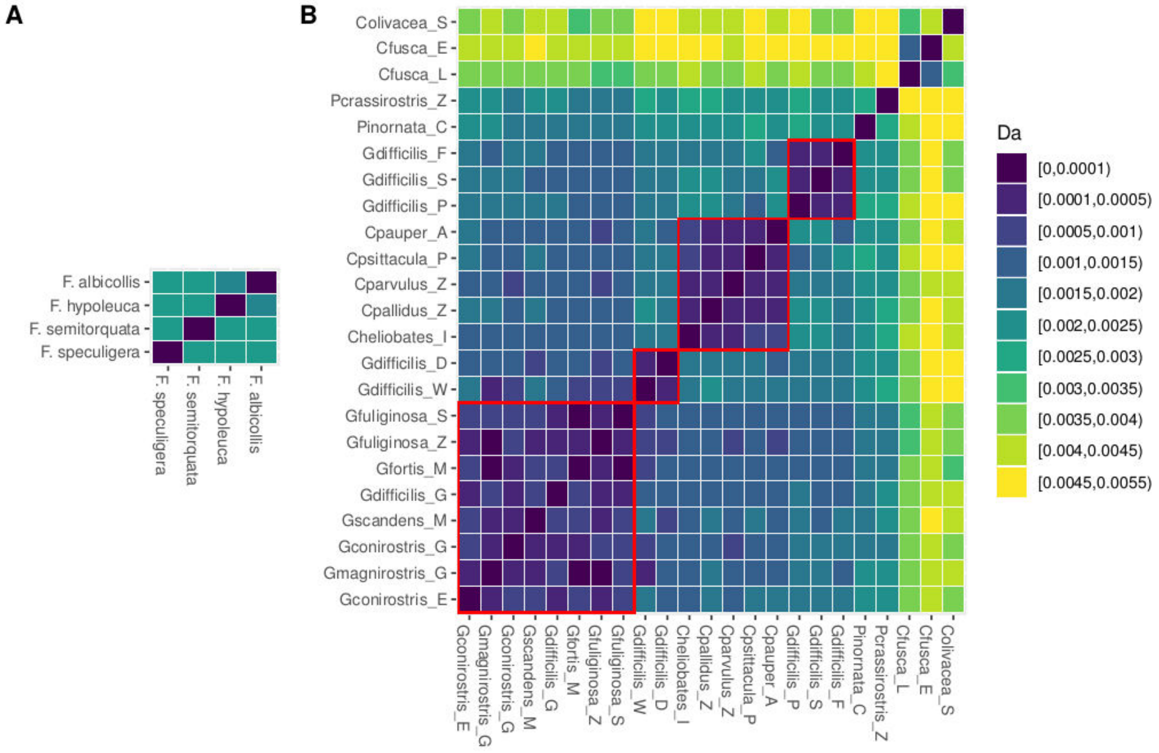


***Fig 1 of the Methods S1**: A) Da values between pairs of Ficedula flycatcher species. B) Da values between pairs of Darwin's finch populations. One population corresponds to a given species on one island. Islands are abbreviated as follows : Genovesa (G), Daphné Major (M), Santa Cruz (Z), Santiago (S), Genovesa (G), Española (E), Daphne Major (M), Pinta (P), Fernandina (F), Santiago (S), Wolf (W), Darwin (D), Genovesa (G), Isabella (I), Pinta (P), Floreana (F), Santiago (S), Española (E), San Cristobal (L), Coco (C). Species names are shown as indicated in Lamichhaney et al. (2015).*

#### Newly generated genome assembly (*Fringilla coelebs*)

DNA from muscle tissues of one *Fringilla* individual captured in Madrid, Spain was sequenced to obtain a draft genome of this species. For this individual, two libraries were prepared, a Chicago library and a Dovetail HiC library (Dovetail Genomics, Scotts Valley, CA), as described previously in Putnam et al. (2016) and Lieberman-Aiden et al. (2009), respectively. Both libraries were sequenced on an Illumina HiSeqX platform to produce over 100 million 2x151 bp reads. A *de novo* assembly was constructed using Meraculous (v. 2.2.2.5 diploid_mode 1, Chapman et al. 2011) with a kmer size of 73 and using 397.3 million paired-end reads (totaling 1,204 Gbp). Reads were trimmed for quality, sequencing adapters, and mate-pair adapters using Trimmomatic (Bolger et al. 2014). The input *de novo* assembly, shotgun reads, Chicago library reads, and Dovetail HiC library reads were used as input data for HiRise, a software pipeline designed specifically for using proximity ligation data to scaffold genome assemblies (Putnam et al. 2016). The HiRise assembly total length was 907.37 Mb obtaining 3,240 scaffolds and a final N50 of 68.85 Mb. An improved version of this genome is now available (Recuerda et al. 2020)

**Empirical estimate of the SNP calling error rate**

One individual of *Zosterops virens* (namely RCHB1917) was sequenced twice at coverage 10x and both replicates went through the entire genotyping pipeline. We estimate the error rate by counting the number of differences between the two replicates. All the differences detected correspond to heterozygous sites. We estimated a rate of $2.9e^{-05}$, which is approximately fifty times lower than the expected genome-wide heterozygosity for this focal species ($1.5e^{-03}$).

**Sequence reconstruction**

For each individual, we then reconstructed fasta sequences from VCF files. First, we discarded all indel variations to ensure an exact match between positions in sequences and coordinates in the reference gff file (i.e. no frameshift mutation allowed). For each scaffold, we performed a base-by-base reconstruction of two sequences per individual by adding either a reference or an alternate allele based on the genotype information available in the vcf file at the position under investigation. Albeit simplistic, this approach is valid here because our study is only based on the nucleotide levels and does not require haplotype phase information. To exclude sites with abnormally low or high coverage, we compute percentiles of the distribution of coverage over the whole genome for each individual and then calibrate a minimum and maximum coverage. All sites exhibiting very low or very high coverage were masked. A minimum threshold value of 3 was set for all species. We then used a home-made python script to extract CDS from these reconstructed sequences based on the gff files.

**Mitochondrial genomes assembly and phylogeny**

For the species with mtDNA genomes available on GenBank, we used the data (accessions: *Ficedula albicollis* : KF293721; Fringilla coelebs : NC_025599; *Fringilla teydea* : KU705740; *Parus major* : NC_026293; *Taeniopygia_guttata* : NC_007897; *Zosterops borbonicus* : MK529728; *Zosterops pallidus* : MK524996. We also add three outgroups : *Corvus_brachyrhynchos* : NC_026461; *Lanius cristatus*: NC_028333; *Menura novaehollandiae* : NC_007883).

For the remaining species, we used MitoFinder (vers. 1.1; https://github.com/RemiAllio/MitoFinder; Allio et al. 2020) to extract all the mitochondrial protein coding genes. One individual per species was randomly selected to use in MitoFinder with default parameters. Alignments were performed gene by gene using macse (vers. 2; Ranwez et al. 2018). Alignments are available at the following URL: https://osf.io/uw6mb/?view_only=a887417cbc91429dac0bbdc1705e2f2b. Then all genes were concatenated to form a supermatrix (missing data = 20% on average, sd = 14%). Next, a phylogenetic analysis was performed using IQTREE (Nguyen et al. 2015) using the "GTR+G4" substitution model and an ultrafast bootstrap option to have a crude approximation of each node support.

### References (SI only):


Allio R, Schomaker-Bastos A, Romiguier J, Prosdocimi F, Nabholz B, Delsuc F. 2020. MitoFinder: efficient automated large-scale extraction of mitogenomic data in target enrichment phylogenomics. *Mol Ecol Resour.* 20: 892– 905.

Almén MS, Lamichhaney S, Berglund J, Grant BR, Grant PR, Webster MT, Andersson L. 2016. Adaptive radiation of Darwin's finches revisited using whole genome sequencing. *BioEssays* 38: 14–20.

Atkinson QD, Gray RD, Drummond AJ. mtDNA Variation Predicts Population Size in Humans and Reveals a Major Southern Asian Chapter in Human Prehistory. *Molecular Biology and Evolution*. 2007;25(2):468-74.

Backström N, Forstmeier W, Schielzeth H, Mellenius H, Nam K, Bolund E, et al. The recombination landscape of the zebra finch Taeniopygia guttata genome. *Genome Res*. 2010/03/31. avr 2010;20(4):485-95.

Baker Z, Schumer M, Haba Y, Bashkirova L, Holland C, Rosenthal GG, et al. Repeated losses of PRDM9-directed recombination despite the conservation of PRDM9 across vertebrates. *eLife*. 2017;6:e24133.

Barker FK, Cibois A, Schikler P, Feinstein J, Cracraft J. Phylogeny and diversification of the largest avian radiation. *Proc Natl Acad Sci U S A.* 2004;101(30):11040.

Baudat F, Buard J, Grey C, Fledel-Alon A, Ober C, Przeworski M, et al. PRDM9 Is a Major Determinant of Meiotic Recombination Hotspots in Humans and Mice. *Science*. 2010;327(5967):836.

Bolger AM, Lohse M, Usadel B. Trimmomatic: a flexible trimmer for Illumina sequence data. *Bioinformatics*. 2014;30(15):2114-20.

Bourgeois YXC, Delahaie B, Gautier M, Lhuillier E, Malé P-JG, Bertrand JAM, et al. A novel locus on chromosome 1 underlies the evolution of a melanic plumage polymorphism in a wild songbird. *Royal Society Open Science*. 4(2):160805.

Burri R, Nater A, Kawakami T, Mugal CF, Olason PI, Smeds L, Suh A, Dutoit L, Bureš S, Garamszegi LZ, et al. 2015. Linked selection and recombination rate variation drive the evolution of the genomic landscape of differentiation across the speciation continuum of Ficedula flycatchers. *Genome Research*.

Charlesworth B. Effective population size and patterns of molecular evolution and variation. *Nature Reviews Genetics.* 2009;10(3):195-205.

Chapman JA, Ho I, Sunkara S, Luo S, Schroth GP, Rokhsar DS. Meraculous: De Novo Genome Assembly with Short Paired-End Reads. *PLOS ONE*. 2011;6(8):e23501.

Chen J, Glémin S, Lascoux M. Genetic Diversity and the Efficacy of Purifying Selection across Plant and Animal Species. *Molecular Biology and Evolution*. 2017;34(6):1417-28.

Corcoran P, Gossmann TI, Barton HJ, Great Tit HapMap Consortium, Slate J, Zeng K. Determinants of the Efficacy of Natural Selection on Coding and Noncoding Variability in Two Passerine Species. *Genome Biol Evol.* 2017;9(11):2987-3007.

Cornetti L, Valente LM, Dunning LT, Quan X, Black RA, Hébert O, et al. The Genome of the "Great Speciator" Provides Insights into Bird Diversification*. Genome Biology and Evolution.* 2015;7(9):2680-91.

Delmore K, Illera J.C, Pérez-Tris J, Segelbacher G, Lugo Ramos JS, Durieux G, Ishigohoka J, Liedvogel M. The evolutionary history and genomics of European blackcap migration. *eLife* 2020; 9 e54462.

Díez-del-Molino D, Sánchez-Barreiro F, Barnes I, Gilbert MTP, Dalén L. Quantifying Temporal Genomic Erosion in Endangered Species*. Trends in Ecology & Evolution*. 2018;33(3):176-85.

Duret L, Galtier N. Biased Gene Conversion and the Evolution of Mammalian Genomic Landscapes. *Annu Rev Genom Hum Genet*. 2009;10(1):285-311.

Ellegren H. Evolutionary stasis: the stable chromosomes of birds. *Trends in Ecology & Evolution*. 2010;25(5):283-91.

Ellegren H, Smeds L, Burri R, Olason PI, Backström N, Kawakami T, et al. The genomic landscape of species divergence in Ficedula flycatchers. *Nature.* 2012;491:756.

Eyre-Walker A. Changing effective population size and the McDonald-Kreitman test. *Genetics.* 2002;162(4):2017-24.

Eyre-Walker A, Keightley PD. Estimating the Rate of Adaptive Molecular Evolution in the Presence of Slightly Deleterious Mutations and Population Size Change. *Molecular Biology and Evolution*. 2009;26(9):2097-108.

Figuet E, Nabholz B, Bonneau M, Mas Carrio E, Nadachowska-Brzyska K, Ellegren H, et al. Life History Traits, Protein Evolution, and the Nearly Neutral Theory in Amniotes. *Molecular Biology and Evolution.* 2016;33(6):1517-27.

Frankham R, Ballou JD, Briscoe DA. Introduction to Conservation Genetics. Cambridge: Cambridge University Press; 2002.

Galtier N. Adaptive Protein Evolution in Animals and the Effective Population Size Hypothesis. *PLOS Genetics.* 2016;12(1):e1005774.

Gossmann TI, Keightley PD, Eyre-Walker A. The Effect of Variation in the Effective Population Size on the Rate of Adaptive Molecular Evolution in Eukaryotes. *Genome Biology and Evolution.* 2012;4(5):658-67.

Gossmann TI, Song B-H, Windsor AJ, Mitchell-Olds T, Dixon CJ, Kapralov MV, et al. Genome Wide Analyses Reveal Little Evidence for Adaptive Evolution in Many Plant Species. *Molecular Biology and Evolution.* 2010;27(8):1822-32.

Hillier LW, Miller W, Birney E, Warren W, Hardison RC, Ponting CP, et al. Sequence and comparative analysis of the chicken genome provide unique perspectives on vertebrate evolution. *Nature.* 2004;432(7018):695-716.

James JE, Lanfear R, Eyre-Walker A. Molecular Evolutionary Consequences of Island Colonization. *Genome Biology and Evolution.* 2016;8(6):1876-88.

James J, Eyre-Walker A. Mitochondrial DNA Sequence Diversity in Mammals: a Correlation Between the Effective and Census Population Sizes. *bioRxiv.* 2020;2020.02.28.969592.

Jensen JD, Payseur BA, Stephan W, Aquadro CF, Lynch M, Charlesworth D, et al. The importance of the Neutral Theory in 1968 and 50 years on: A response to Kern and Hahn 2018. *Evolution.* 2019;73(1):111-4.

Johnson TH, Stattersfield AJ. A global review of island endemic birds. *Ibis*. 1990;132(2):167-80.

Keightley PD, Eyre-Walker A. Joint inference of the distribution of fitness effects of deleterious mutations and population demography based on nucleotide polymorphism frequencies. *Genetics.* 2007;177(4):2251-61.

Keightley PD, Eyre-Walker A. What can we learn about the distribution of fitness effects of new mutations from DNA sequence data? *Philosophical Transactions of the Royal Society B: Biological Sciences.* 2010;365(1544):1187-93.

Kern AD, Hahn MW. The Neutral Theory in Light of Natural Selection. *Molecular Biology and Evolution.* 2018;35(6):1366-71.

Kimura M. The Neutral Theory of Molecular Evolution. Cambridge: Cambridge University Press; 1983.

Kimura M, Crow JF. The number of alleles that can be maintained in a finite population. *Genetics*. 1964;49(4):725.

Kutschera VE, Poelstra JW, Botero-Castro F, Dussex N, Gemmell NJ, Hunt GR, et al. Purifying Selection in Corvids Is Less Efficient on Islands. *Molecular Biology and Evolution*. 2019;37(2):469-74.

Laine VN, Gossmann TI, Schachtschneider KM, Garroway CJ, Madsen O, Verhoeven KJF, et al. Evolutionary signals of selection on cognition from the great tit genome and methylome. *Nature Communications.* 2016;7(1):10474.

Lamichhaney S, Berglund J, Almén MS, Maqbool K, Grabherr M, Martinez-Barrio A, et al. Evolution of Darwin's finches and their beaks revealed by genome sequencing. *Nature.* 2015;518(7539):371-5.

Lanfear R, Kokko H, Eyre-Walker A. Population size and the rate of evolution. *Trends in Ecology & Evolution.* 2014;29(1):33-41.

Leroy T, Anselmetti Y, Tilak M-K, Bérard S, Csukonyi L, Gabrielli M, et al. A bird's white-eye view on neosex chromosome evolution. *BioRxiv.* 2019;505610, ver. 4 peer-reviewed and recommended by Peer Community in Evolutionary Biology.

Li H. Aligning sequence reads, clone sequences and assembly contigs with BWA-MEM. *arXiv*. 2013;(1303.3997v1).

Lieberman-Aiden E, van Berkum NL, Williams L, Imakaev M, Ragoczy T, Telling A, et al. Comprehensive Mapping of Long-Range Interactions Reveals Folding Principles of the Human Genome. *Science.* 2009;326(5950):289.

Loire E, Chiari Y, Bernard A, Cahais V, Romiguier J, Nabholz B, et al. Population genomics of the endangered giant Galápagos tortoise. *Genome Biology.* 2013;14(12):R136.

Lundberg M, Liedvogel M, Larson K, Sigeman H, Grahn M, Wright A, et al. Genetic differences between willow warbler migratory phenotypes are few and cluster in large haplotype blocks. *Evolution Letters.* 2017;1(3):155-68.

McDonald JH, Kreitman M. Adaptive protein evolution at the Adh locus in Drosophila. *Nature.* 1991;351(6328):652-4.

McKenna A, Hanna M, Banks E, Sivachenko A, Cibulskis K, Kernytsky A, et al. The Genome Analysis Toolkit: a MapReduce framework for analyzing next-generation DNA sequencing data. *Genome Res.* 2010;20(9):1297-303.

Moutinho AF, Bataillon T, Dutheil JY. Variation of the adaptive substitution rate between species and within genomes. *Evolutionary Ecology*. 2019, in press

Nam K, Munch K, Mailund T, Nater A, Greminger MP, Krützen M, et al. Evidence that the rate of strong selective sweeps increases with population size in the great apes. *Proc Natl Acad Sci USA*. 2017;114(7):1613.

Nguyen L-T, Schmidt HA, von Haeseler A, Minh BQ. 2015. IQ-TREE: A Fast and Effective Stochastic Algorithm for Estimating Maximum-Likelihood Phylogenies. *Molecular Biology and Evolution* 32: 268–274.

Ohta T. Slightly Deleterious Mutant Substitutions in Evolution. *Nature.* 1973;246(5428):96-8.

Ohta T. The Nearly Neutral Theory of Molecular Evolution. *Annu Rev Ecol Syst*. 1992;23(1):263-86.

Oliveros CH, Field DJ, Ksepka DT, Barker FK, Aleixo A, Andersen MJ, et al. Earth history and the passerine superradiation. *Proc Natl Acad Sci USA.* 2019;116(16):7916.

Parvanov ED, Petkov PM, Paigen K. Prdm9 Controls Activation of Mammalian Recombination Hotspots. *Science.* 2010;327(5967):835.

Peart CR, Tusso S, Pophaly SD, Botero-Castro F, Wu C-C, Aurioles-Gamboa D, Baird AB, Bickham JW, Forcada J, Galimberti F, et al. 2020. Determinants of genetic variation across eco-evolutionary scales in pinnipeds. *Nature Ecology & Evolution* 4: 1095–1104.

Peñalba JV, Joseph L, Moritz C. 2019. Current geography masks dynamic history of gene flow during speciation in northern Australian birds*. Molecular Ecology* 28: 630–643.

Popadin K, Polishchuk LV, Mamirova L, Knorre D, Gunbin K. Accumulation of slightly deleterious mutations in mitochondrial protein-coding genes of large versus small mammals. *Proc Natl Acad Sci USA.* 2007;104(33):13390.

Prum RO, Berv JS, Dornburg A, Field DJ, Townsend JP, Lemmon EM, et al. A comprehensive phylogeny of birds (Aves) using targeted next-generation DNA sequencing. *Nature.* 2015;526(7574):569-73.

Putnam NH, O'Connell BL, Stites JC, Rice BJ, Blanchette M, Calef R, et al. Chromosome-scale shotgun assembly using an in vitro method for long-range linkage. *Genome Research*. 2016;26(3):342-50.

Qvarnström A, Rice AM, Ellegren H. 2010. Speciation in *Ficedula* flycatchers. *Philosophical Transactions of the Royal Society B: Biological Sciences* 365: 1841–1852.

Ranwez V, Douzery EJP, Cambon C, Chantret N, Delsuc F. MACSE v2: Toolkit for the Alignment of Coding Sequences Accounting for Frameshifts and Stop Codons. *Molecular Biology and Evolution.* 2018;35(10):2582-4.

Recuerda, M., Vizueta, J., Cuevas-Caballé, C., Blanco, G., Rozas, J., and Milá, B. Chromosome-level genome assembly of the common chaffinch (Aves: *Fringilla coelebs*): a valuable resource for evolutionary biology. bioRxiv, 2020; 2020.11.30.404061.

Ricketts TH, Dinerstein E, Boucher T, Brooks TM, Butchart SHM, Hoffmann M, et al. Pinpointing and preventing imminent extinctions. *Proc Natl Acad Sci U S A.* 2005;102(51):18497.

Robinson JA, Brown C, Kim BY, Lohmueller KE, Wayne RK. Purging of Strongly Deleterious Mutations Explains Long-Term Persistence and Absence of Inbreeding Depression in Island Foxes. *Current Biology.* 2018;28(21):3487-3494.e4.

Robinson JA, Ortega-Del Vecchyo D, Fan Z, Kim BY, vonHoldt BM, Marsden CD, et al. Genomic Flatlining in the Endangered Island Fox. *Current Biology.* 2016;26(9):1183-9.

Rogers RL, Slatkin M. Excess of genomic defects in a woolly mammoth on Wrangel island. *PLOS Genetics.* 2017;13(3):e1006601.

Romiguier J, Gayral P, Ballenghien M, Bernard A, Cahais V, Chenuil A, et al. Comparative population genomics in animals uncovers the determinants of genetic diversity. *Nature.* 2014;515(7526):261-3.

Rousselle M, Laverré A, Figuet E, Nabholz B, Galtier N. Influence of Recombination and GC-biased Gene Conversion on the Adaptive and Nonadaptive Substitution Rate in Mammals versus Birds. *Molecular Biology and Evolution.* 2019;36(3):458-71.

Rousselle M, Mollion M, Nabholz B, Bataillon T, Galtier N. Overestimation of the adaptive substitution rate in fluctuating populations. *Biology Letters.* 2018;14(5):20180055.

Rousselle M, Simion P, Tilak M-K, Figuet E, Nabholz B, Galtier N. Is adaptation limited by mutation? A timescale dependent effect of genetic diversity on the adaptive substitution rate in animals. *Plos Genetics*. 2020,16(4):e1008668
Roux C, Fraïsse C, Romiguier J, Anciaux Y, Galtier N, Bierne N. 2016. Shedding Light on the Grey Zone of Speciation along a Continuum of Genomic Divergence. *PLOS Biology* 14: e2000234.

Schiffels S, Durbin R. Inferring human population size and separation history from multiple genome sequences. *Nature Genetics.* 2014;46(8):919-25.

Singhal S, Leffler EM, Sannareddy K, Turner I, Venn O, Hooper DM, et al. Stable recombination hotspots in birds*. Science.* 2015;350(6263):928.

Steadman DW. Prehistoric Extinctions of Pacific Island Birds: Biodiversity Meets Zooarchaeology. *Science.* 1995;267(5201):1123.

Tataru P, Mollion M, Glémin S, Bataillon T. Inference of Distribution of Fitness Effects and Proportion of Adaptive Substitutions from Polymorphism Data. *Genetics.* 2017;207(3):1103-19.

Zhang G, Li C, Li Q, Li B, Larkin DM, Lee C, Storz JF, Antunes A, Greenwold MJ, Meredith RW, et al. 2014. Comparative genomics reveals insights into avian genome evolution and adaptation. *Science* 346: 1311.
